# Universal annotation of the human genome through integration of over a thousand epigenomic datasets

**DOI:** 10.1101/2020.11.17.387134

**Authors:** Ha Vu, Jason Ernst

## Abstract

**Background:** Genome-wide maps of chromatin marks such as histone modifications and open chromatin sites provide valuable information for annotating the non-coding genome, including identifying regulatory elements. Computational approaches such as ChromHMM have been applied to discover and annotate chromatin states defined by combinatorial and spatial patterns of chromatin marks within the same cell type. An alternative ‘stacked modeling’ approach was previously suggested, where chromatin states are defined jointly from datasets of multiple cell types to produce a single universal genome annotation based on all datasets. Despite its potential benefits for applications that are not specific to one cell type, such an approach was previously applied only for small-scale specialized purposes. Large-scale applications of stacked modeling have previously posed scalability challenges.

**Results:** Using a version of ChromHMM enhanced for large-scale applications, we applied the stacked modeling approach to produce a universal chromatin state annotation of the human genome using over 1000 datasets from more than 100 cell types, with the learned model denoted as the full-stack model. The full-stack model states show distinct enrichments for external genomic annotations, which we used in characterizing each state. Compared to per-cell-type annotations, the full-stack annotations directly differentiate constitutive from cell type specific activity and is more predictive of locations of external genomic annotations.

**Conclusions:** The full-stack ChromHMM model provides a universal chromatin state annotation of the genome and a unified global view of over 1000 datasets. We expect this to be a useful resource that complements existing per-cell-type annotations for studying the non-coding human genome.

## Introduction

Genome-wide maps of histone modifications, histone variants and open chromatin provide valuable information for annotating the non-coding genome features, including various types of regulatory elements [1–5]. These maps -- produced by assays such as chromatin immunoprecipitation followed by high-throughput sequencing to map histone modifications or DNase-seq to map open chromatin-- can facilitate our understanding of regulatory elements and genetic variants that are associated with disease [6–12]. Efforts by large scale consortia as well as many individual labs have resulted in these maps for many different human cell and tissue types for multiple different chromatin marks [1,8,13–20].

The availability of maps for multiple different chromatin marks in the same cell type motivated the development of methods such as ChromHMM and Segway that learn ‘chromatin states’ based on the combinatorial and spatial patterns of marks in such data [21–23] These methods then annotate genomes in a per-cell-type manner based on the learned chromatin states. They have been applied to annotate more than a hundred diverse cell and tissue types [3,16,24]. Previously, large collections of per-cell-type chromatin state annotations have been generated using either (1) independent models that learn a different set of states in each cell type or (2) a single model that is learned across all cell types, resulting in a common set of states across cell types, yet generating per-cell-type annotations (in some cases per-tissue-type annotations are generated, but we will use the terms cell-type and tissue interchangeably for ease of presentation). This latter approach is referred to as a ‘concatenated’ approach (**Supp. Fig. 1**) [22, 25]. Variants of the concatenated approach attempt to use information from related cell types to reduce the effect of noise, but still output per-cell-type annotations [26, 27]. These models that produce per-cell-type annotations tend to be most appropriate in studies where researchers are interested in studying individual cell types.

A complementary approach to applying ChromHMM to data across multiple different cell types referred to as the ‘stacked’ modeling approach was also previously suggested (**Supp. Fig. 1)** [22, 25]. Instead of learning per-cell-type annotations based on a limited number of datasets per-cell-type, the stacked modeling approach learns a single universal genome annotation based on the combinatorial and spatial patterns in datasets from multiple marks across multiple cell types. This approach differs from the concatenated and independent modeling approaches as those approaches only identify combinatorial and spatial patterns present among datasets within one cell type.

Such a universal annotation from stacked modeling provides potential complementary benefits to existing per-cell-type chromatin state annotations. First, since the model can learn patterns from signals from the same assay across cell types, a stacked model may help differentiate regions with constitutive chromatin activities from those with cell-type-specific activities. Previously, subsets of the genome assigned to individual chromatin states from ‘concatenated’ per-cell-type annotations were post-hoc clustered to analyze chromatin dynamics across cell and tissue types [3, 16]. However, such an approach does not provide a view of the dynamics of all the data at once, which the stacked modeling provides. Second, the stacked modeling approach bypasses the need to pick a specific cell or tissue type when analyzing a single partitioning and annotation of the genome. Focusing on a single cell or tissue type may not be desirable for many analyses involving other data that are not inherently cell-type-specific, such as those involving conserved DNA sequence or genetic variants. For example, when studying the relationship between chromatin states and evolutionarily conserved sequences, if one uses per-cell-type chromatin state annotations from one cell type, many bases will lack an informative chromatin state assignment (e.g. many bases are in a quiescent state), while subsets of those bases will have a more informative annotation in other cell types. Third, if one tries to analyze per-cell-type annotations across cell types, one would need a post-hoc method to reason about an exponentially large number of possible combinations of chromatin states across cell types (if each of *K* cell types has *M* states, there are *M^K^* possible combinations of states for a genomic position) many of which would likely lack biologically meaningful distinctions. In contrast, for the stacked model, there will be a single annotation per position out of a possibly much smaller fixed number of states (compared to *M^K^*). These states are directly informative of cross-cell type activity, though the state definitions can be more complex. Finally, annotations by the stacked modeling leverages a larger set of data for annotation, and thus has the potential to be able to identify genomic elements with greater sensitivity and specificity.

Despite the potential complementary advantages of the ‘stacked’ modeling approach, it has only been applied on a limited scale to combine data from a small number of cell types for highly specialized purposes [28, 29]. No large-scale application of the stacked modeling approach to many diverse cell and tissue types has been previously demonstrated. This may have in part been due to large-scale applications of stacked modeling raising scalability challenges not present in modeling approaches for per-cell-type annotations.

Here, we present a large-scale application of the stacked modeling approach with more than a thousand human epigenomic datasets as input, using a version of ChromHMM for which we enhanced the scalability. We conduct various enrichment analyses on the states resulting from the stacked modeling and give biological interpretations to them. We show that compared to the per-cell-type annotations, the stacked model’s annotation shows greater correspondence to various external genomic annotations not used in the model learning. We analyze the states in terms of enrichment with different types of variation, and highlight specific states of the stacked model that are enriched with phenotypically associated genetic variants, cancer-associated somatic mutations, and structural variants. We expect the stacked model annotations and detailed characterization of the states that we provide will be a valuable resource for studying the epigenome and non-coding genome, complementing existing per-cell-type annotations.

## Results

### Annotating the human genome into universal chromatin states

We used the stacked modeling approach of ChromHMM to produce a universal chromatin state annotation of the genome based on data from over 100 cell and tissue types from the Roadmap Epigenomics and ENCODE projects (**Fig. 1**) [14, 16]. In total we applied ChromHMM to 1032 datasets for 30 histone modifications, a histone variant (H2A.Z), and DNase I hypersensitivity (**Supp. Fig. 2**). The set of cell and tissue types were the same as those for which per-cell-type annotations were previously generated by applying the ‘concatenated’ modeling approach of ChromHMM [16,22,25]. We note that not all chromatin marks were profiled in all cell or tissue types, but the stacked modelling can still be applied directly.

**Figure 1:**
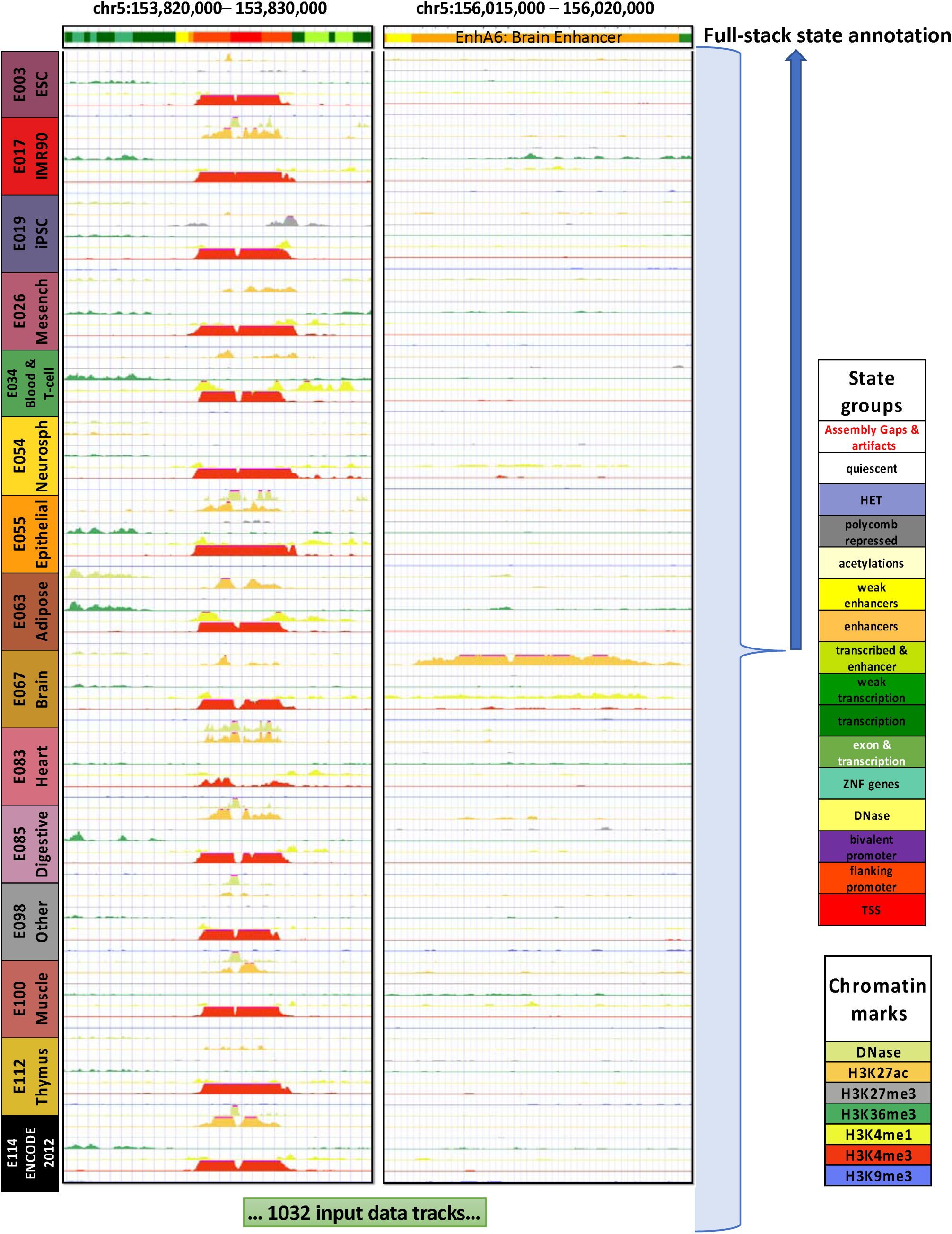
Illustration of full-stack modeling annotations. The figure illustrates the full-stack modeling at two loci. The top track shows chromatin state annotations from the full-stack modeling colored based on the legend at right. Below it are signal tracks for a subset of the 1032 input datasets. Data from seven (DNase I hypersensitivity, H3K27me3, H3K36me3, H3K4me1, H3K4me2, H3K4me3, and H3K9me3) of the 32 chromatin marks are shown, colored based on the legend at right. These data are from 15 of the 127 reference epigenomes each representing different cell and tissue groups. The loci on left highlights a genomic region for which a portion is annotated as constitutive promoter states (TSS1-2). The loci on the right panel highlights a region for which a portion is annotated as a brain enhancer state (EnhA6), which has high signals of H3K27ac in reference epigenomes of the group Brain. Per-cell-type concatenated model annotations for these loci from these and additional reference epigenomes can be found in **Supp. Fig. 24.**

We focused our analysis on a model with 100 states (**Methods**). The number of states is larger than typically used for models that generate per-cell-type annotations, which reflects the greater information available when defining states based on data from many cell types. This number of states was large enough to be able to capture some relatively cell-type-specific regulatory activity, while being small enough to give distinct biological interpretations to each state (**Supp. Fig. 3**). We denote the model’s output chromatin state annotation the ‘full-stack’ genome annotation.

### Major groups of full-stack states

We characterized each state of the model by analyzing the model parameters (emission probabilities and transition probabilities) and state enrichments for other genome annotations (**Fig.2, 3A, Supp. Fig. 4-7**). The other genomic annotations include previous per-cell-type chromatin state annotations (**Supp. Fig. 8-9**), cell-type-specific gene expression data (**Supp. Fig. 10-11**), and various independent existing genomic annotations (**Fig. 3A**). These independent genomic annotations included annotated gene features, evolutionary constrained elements, and assembly gaps, among others (**Methods**).

**Figure 2:**
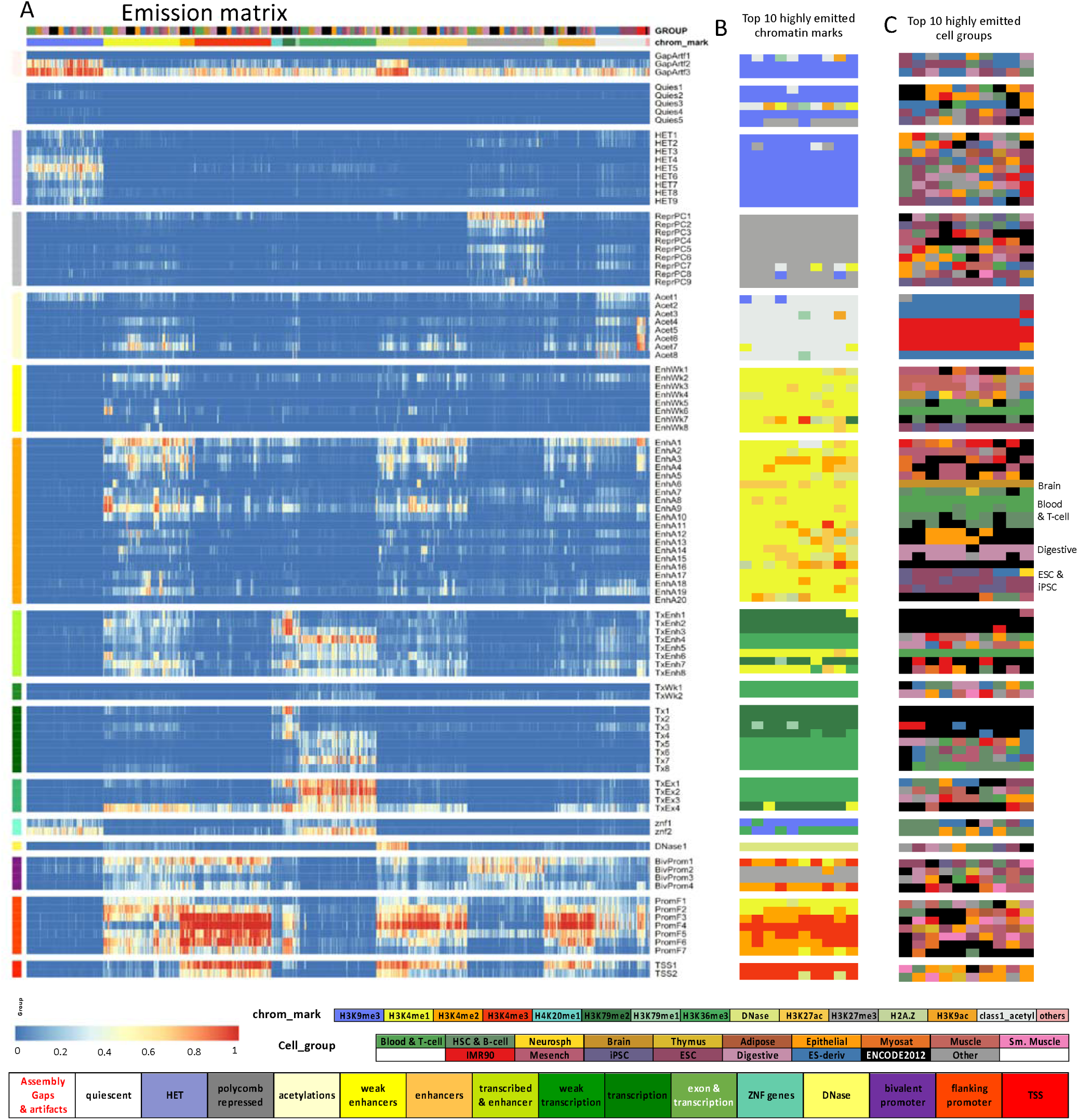
Full-stack state emission parameters. **(A)** Each of the 100 rows in the heatmap corresponds to a full-stack state. Each of the 1032 columns corresponds to one experiment. For each state and each experiment, the heatmap gives the probability within the state of observing a binary present call for the experiment’s signal. Above the heatmap there are two rows, one indicating the cell or tissue type of the experiment and the other indicating the chromatin mark. The corresponding color legends are shown towards the bottom. The states are displayed in 16 groups with white space between each group. The states were grouped based on biological interpretations indicated by the color legend at the bottom. Full characterization of states is available in **Supplementary Data**. The model’s transition parameters between states can be found in **Supp. Fig. 6.** Columns are ordered such that experiments profiling the same chromatin marks are next to each other. **(B)** Each row corresponds to a full-stack state as ordered in (A). The columns correspond to the top 10 experiments with the highest emission value for each state, in order of decreasing ranks, colored by their associated chromatin marks as in (A). **(C)** Similar to **(B)**, but experiments are colored by the associated cell or tissue type group. We noted on the right the cell or tissue groups of some per-cell-type enhancer states.

**Figure 3:**
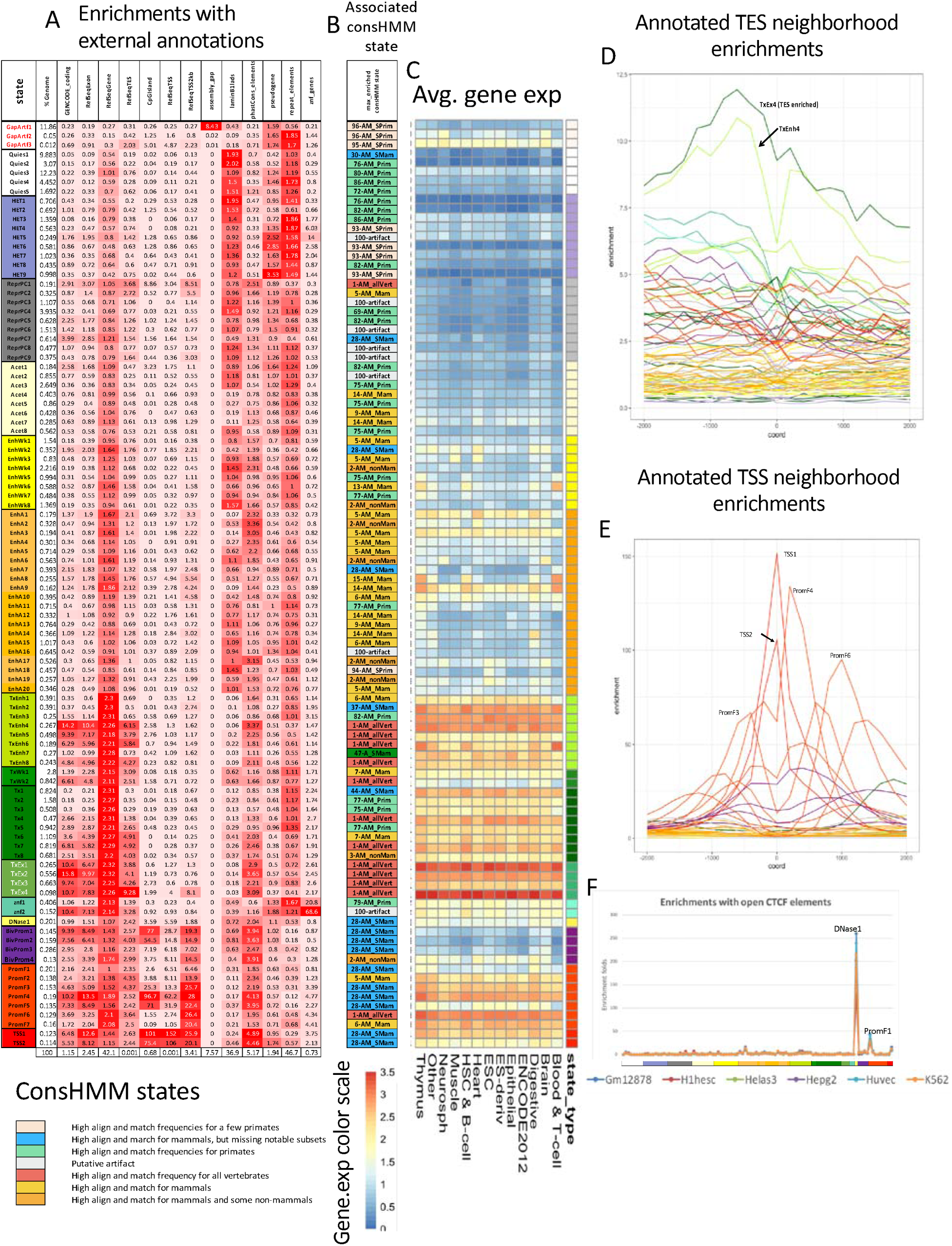
Full-stack states enrichments for external genomic annotations. **(A)** Fold enrichments of full-stack states with external genome annotations (**Methods**). Each row corresponds to a state and each column corresponds to one external genomic annotation: CpG Islands, Exons, coding sequences, gene bodies (exons and introns), transcription end sites (TES), transcription start sites (TSS), TSS and 2kb surrounding regions, lamina associated domains (laminB1lads), assembly gaps, annotated ZNF genes, repeat elements and PhastCons constrained element (**Methods**). The last row shows the percentage of the genome that each external genome annotation covers. The heatmap colors are column-normalized, i.e. within each column, the color of the cells are such that highest values are colored red and lowest values are colored white. **(B)** Each row indicates the ConsHMM state [46] that has the highest enrichment fold in each full-stack state as ordered in **(A).** Legends of the ConsHMM state groups indicated with different colors are shown below the heatmap in **(A)**, and descriptions of select ConsHMM states curated from <u>(Arneson & Ernst, 2019)</u> are available in **Supplementary Data 6.** **(C)** Average weighted expression of genes that overlap each full-stack state in different groups of cells (**Methods**). Each column corresponds to a cell group indicated at the bottom. Each row corresponds to a state, as ordered in **(A).** **(D-E)** Positional enrichments of full-stack states relative to annotated **(D)** transcription end sites (TES) and **(E)** transcription start sites (TSS). Positive coordinate values represent the number of bases downstream in the 5’ to 3’ direction of transcription, while negative values represent the number of bases upstream. Each line shows the positional enrichments in a state. Lines are colored as indicated in **(A**). **(F)** Enrichments of full-stacks states with per-cell-type chromatin states associated with CTCF and open chromatin, but limited histone modifications in six cell types [31] (**Methods**). The six cell types are indicated along the bottom of the figure. States are displayed horizontally in the same order as **(A)**. The DNase1 state showed the strongest enrichment for the per-cell-type chromatin states associated with CTCF and open chromatin in all six cell types.

These analyses led us to group the 100 full-stack states into 16 groups (**Fig. 2A**). One group includes states associated with assembly gaps (GapArtf1) and alignment artifacts (GapArtf2-3). Some other groups are associated with repressive or inactive states, including quiescent states (Quies1-5) (low emissions of all experiments, except possibly weak signals in H3K9me3), heterochromatin states associated with H3K9me3 (HET1-9), and polycomb repressed states associated with H3K27me3 (ReprPC1-9). There is an acetylations group marked primarily by high emission of various acetylation marks profiled only in IMR90 or ESC derived cells, while having weaker signals for enhancer or promoter associated marks such as H3K4me1/2/3, H3K27ac, and H3K9ac (Acet1-8). We also identified active and weak candidate enhancers groups (EnhW1-8 and EnhA1-20, respectively) associated with H3K4me1, DNase, H2A.Z, and/or H3K27ac. Four groups are associated with transcriptional activities, including a group of transcribed enhancers (TxEnh1-8), two groups of weak or strong transcription (TxWk1-2, Tx1-8, respectively), and one group associated with exons and transcription (TxEx1-4). These transcriptional activities groups are associated with at least one of these marks: H3K36me3, H3K79me1, H3K79me2, and H4K20me1. Another group consists of two zinc finger (ZNF) gene states associated with H3K36me3 and H3K9me3 (ZNF1-2). A DNase group consists of one state (DNase1) with strong emission of *only* DNase I hypersensitivity in all profiled cell types. Three groups are associated with promoter activities, marked by emission of some promoter marks such as H3K4me3, H3K4me2, and H3K9ac. One promoter group was of bivalent states associated with promoter marks and H3K27me3 (BivProm1-4). The other two promoter groups were flanking promoter states (PromF1-7) and transcription start sites (TSS) states (TSS1-2) where the flanking promoter states also show emission of H3K4me1.

Enrichments for external annotations supported these state groupings (**Fig. 3A**), as well as further distinctions or characterizations among states within each group. For example, the state GapArt1 had ∼8 fold enrichment for assembly gaps and contained 99.99% of all assembly gaps in hg19 (**Fig. 3A**). In previous concatenated models based on the Roadmap Epigenomics data [16], no specific state was associated with assembly gaps likely due to the limited number of input chromatin mark signals compared to the number of states, leading to assembly gaps being incorporated in the general quiescent state. The states in the zinc finger gene group, ZNF1-2, had 20.8 and 68.6 fold enrichment for zinc finger named genes, respectively (**Fig. 3A**). States in the Acet group had a lower average expression of proximal genes compared to states in the enhancer and promoter groups (**Fig. 3C**) while higher compared to ReprPC and HET groups. States in the transcription groups (TxEnh1-8, TxWk1-2, Tx1-8, TxEx1-4) were all at least 2.1 fold enriched for annotated gene bodies; these gene bodies covered 88.8–97.5% of the states. These states are associated with higher expression of genes across different cell types, particularly when downstream of their TSS (**Fig 3A, C, Supp. Fig. 10-11**). Distinctions were seen among these states, for example, in terms of their positional enrichments relative to TES (**Fig. 3A, D, Supp. Fig. 12**). States in the flanking promoter group (PromF1-7) showed 6.5-28 fold enrichment for being within 2kb of annotated TSS, and genes whose TSS regions overlapped these states had higher average gene expression across different cell types (**Fig. 3A, C, Supp. Fig. 10-11**). These states differed among each other in their enrichments with upstream or downstream regions of the TSS (**Fig. 3E, Supp. Fig. 12**). The states in the transcription start site group (TSS1-2) had enrichment values that peaked at the TSS (>= 100 fold enrichment) (**Fig. 3A, E**). States in promoter-associated groups (TSS, PromF, BivProm), along with those in other groups, show various enriched gene-ontology (GO) terms based on genes overlapping or proximal to each state (**Supp. Fig. 13, Supp. Data File 1, Methods**). For example, among biological process terms, BivProm1 is most enriched for ‘embryonic organ morphogenesis’ genes, while TSS1 is most enriched for ‘nucleic acid metabolic process’, consistent with the bivalent [30] and the constitutively active nature of the two states, respectively. The DNase-specific state DNase1, showed distinct enrichment for CTCF-specific per-cell-type chromatin states defined in six cell types compared to other full-stack states [31] (**Fig. 3F, Supp. Fig. 14).** These CTCF-specific states have previously been suggested to be candidate insulators and may have other roles that CTCF has been implicated in such as demarcating TAD boundaries [32–35]. The DNase1 state may correspond to similar roles, particularly where the CTCF-binding is relatively stable across cell types.

Compared to other full-stack state groups, those associated with promoters (TSS, flanking promoters, bivalent promoters) and the DNase group showed lower average DNA methylation levels across cell types (**Supp. Fig 15**). Among promoter-associated states, those showing stronger enrichments with CpG Islands also show lowest methylation levels (**Fig. 3A, Supp. Fig. 15**), consistent with previous studies [36, 37]. Some promoter-associated states (TSS1-2, PromF3-5, BivProm1-2) are most enriched at the center of binding regions of polycomb repressed complex 1 and 2 (PRC1 and and PRC2) sub-units. In addition, several ReprPC and BivProm states are among the most enriched states in windows surrounding binding regions of the EZH2 and SUZ12 subunits of PRC2, consistent with these states’ association with H3K27me3 (**Supp. Fig. 16-17**). A detailed characterization of all states in terms of associated chromatin marks, genomic elements and different associated per-cell-type chromatin states across cell groups can be found in **Supplementary Data File 1-3. W**e expect it will serve as a resource for future applications using the full-stack annotations.

We verified that enrichments computed based on hg19 were highly similar to those computed for full-stack annotations mapped to hg38 (average correlation 0.99; **Methods, Supp. Fig. 18)**, ensuring the applicability of state annotations in hg38. We also confirmed that the full-stack annotations were generally more predictive of the positions of a variety of external genome annotations considered in **Fig. 3A** than two sets of per-cell-type annotations, a previous 18-state per-cell-type chromatin state annotation based on concatenated models from 127 cell types [16] and 100-state per-cell-type annotations learned independently in each cell type (**Methods**). As expected, since the full-stack model uses more data representing more cell types, the full-stack annotations had greater predictive performance in most cases (**Supp. Fig. 19-22, Supplementary Data 4)**. One of the exceptions to this was lambin B1 associated domains from Tig3 human lung fibroblasts [38], where six per-cell type annotations were more predictive, three of which are annotations for fibroblasts. We note that these evaluations were done under the assumption that a chromatin state annotation that is more predictive of well-established external genomic elements will also be more informative of less well-established classes of elements. These results suggest that the full-stack annotations will, in most cases, have greater information than any single per-cell-type annotations about localization patterns of some target genomic elements, with likely exceptions when the target of interest is specific to a certain cell type. In such cases, the corresponding per-cell type chromatin state annotation may show better recovery.

### Stacked Model Differentiates Cell-Type-Specific from Constitutive Activity

While the major groups of states outlined above can correspond to states from models producing per-cell-type annotations [3, 16], the full-stack states provide additional information. For example, the states can differentiate cell-type-specific from constitutive activities. This cell type specificity in the full-stack states is reflected in the emission parameters of cell types from different tissue groups **(Fig. 2B,C, Supp. Data. File 1)** and the overlap of per-cell-type chromatin state annotations from different cell types [39] (Supp. Figure 8-9, Supp. Data 3).

Consistent with previous findings that enhancers tend to be relatively cell-type-specific while promoters tend to be shared across cell types [3, 40], enhancer states exhibited clearer cell-type-specific associations than those of the promoter states (**Figure 2C, Supplementary Data. File 1,3**). On average, states of enhancer and weak enhancer groups (EnhW1-8, EnhA1-20) show at least two-fold higher coefficients of variations, in terms of emission probabilities for various marks, compared to states in the TSS, flanking and bivalent promoter groups (**Supp. Fig. 23**). The enhancer states differed among each other in their associations with different cell/tissue types such as brain (EnhA6), blood (EnhA7-9 and EnhWk6), digestive tissue (EnhA14-15), and embryonic stem cells (EnhA18) (**Fig. 1-2, Supp. Fig. 24-25**). These differences in cell-type-specific activities are also associated with different gene expression levels of overlapping genes with the states. For example, some blood enhancer states (EnhA8, EnhA9, EnhWk6) overlapped genes with higher average gene expression in cell types of the blood group, while some enhancer states specific to digestive group or liver tissues (EnhA14, EnhA15) showed higher gene expression in the corresponding cell or tissue types (**Fig. 3C, Supp. Fig. 10**).

Other groups of states besides enhancers also had individual states with cell-type-specific differences. For example, four of the nine states in the heterochromatin group show higher emission probabilities of H3K9me3 in only subsets of cell types (states HET1-2 with IMR90 and Epithelial cells; state HET4 with adipose, mesench, neurospheres, ESC, HSC&B-cells). Additionally, state HET9 showed strong association as heterochromatin in ESC/iPSC groups, while being mostly quiescent in other cell types based on per-cell-type annotations (**Fig. 2C, Supp. Fig. 26, Supplementary Data File 1,3**). State PromF5 is associated with putative bivalent promoter chromatin states in some blood and ESC-related cell types, but with flanking promoter states in most other cell types (**Supp. Fig. 27, Supplementary Data 1,3**). In addition, some quiescent states (Quies1-2, Quies4-5) show weak signals of H3K9me3 in specific groups of cell types (**Supplementary Data File 1**). States in the polycomb repressed and bivalent promoter groups (ReprPC1-9, BivProm1-4) also show differences in signals across cell groups, such as state ReprPC9, which showed H3K27me3 signals in only ESC/iPSC cell types (**Supplementary Data**). The ability of the stacked modeling approach to explicitly annotate both cell-type-specific and constitutive patterns for diverse classes of chromatin states highlights a complementary advantage of this approach relative to approaches that provide per-cell-type annotations.

### Full-stack states show distinct enrichments for repeat elements

As the full-stack model showed greater predictive power for repeat elements than cell-type-specific models (**Supp. Fig. 19-21**), we next analyzed which states contributed most to this power. The full-stack state enrichments for bases in repeat elements ranged from 10-fold depletion to 2-fold enrichment (**Fig. 3A**). The top ten states most enriched with repeat elements were chromatin states associated with H3K9me3 marks and in the heterochromatin, artifact, quiescent, or ZNF genes groups (**Fig. 4A-B).** Repeats being consistently enriched in H3K9me3-marked states is a natural mechanisms of cells to reduce the repeats’ risks to genome integrity, since H3K9me3 is characteristic of tightly-packed DNA (heterochromatin) that is physically inaccessible [41]. We also observed that individual full-stack states had distinct enrichments for different repeat classes (**Fig. 4C, Supp. Fig. 28**). For example, Acet1, a state associated with various acetylation marks and H3K9me3 had a 23-fold enrichment for simple repeats largely driven by (CA)n and (TG)n repeats which were 72 and 76 fold enriched and comprised 74% of all simple repeats in this state (**Supp. Fig. 28**). As (CA)n and (TG)n repeats are known to be highly polymorphic in humans [42], this suggests the possibility that signal detected in these regions may in part be due to technical issues related to deviations from the reference genome. The two states in the artifact group, GapArtf2-3, had a particularly high enrichment for satellite (181 and 145 fold, respectively) and rRNA repeat classes (75 and 580 fold, respectively) (**Fig. 4C, Supp. Fig. 28**), likely associated with sequence mapping artifacts. States in the transcription start site group, TSS1-2, were most strongly enriched with low complexity repeat class (10.5-18 fold) and most notably GC rich repeats (195-303 fold), consistent with these states being most enriched for windows of high GC content (**Supp. Fig. 28**). Moreover, the TSS1-2 states are also most enriched with tRNA class (50-61 fold) (**Supp. Fig. 28**), consistent with tRNAs being short genes [43].

**Figure 4:**
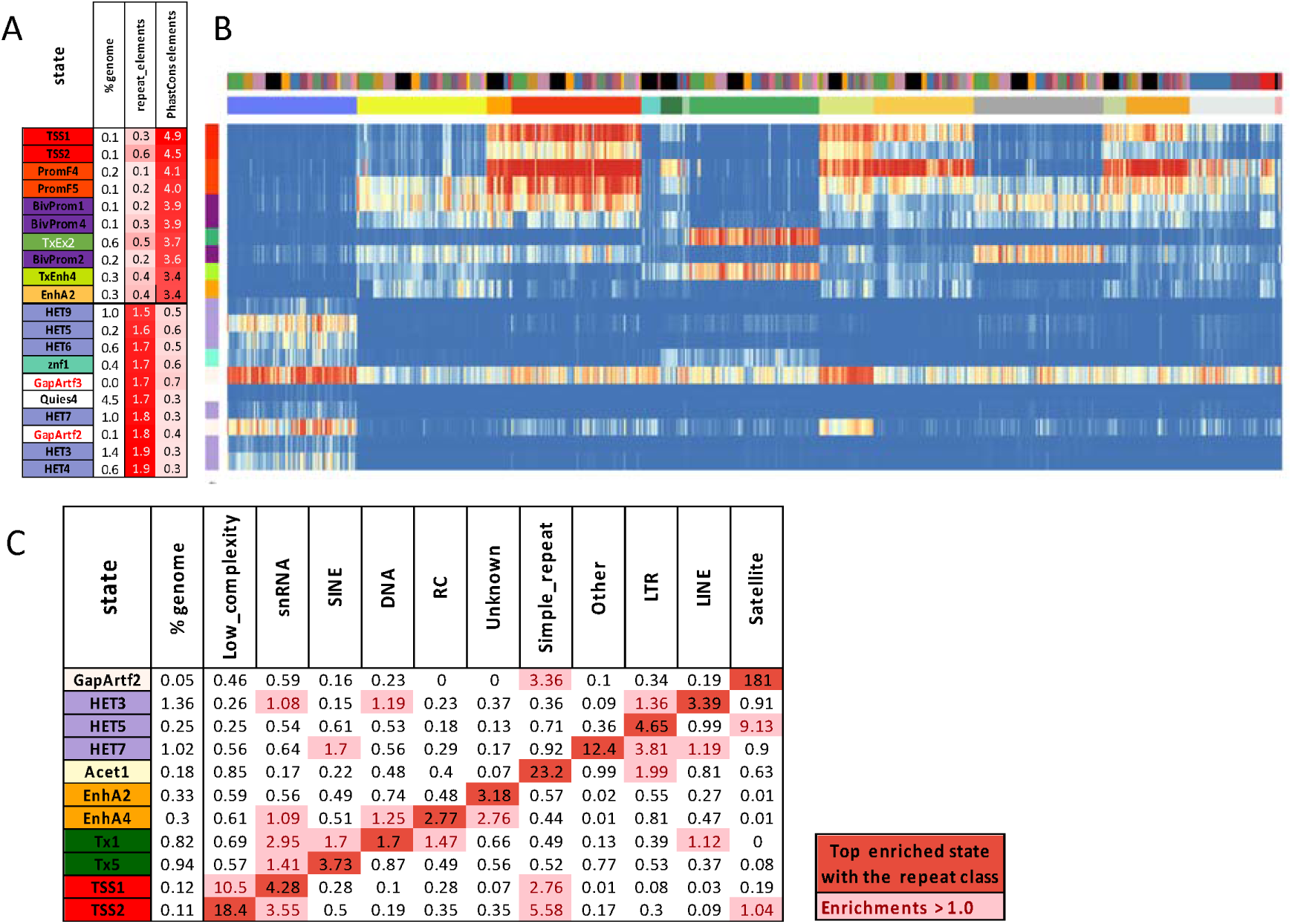
Full-stack states enrichments with conserved elements and repeat classes. **(A)** The first ten rows show the states most enriched with PhastCons elements and concurrently least enriched with RepeatMasker repeat elements, ordered by decreasing enrichments with PhastCons elements. The bottom ten rows show the states most enriched with repeat elements and concurrently least enriched with PhastCons elements, ordered by increasing enrichments with repeat elements. The columns from left to right list the state ID, the percent of the genome that each state covers, and the fold enrichments for repeat elements and PhastCons elements. **(B)** Heatmap of the state emission parameters from Fig. 2A for the subset of states highlighted in panel (A). The colors are the same in Fig. 2A. **(C)** Fold enrichments of full-stack states with different repeat classes (**Methods**). Rows correspond to states and columns to different repeat classes. Only states that are most enriched with at least one repeat class are shown. Fold enrichment values that are maximal for a given are shown in dark red. Other fold enrichments greater than one are shaded light red.

We also saw specific states associated with the largest repeat classes of the genome, SINEs, LINEs, and LTRs. SINE repeats were most enriched in state Tx5 (3.7 fold) (**Fig. 5C**), which had high emission of H3K36me3 (**Fig. 2A-B, Supp. Fig. 4-5**), consistent with previous studies showing that SINEs are more enriched in gene-rich regions and in transcription-related states based on per-cell-type annotations [21,44,45]. In contrast, LINEs are depleted in most transcription-related states, reflecting the negative selection against long-sequence insertions in or near genes [44]. LINEs are most enriched in state HET3 (3.4 folds) (**Supp. Fig. 28**), and a notable property of this state is it does not show signals of H3K27me3 and acetylation marks across cell/tissue types. This property of HET3 is a pattern that would be difficult to recognize without stacked-modeling, and was only shared with HET4 and HET9 among states in the heterochromatin group. HET4 was also the second most enriched state in the heterochromatin group for LTRs (2.0-fold) while HET9 was not enriched, but is distinct in that it identifies regions where H3K9me3 is relatively specific to cell types in the embryonic and iPSC groups. LTRs are most enriched in state HET5 (4.7 fold), and this state is marked by its highest signals of H3K9me3 compared to other states in the heterochromatin group (**Fig. 4C, Supp. Fig. 28**). LTRs showing strong enrichment with states associated with strong presence of H3K9me3 is consistent with per-cell-type chromatin state analyses [21, 45]. We also confirmed that the increased predictive power of the full-stack model over per-cell-type annotations, which was previously seen for repeat elements overall, also held for most of the individual repeat classes (**Supp. Fig. 29**). Overall, results of enrichment of full-stack state annotations for repeat classes offer further details in stratifying the states’ characteristics, and concurrently confirm and refine existing knowledge about different repeat classes chromatin state associations.

**Figure 5:**
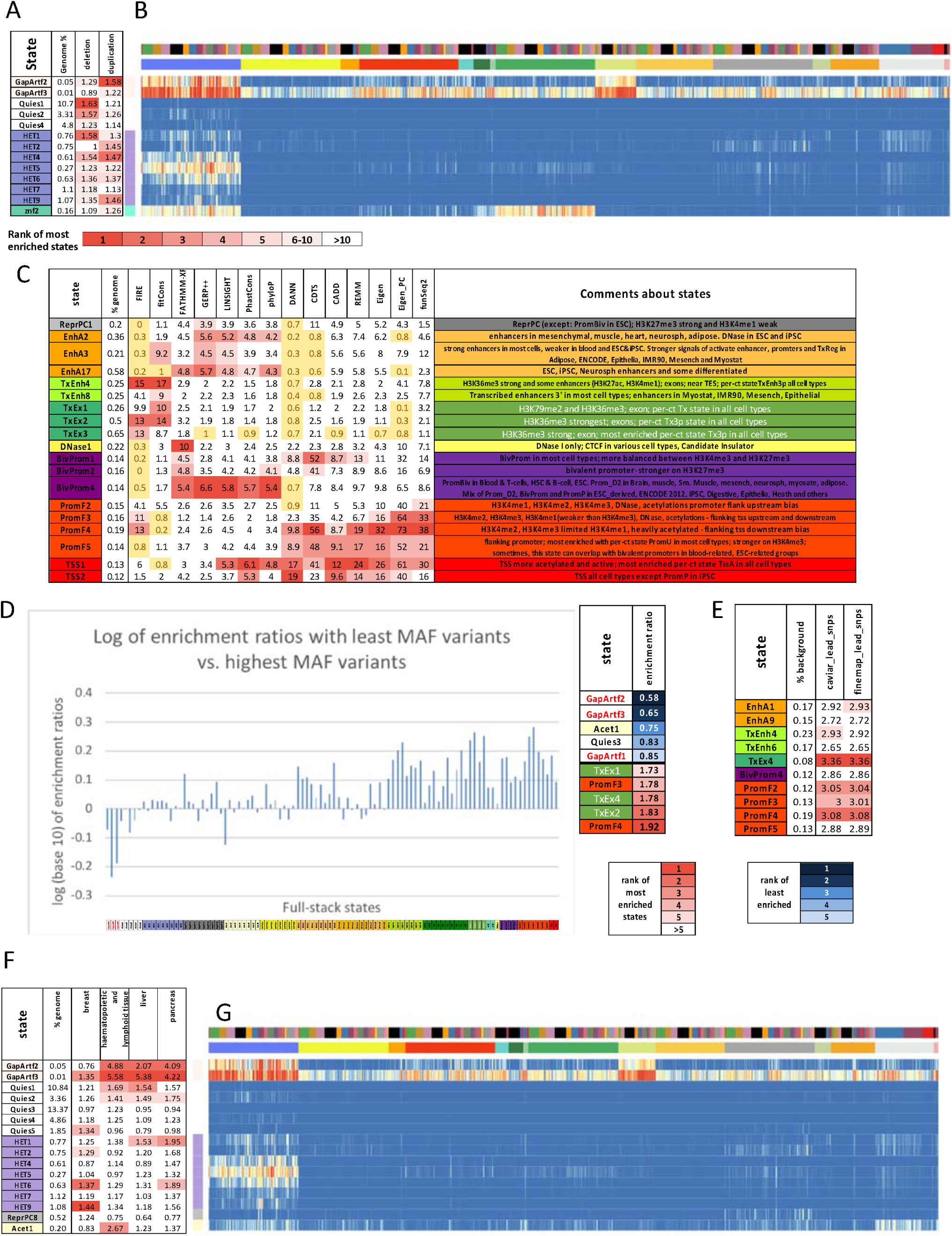
Full-stack states’ relationship with human genetic variants. **(A)** Enrichments of full-stack states with duplications and deletions from [47]. Only states that are in the top ten most enriched states are shown. Top five fold-enrichments for each class of structural variants are colored in increasing darker shades of red for higher ranked enrichments. Enrichment values below one, corresponding to depletions, are colored yellow. The columns from left to right are the state label, percent of genome the state covers, the fold enrichment for deletions, and fold enrichment for duplications. **(B)** Emission probabilities corresponding to states in **(A).** The coloring is the same as Fig. 2A. The figure highlights how states most associated with structural variants generally had higher emission of H3K9me3 compared to other chromatin marks. **(C)** Enrichments of full-stack states with top 1% prioritized bases in the non-coding genome by 14 variant prioritization scores previously analyzed [46]. Only states that are among the top five most enriched states by at least one score are shown. The top five enrichment values for each score are colored in increasing darker shades of red for higher ranked enrichment values. Enrichment values below one, corresponding to depletions, are colored in yellow. The columns from left to right are the state label, percent of the genome covered, the 14 score enrichments, and a detailed description of the state. **(D)** Log base 10 of ratios of states’ enrichment with GNOMAD variants [62] with the lowest MAFs (< 0.0001) vs. GNOMAD variants with the highest MAFs (0.4-0.5). States are ordered as in Fig. 2A. Top five states with the highest and lowest enrichment ratios are labeled to the right. **(E)** States most enriched with fine-mapped phenotypic variants against the background of common variants. Fine-mapped phenotypic variants were identified by either CAVIAR [67] or FINEMAP [68] (**Methods**). **(F)** State enrichments with somatic mutations associated with four cancer types in the non-coding genome. Only states that are among the ten most enriched with variants from at least one cancer type are shown. States in the top five are colored according to their ranks. The top five enrichment values for each cancer type are colored in increasing darker shades of red for higher ranked enrichment values. The columns are the state label, the percent of the genome the state covers, and the fold enrichments of variants from breast, haematopietic and lymphoid, liver, and pancreas cancer types. **(G)** Emission probabilities corresponding to states in (F), as subsetted from Fig. 2A. The coloring is the same as Fig. 2A. The figure highlights how states with the greatest enrichments for cancer-associated variants tend to have higher emission probabilities for H3K9me3 compared to other chromatin marks.

### Full-stack states show distinct enrichments for constrained elements and conservation states

Sequence constrained elements are another class of genomic elements that are not cell-type-specific and for which the full-stack annotations showed greater predictive power than per-cell-type annotations (**Supp. Fig. 19-21**). We next sought to better understand the relationship between full-stack states and sequence conservation annotations. We observed 10 states that had at least a 3.4 fold enrichment for PhastCons elements (**Fig. 4A**). These states were associated with the TSSs or being proximal to them (TSS1-2 and PromF4-5), transcription with strong H3K36me3 signals (TxEx2 and TxEnh4), or enhancers associated with mesenchymal, muscle, heart, neurosph, adipose (EnhA2) (**Fig. 4A-B**). In contrast, seven states (HET3-4,6-7,9, Quies4, Gap Artf2) were more than two-fold depleted for PhastCons elements, which all had more than a 1.5 fold enrichment for repeat elements (**Fig. 4A**).

To gain a more refined understanding of the relationship between the full-stack chromatin states and conservation, we analyzed their enrichment using 100 previously defined conservation states by the ConsHMM method [46]. These conservation states were defined based on the patterns of other species’ genomes aligning to or matching the human reference genome within a 100-way vertebrate alignment. We observed 29 different conservation states maximally enriched for at least one full-stack state (**Fig. 3B, Supp. Fig. 30-31**).

These states included, for example, ConsHMM state 1, a conservation state corresponding to bases aligning and matching through all vertebrates and hence most associated with constraint. ConsHMM state 1 had >= 10 fold enrichment for exon associated full-stack states TxEx1-4 and TxEnh4 (**Supp. Fig. 30**). Another ConsHMM state, state 28, which is associated with moderate aligning and matching through many vertebrates and strongly enriched around TSS and CpG islands, had a 44.5 and 47.8 fold enrichment for TSS-associated full-stack states TSS1 and TSS2, respectively (**Supp. Fig. 30**). Additionally, this conservation state is consistently the most enriched conservation state for full-stack states associated with flanking and bivalent promoters (**Fig. 3B, Supp. Fig. 30**). ConsHMM state 2, which has high aligning and matching frequencies for most mammals and a subset of non-mammalian vertebrates and previously associated with conserved enhancer regions [46], showed >2.7 fold enrichment for some full-stack enhancer states for Brain (EnhWk4 and EnhA6), ESC & iPSC (EnhA17,19 and EnhWk8), neurosph (EnhWk4, EnhA2,17), and mesenchymal, muscle, heart, adipose (EnhA2) (**Fig. 3B, Supp. Fig. 30**). ConsHMM state 100, a conservation state associated with alignment artifacts, was 10.9 fold and 2.1 fold enriched for full-stack state ZNF2 and ZNF1, respectively (**Fig. 3A-B, Supp. Fig. 30**). This is consistent with previous analysis using per-cell-type annotations showing that ConsHMM state 100 was most enriched in a ZNF gene-associated chromatin state [46]. State znf2 also showed a 5.4-fold enrichment for ConsHMM state 1 which contrasts with state znf1, which showed a 0.8 fold enrichment for ConsHMM state 1. This difference is consistent with the znf2 state’s larger fold enrichment for coding exons than znf1 (10.4 vs 1.1). The znf2 state also had a greater fold enrichment for ZNF named genes in general (68.6 vs. 20.8 fold), with those enrichment stronger and the difference greater when restricting to C2H2 annotated genes (86.8 vs. 25.1 fold) (**Fig. 3A-B, Supp. Fig 51**). Therefore, the full-stack annotation helped distinguish two ZNF-gene associated states, which are associated with distinct conservation states. As this example illustrates, the full-stack annotation captured conservation state enrichments that were generally consistent with those seen in per-cell-type annotations, but could also identify additional refined enrichment patterns.

### Specific full-stack states show distinct enrichments and depletions for structural variants

We also analyzed the enrichment of the full-stack states for overlap with structural variants (SVs) mapped in 17,795 deeply sequenced human genomes [47], and focused on the two largest classes of SVs, deletions and duplications. Abel et al., 2020 [47] analysed the enrichments of these deletions and duplications with per-cell-type chromatin states in 127 reference epigenomes [16], and observed that ZNF gene and heterochromatin states were enriched for deletions and duplications, with the enrichments being stronger in regions annotated as these states (ZNF or heterochromatin) in larger number of cell or tissue types (e.g. more constitutive HET/ZNF regions).

Consistent with those previous results, using the full-stack model, we observed that of the 13 states that were in the top 10 maximally enriched states with either deletions or duplications (1.18 fold or greater), seven were in the heterochromatin group (HET1-2,4-7,9) and one was the ZNF2 state (**Fig. 5A, Supp. Fig. 32**). The enrichment of structural variation in HET states is consistent with the notion that potentially larger effect structural variants would less likely experience negative selection in these regions of the genome. As the ZNF2 state is most enriched for a conservation state associated with putative alignment artifacts this raises the possibility that technical issues may be contributing to the SV enrichments (**Supp. Fig. 30**). The other five states included two artifact states (GapArtf2-3) and three quiescent states (Quies1-2,4) (**Fig. 5A**). The quiescent states Quies1-2,4, despite the generally low frequencies for all marks, did have higher emission probabilities for H3K9me3 compared to other chromatin marks (**Fig. 5B**).

The full-stack model was also more predictive of SV than per-cell-type annotations (**Supp. Fig. 33-34, Supplementary Data 4**). Additionally, we verified that the full-stack model had higher AUROC in predicting duplications and deletions compared to annotations obtained by ranking genomic bases based on the number of cell or tissue types that a state was observed, as in the approach of [47] (**Methods**, **Supp. Fig. 35**). These results show that the full-stack annotation can uncover enrichment patterns with SVs that are consistent with per-cell-type annotations, yet highlight states with greater predictive power and offer a more refined chromatin annotation of the regions enriched with SVs.

### Full-stack states gives insights into bases prioritized by different variant prioritization scores

Various scores have been proposed to prioritize deleterious variants in non-coding regions of the genome or genome-wide. These scores are based on either conservation or on integrating diverse sets of genomic annotations. Though the scores all serve to prioritize variants, they can vary substantially from each other and it is often not clear the differences among the types of bases that different scores prioritize. To better understand the epigenomic contexts of bases that each score tends to prioritize, we analyzed the full-stack state enrichment for bases they prioritize. As the scores we considered are not specific to a single cell type, the full-stack states have the potential to be more informative for this analysis than per-cell-type annotations. We considered a set of 14 different variant prioritization scores that were previously analyzed in the context of conservation state analysis [46]. The 14 scores for which we analyzed prioritized variants in non-coding regions were CADD(v1.4), CDTS, DANN, Eigen, Eigen-PC, FATHMM-XF, FIRE, fitCons, FunSeq2, GERP++, LINSIGHT, PhastCons, PhyloP, and REMM [48–61]. For each of these scores, we first analyzed the full-stack state enrichments for the top 1% prioritized non-coding variants relative to the background of non-coding regions on the genome (**Methods**).

In the top 1% prioritized non-coding bases, 19 states were among the top five most enriched states ranked by at least one of the 14 scores (**Fig. 5C, Supp. Fig. 36-37**). These 19 states include nine states in promoter-associated groups, five states in enhancers-related groups, three states in the exon-associated transcription group, one polycomb repressed state, and one DNase state (**Fig. 5C**). Seven scores (DANN, Eigen, Eigen_PC, funSeq2, CDTS, CADD and REMM) had their top five enriched states exclusively associated with promoter and TSS states, with enrichments ranging between 8.6 and 70 fold (PromF2-5, TSS1-2, BivProm1-2,4) (**Fig. 5C**). In contrast, the fitCons score showed depletions for three of these states and relatively weaker enrichment for the others. This difference might be related to fitCons’ approach of prioritizing bases showing depletion of human genetic polymorphisms, potentially without sufficiently accounting for the increased mutation rates in regions with high CpG content that are observed in promoter-associated states (**Supp. Fig. 42**) [62]. FIRE’s prioritized variants showed depletions in the bivalent promoter states (BivProm) and PromF5, which have generally lower average gene expression across cell types (**Fig. 3C**). This depletion is reflective of the fact that FIRE was trained to prioritize variants in cis-expression quantitative trait loci (cis-eQTLs) in one cell type (LCL) [54], and few eQTLs are expected to be proximal to genes with limited or no expression in that cell type. Enhancer states EnhA2-3,17 were among the states in the top five most enriched for FATHMM, GERP++, LINSIGHT, PhastCons, and PhyloP prioritized non-coding variants. In contrast, FIRE, DANN and CDTS were depleted for prioritized variants in all these enhancer states (**Fig. 5C**). FIRE and fitCons showed strong enrichment for exon states (TxEx1-3), which are associated with coding regions, even though coding bases were excluded in this analysis (Fig. 5C). FATHMM had the greatest relative enrichment for the primary DNase state associated with CTCF cell type-specific chromatin states (DNase1) (∼10 fold), and was the only score for which this state was among the top five most enriched states (Fig. 5C, Supp. Fig. 36).

We conducted similar analyses based on top 5% and 10% prioritized non-coding variants and observed relatively similar patterns of enrichments, though there did exist some differences at these thresholds (**Supp. Fig. 36, 38-39**). One difference was that alignment artifact states GapArtf2-3 were among the top two most enriched states with top non-coding bases prioritized by FATHMM-XF, while a number of other scores showed depletions with these states (**Supp. Fig. 36**). In addition, we analyzed top 1%, 5%, and 10% prioritized variants genome-wide from 12 of the scores (**Methods**) (**Supp. Fig. 37-40**). Compared to the non-coding analysis, we saw a majority scores that have exon-associated transcription states (TxEx1-TxEx4) among the top five enriched states with top 1% variants genome-wide, while we saw no enhancer state among the top five enriched states with top 1% variants by any score and only one enhancer state among the top five by one score (GERP++) for top 5% and 10% variants.

Overall, this analysis shows that the scores tend to prioritize bases in different epigenetic contexts. As these scores vary in the genomic features selected as input, and the predictive model for scoring bases, it is expected that different methods may show higher scores for in different classes of genomic contexts. By analyzing the state enrichments, one can gain some expectations for what types of evaluation criteria different scores might perform better, but we note that this analysis is not trying to directly conclude one method is preferred. Also, while in general it is difficult to conclude confidently what enrichments are due to technical or biological biases, by comparing enrichments across scores and considering what else is known about the states, one can still gain insights into this. For example, the inconsistent enrichments of different methods for prioritized variants in GapArtf2-3 states (**Supp. Fig. 36**), along with these states’ association with sequencing artifacts, is suggestive of technical biases. Similarly, DANN’s top 1% non-coding bases showing enrichments in five heterochromatin states, while not showing any enrichments in enhancer states, and no other scores showing enrichments in heterochromatin states, is also suggestive of technical biases (**Supp. Fig. 37**).

We verified that the full-stack annotation showed the highest AUROC in recovering the top 1% non-coding variants compared to all 18-state per-cell-type annotations from a concatenated model for all 14 scores (**Supp. Fig. 33**). Compared to all 100-state per-cell-type annotations from independent models, the full-stack model showed the highest AUROC for 13 out of 14 scores in all 127 cell types (**Supp. Fig. 33**).

### Full-stack states show distinct enrichments and depletions for human genetic variation

We next analyzed full-stack states for their enrichment with human genetic sequence variation. We calculated enrichments of full-stack states with genetic variants sequenced in 15,708 genomes from unrelated individuals in the GNOMAD database stratified by minor allele frequencies (MAFs) [62]. Across eleven ranges of MAFs, the state enrichments ranged from a 2-fold enrichment to a 4-fold depletion (**Supp. Fig. 42**). As expected, the state associated with assembly gaps (GapArtf1) is most depleted with variants, regardless of the MAF range. At the other extreme, state Acet1, which is associated with simple repeats, is the most enriched state with variants for all ten minor allele frequency (MAF) ranges that are greater than 0.0001, with fold enrichments between 1.8 and 2.0 (**Supp. Fig. 42**). We verified that the high enrichment for state Acet1 was not specific to GNOMAD’s calling of variants as it had a 2.0 fold enriched with common variants from dbSNP (**Methods**) (**Supp. Fig. 42**). TSS and promoters associated states, PromF4 and TSS1-2, were maximally enriched for variants in the lowest range of MAF (0 < MAF <= 0.0001), 1.5-1.7 fold. The enrichment of variants for these states decreased as the MAF ranges increased, falling to 0.8-1.2 fold for variants of the highest range of MAF (0.4-0.5) (**Supp. Fig. 42**). The high enrichment for states PromF4 and TSS1-2 for rare variants, despite their being the most enriched states with PhastCons conserved elements, can be explained by these states’ high enrichment of CpG dinucleotides, which are associated with higher mutation rates (**Fig. 3A, Supp. Fig. 42)** [62]. At the same time, purifying selection can have a weaker effect on large-effect rare variants than on large-effect common variants. We also observed the pattern of decreasing enrichments for variants with increasing MAF in other states associated with transcriptional activities, enhancers, DNase, or promoters (**Supp. Fig. 42**). This pattern was not observed in most states from other groups such as heterochromatin, polycomb repressed, quiescent, and acetylations only (**Supp. Fig. 42**).

To better identify states with a depletion of common variants that are more likely due to selection, we ranked states based on their ratios of enrichments for the rarest variants (MAF < 0.0001) relative to the most common variants (MAF 0.4-0.5) (**Fig. 5D**). The states with the highest ratio included a number of flanking promoter (PromF3-4) and exon-transcription states (TxEx1,2,4) that were also associated with strong sequence conservation across species (**Fig. 3B, Fig. 5D**). These results are consistent with previous analyses supporting a depletion of common human genetic variation in evolutionary conserved regions (Lindblad-Toh et al., 2011). States associated with assembly gaps and alignment artifacts (GapArtf1-3), quiescent (Quies3), or acetylations and simple repeats (Acet1) were most depleted for rare variants relative to the common variant enrichment (**Fig. 5D**).

### Full-stack states show enrichment for phenotype-associated genetic variants

We next analyzed the relationship between the full-stack states and phenotypic associated genetic variants. We first evaluated the enrichment of the full-stack state for variants curated into the Genome-wide Association Study (GWAS) catalog relative to a background of common variation [63] (**Methods**). This revealed six states with at least a two-fold enrichment (**Supp. Fig. 43**). Four of these states, TxEx1- 2,4 and TxEnh4, were all transcription associated states that are >= 10-fold enriched with coding sequences and >=11 fold for ConsHMM state 1, associated with the most constraint in a sequence alignment of 100 vertebrates (**Fig. 3B**). This observation is consistent with previous results that GWAS catalog variants show enrichments for coding sequence and sequence constrained bases [46,64,65]. The other two states with greater than two-fold enrichment for GWAS catalog variants relative to common variants were two promoter states, PromF2-3 (**Supp. Fig. 43**). On the other hand, four states were more than two-fold depleted for GWAS catalog variants, and were associated with artifacts (GapArtf2-3), or quiescent and polycomb repressed states with weak signals of H3K9me3 (Quies5) or H3K27me3 (ReprPC8) (**Supp. Fig. 43**). Both Quies5 and ReprPC8 are highly specific to chrX, 18.3 and 19.1 fold enriched respectively (**Supp. Fig. 43**).

We also analyzed the full-stack state enrichments for fine-mapped variants previously generated from a large collection of GWAS studies from the UK Biobank database and other public databases [66]. Specifically, we considered separately the fine mapped variants from two fine-mapping methods, CAVIAR [67] and FINEMAP [68], for 3052 traits. For each method and trait, we identified the single variants that had the greatest probability of being causal at a set of distinct loci, and computed the enrichment of these variants for the full-stack states relative to a background of common variants (**Methods**).

Fold enrichment results of full-stack states for the most likely causal variants were highly consistent between fine-mapping methods (FINEMAP and CAVIAR) (**Supp. Fig. 44**). The ten states maximally enriched with fine-mapped variants relative to common variants, which were the same states by both methods, included five states associated with flanking and bivalent promoter activities (PromF2-5, BivProm4), an enhancer state in blood and thymus (EnhA9) and an enhancer state in most other cell types except blood (EnhA1), and three highly conserved transcription-associated states (TxEnh4,6, TxEx4) (**Fig. 5E**). Notably, five of 10 states maximally enriched with fine-mapped variants, PromF2-5, and BivProm4, were associated with promoter regions and also among the 19 states most enriched with top 1% prioritized variants by at least two of the 14 different variant prioritization scores (**Fig. 5E, C**). These results show that there are agreements in the types of chromatin states preferentially overlapped by phenotype-associated fine mapped variants and variants predicted to have greater effects based on variant prioritization scores. We also confirmed that the full-stack model consistently resulted in higher AUROC in predicting locations of fine-mapped variants within a background of common variants, compared to the per-cell-type annotations in all cell types (**Supp. Fig. 45-46**).

### Full-stack states show enrichments for cancer-associated variants

In addition to investigating germline variants, we also investigated the enrichment of full-stack states for somatic variants identified from whole genome sequencing of cancer samples. We analyzed data of variants from four cancer types that have the largest number of somatic variants in the COSMIC database [69]: liver, breast, pancreas and haematopoietic_and_lymphoid_tissue (**Methods**). Sixteen states were among the top 10 most enriched with at least one type of cancer’s associated variants (1.2-1.4 fold in breast cancer, 1.2-5.6 fold in lymphoid cancer, 1.2-5.4 in liver cancer, 1.4-4.2 in pancreas cancer) (**Fig. 5F**). Among these 16 states, 15 states showed higher signals of H3K9me3 compared to most other chromatin marks, including seven states in heterochromatin group (HET1-2, 4-7,9), four states in quiescent group with weak emissions of H3K9me3 (Ques1-2,4-5), one state in the polycomb repressed group with weak signals of H3K9me3 and H3K27me3 (ReprPC8), one state in the acetylation group with signals of H3K9me3 and various acetylation marks (Acet1), two artifact-associated states with higher signals of H3K9me3 and DNase relative to other marks (GapArtf2-3) (**Fig. 5G**). This pattern of H3K9me3-associated states being enriched with somatic mutations in cancer is previously confirmed in multiple studies where H3K9me3 and other repressive epigenetic features showed positive association with mutation density across different types of cancer cells [70–72]. One possible explanation for this association is the more limited access of DNA mismatch repair machinery in these regions due to the tightly packed nature of the genome in heterochromatin [73, 74]. Notably, the GapArtf2-3 states, which had strong satellite repeats enrichments (**Fig. 4C, Supp. Fig. 28**) were the top two most enriched states with somatic variants associated with liver, pancreas and haematopoietic and lymphoid tissue (haem-lymphoid) cancers (2.0-5.6 folds enriched) (**Fig. 5F, Supp. Fig. 47**). We suspect that the enrichments in these putative alignment artifact states are driven at least in part by false variant calls due to sequence mapping errors associated with these regions. Similarly, enrichments of somatic mutations in haem-lymphoid cancer in state Acet1 is also suggestive of the possibility of false calls given this state’s combination of H3K9me3 and acetylation signal and enrichment for simple repeats (**Fig. 2, 4C, Supp. Fig. 42**). We note that the presence of cancer variants is better recovered by full-stack annotation as compared to the per-cell-type chromatin state annotations for all four cancer types (**Supp. Fig. 40-41**).

## Discussion

We demonstrated a large-scale application of the stacked modeling approach of ChromHMM using over a thousand epigenomic datasets to annotate the human genome. In the datasets, 32 chromatin marks and 127 reference epigenomes were represented. We note that even though not every chromatin mark was profiled in every reference epigenome, we were still able to directly apply the stacked modeling to such data. Previously, concatenated models were applied to observed and imputed data [39], however, we chose not to use imputed data as input to the full-stack model since imputed data would still be based on the same observed input data used in stacked-modeling. We conducted extensive enrichment analyses of the states with various other genomic annotations and datasets, including gene features, genetic variation, repetitive elements, comparative genomic annotations, and bases prioritized by different variant prioritization scores. These analyses highlighted diverse enrichment patterns of the states. Using these enrichments along with the model parameters, we provided a detailed characterization of each of the 100 states in the model.

We grouped these 100 states into 16 groups that included promoters, enhancers, transcribed regions, polycomb repressed regions, zinc finger genes among others. We also highlighted important distinctions among states within the groups. In many cases, identifying these distinctions was enabled by the full-stack modeling using data from multiple cell types for genome annotation. For example, we identified enhancer and repressive states that were active in different subsets of cell types. We also highlighted how different states in some of the groups such as those associated with transcribed and ZNF genes showed distinct enrichments for conservation states. Overall, the full-stack model showed enrichment patterns supporting observations held for per-cell-type annotations, yet it provided more detailed stratification of genomic regions into chromatin states with heterogeneous associations with other genomic information. We provide extensive characterizations of full-stack states in **Supplementary Data Files 1-5** that will be resources in future applications of the full-stack annotations.

The full-stack modeling has advantages to commonly used per-cell-type chromatin state annotations in several respects. First, the full-stack model learns patterns of signals of the same or different assays across cell types, hence can provide a unified view of all the data and directly uncover states that correspond to constitutive or cell-type-specific activities. For example, a state from the model, HET9, was associated with only the mark H3K9me3 specifically in ESCs and iPSCs even though this mark is typically associated with constitutive repression. Second, the full-stack annotation consistently showed better recovery of various genomic features compared to per-cell-type annotations. This improvement is expected since full-stack models can leverage information from multiple cell types for genome annotations. Third, in cases where it is not desirable to focus on only one specific cell or tissue for analysis, the full-stack modeling can bypass the need to pick one such cell or tissue type or to consider a large number of different per-cell-type chromatin state annotations simultaneously. Such cases may arise when studying other genomic information that is not inherently cell-type-specific such as genome variation and sequence conservation.

Despite these advantages, there are trade-offs in using the stacked modeling approach, and we emphasize that the stacked modeling approach should be considered a complement to and not a replacement of the per-cell-type annotations. Compared to typical concatenated models, the full-stack model has increased model complexity because of the increased number of parameters from the larger number of states and input features, which can make interpreting some model states relatively more difficult. Additionally, if one is interested in a specific cell type, then corresponding per-cell-type annotations can have advantages in that all the annotations are directly informative about the chromatin state in the cell type of interest. An additional trade-off is that with the stacked model, it is not possible to incorporate additional data without relearning a model, while for a concatenated-model one can annotate a new cell based on an existing model, provided that the set of marks in the new cell type are the same as the existing model. We also note that post-hoc per-cell-type state annotations can also be used for cross-cell type analyses, particularly on a per-state basis by analyzing the frequency of a specific state across cell types. While per-state analyses using per-cell-type concatenated annotations can be relatively straightforward and informative, they give only partial and potentially oversimplified views of all the data, ignoring distinctions among different per-cell-type states. Whether to use per-cell-type annotations or full-stack annotations will depend on the specific application. Per-cell-type annotations may be preferable when one is interested in studying a specific cell type, while full-stack annotations can be preferred in joint analyses of multiple cell types.

We expect many applications of the full-stack annotations that we generated here and they have already begun to be applied in other work [75–77], which we expect to further elucidate the biological significance of different states. The full-stack annotation can be used as a resource to interpret genetic variation. A possible avenue for future work is to incorporate the full-stack annotation into scoring methods to better predict genetic variants’ phenotypic influences. Given the increasing availability of epigenomic datasets [18], future work could also learn new stack-models to incorporate such data. The state characterizations (**Supplementary Data File 1-5**) and analyses introduced through this work will be useful in interpreting biological implications of new models’ states. Future work can also include training and deriving the full-stack annotations for key model organisms such as mice. This work provides a new annotation resource for studying the human genome, non-coding genetic variants, and their association with diseases.

## Methods

### Input data and processing

We obtained coordinates of reads aligned to Human hg19 in .tagAlign format for the consolidated epigenomes as processed by the Roadmap Epigenomics Consortium from https://egg2.wustl.edu/roadmap/data/byFileType/alignments/consolidated/. In total we obtained data for 1032 experiments and their corresponding input control data. The experiments correspond to 127 reference epigenomes, 111 of which were generated by the Roadmap Epigenomics Consortium and 16 were generated by the ENCODE Consortium. Of the 1032 experiments, 979 were of ChIP-seq data targeting 31 different epigenetic marks and 53 were of DNase-seq (**Sup Fig. 2**). For each of the 127 reference epigenomes there was a single ChIP-seq input control experiment. For the 53 reference epigenomes that had a DNase-seq experiment available there was an additional DNase control file.

We next binarized the data at 200 base pair resolution using the BinarizeBed command of ChromHMM (v.1.18). To apply BinarizeBed in stacked mode we generated a cell_mark_file input table for ChromHMM with four tab-delimited columns. The first column had the word ‘genome’ for all datasets, the second column contained entries of the form ‘<EID>-<MARK>’ where ‘EID’ is the epigenome ID and ‘mark’ is the mark name, the third column specifies the name of the corresponding file with aligned reads, and the fourth column is the name of the file with the corresponding control reads. Each row in the table corresponds to one of the 1032 experiments.

In order to reduce the memory and time needed to execute BinarizeBed on a large number of datasets, we split the cell_mark_file table into 104 smaller tables with each table having at most 10 entries corresponding to at most 10 datasets to be processed. This was done with a custom script, but the same functionality has been included with the ‘-splitcols’ and ‘-k’ flags of BinarizedBed in ChromHMM v1.22. We then ran BinarizeBed in parallel for each of these smaller cell_mark_file tables and generated output into separate sub-directories. We ran BinarizeBed with the option ‘-gzip’ which generates gzipped files.

To merge data from the 104 subdirectories from the previous step into files containing binarized data of all experiments, we ran the command ‘MergeBinary’, which we added in v1.18 of ChromHMM. We ran the command with the options ‘-gzip -splitrows’. The ‘-splitrows’ option generates multiple files of merged binarized data for each chromosome, where, under the default settings that we used, each file contains data for a genomic region of at most 1MB. Splitting each chromosome into smaller regions allows the model learning step of ChromHMM to scale in terms of memory and time to the large number of input data tracks (i.e. features) that we were using. We used chr1-22, chrX, chrY, and chrM in the binarization and model learning.

### Training full-stack model and generating genome-wide state annotations

We learned the full-stack chromatin state model for the 1032 datasets using the LearnModel command of ChromHMM (v1.18). This version of ChromHMM includes several options that we added to improve the scalability when training with large numbers of features. One of these features was to randomly sample different segments of the genome for training during each iteration, instead of training on the full genome. This sampling strategy was previously used by ConsHMM [46], which was built on top of ChromHMM. We note that this sampling procedure can also be applied to learn concatenated models, in which case there would be no requirement that the same segments are sampled in each cell type. However, sampling can be unnecessary for typical instances of learning concatenated chromatin state models, given that it usually involves fewer different inputs to the model, fewer number of states, and in training these models, ChromHMM is able to tolerate more parallel cores without reaching memory limits.

To learn the full-stack model with input data processed as outlined above, we used ChromHMM’s LearnModel command with the options ‘-splitrows -holdcolumnorder -pseudo -many -p 6 -n 300 -d -1 -lowmem -gzip’. The ‘-splitrows’ flag informs ChromHMM that binarized data for a chromosome is split into multiple files, which reduces the memory requirements and allows ChromHMM to select a subset of the genome to train on for each iteration. The ‘-holdcolumnorder’ flag prevents ChromHMM from reordering the columns of the output emission matrix, which saves time when there are a large number of features.

The ‘-pseudo’ flag specifies that in each update of model parameters, ChromHMM adds a pseudo count of one to the numbers of observations of transition between each pair of states, presence and absence of each mark from each state, and initial state assignments of the training chromatin state sequence. This prevents model parameters from being set to zero, which is needed for numerical stability when some features are sparse and ChromHMM does not train on the full genome in each iteration.

The ‘-many’ flag specifies ChromHMM to use an alternative procedure for calculating the state posterior probabilities that is more numerically stable when there are a large number of features. The procedure is designed to prevent all states from having zero posterior probability at any genomic position, which can happen due to the limits of floating-point precision. The procedure does this by leveraging the observation that only the relative product of emission probabilities across states are needed at each position to determine the posterior probabilities. Specifically, for each position, the procedure initializes the product of emission probabilities for all features, i.e. the emission product, from each state to one. For each feature, the procedure then multiplies the current emission products from each state by the emission probability of the feature in the state, and divides all the resulting products by their maximum to obtain updated emission products. We iteratively repeat these steps of multiplication and normalization until all features have been included into the calculation of relative emission products across states.

The ‘-p 6’ flag specifies to ChromHMM to train the model in parallel using 6 processors. The ‘-n 300’ flag specifies to ChromHMM to randomly pick 300 files of binarized data, corresponding to 300 regions of 1 MB (or less if the last segment of the chromosome was selected) for training in each iteration. The ‘-d -1’ option has ChromHMM not to require an evaluated likelihood improvement between iterations to continue training since evaluated likelihood decreases are expected, as on each iteration the likelihood is evaluated on a different subset of data. The ‘-lowmem’ flag has ChromHMM reduce main memory usage by not storing in main memory all the input data and instead re-loading from disk when needed. The asymptotic worst-case time and memory of the model learning is discussed in the S**upplementary Information**.

### Choice of number of states

We trained full-stack models with 10-120 states, in 5 state increments, using the data and procedure outlined above. For each of these models, we calculated an estimated Akaike Information Criterion (AIC) [78] and Bayesian Information Criterion (BIC) [79] value based on a subset of the genome (**Supp. Fig. 3**). AIC and BIC are calculated based on the log likelihood for 300 random 1Mb regions outputted by ChromHMM from the last training iteration. In general, both the AIC and BIC decrease as the numbers of states increase, but with diminishing improvements. We also applied the CompareModels command of ChromHMM [25] with the 100-state model as a reference model, which reports, for each state of the 100-state model, the maximum correlations of emission parameters between the state in the 100-state model and any state for each other models (**Supp. Fig. 3C**). We conducted a similar analysis with the emission parameters of H3K4me1, hence for each state in each model, we obtained emission probabilities of H3K4me1 in 127 cell types. For this analysis, for each of the 19 tissue groups previously defined [16], we calculated the correlation of each state’s H3K4me1 emission parameters with a binary vector indicating if the cell type in each parameter is in the tissue group (1) or not (0). We then report, for each tissue group, the maximum correlations among all states in each model (**Supp. Fig. 3D**). These analyses showed, for instance, that a state corresponding to Brain-specific enhancers in the 100-state model, EnhA6, was well captured in models with 55-states or more (correlation of >=0.98 with states in models with >=55 states and correlation >=0.78 for H3K4me1-emissions with the Brain binary vector). A state characterized as enhancers specific to Huvec cells in the 100-state model, EnhA20, was well captured in models with 100 or more states (correlation >=1.00 based on all marks’ emission parameters).

Additionally, for models with 20, 40, 60, 80, 100 and 120 states, we also produced genome annotations and then quantitatively compared the chromatin state annotations from models in terms of their power to predict locations of various other genomic annotations not used in the model training: Exon, Gene Body, TSS, TSS2kb, CpG Islands, TES, laminB1lads elements (listed in section *External Annotation Sources* section). Specifically, we evaluated the predictive power using the AUROCs that are calculated as described in a subsection below. Across different genomic contexts, as the number of full-stack states increased, the AUROC increased, but with diminishing improvements as the number of states increased (**Supp. Fig. 3A**).

To balance the additional information available in models with an increased number of states, while keeping the number of states manageable for interpretation and downstream analysis, we choose to focus on a model with 100 states. We note that this choice is greater than previously used for concatenated models with per-cell-type annotations [3,16,21], which reflects the additional information available for genome annotation based on the large number of datasets spanning many cell types that we are using.

### Lifting chromatin state annotations to hg38

The full stack chromatin state annotations were learned directly in hg19, as this was the assembly for which uniformly processed data from the Roadmap Epigenomics integrative analysis was available. Learning full-stack annotation directly in hg19 allowed direct comparison with existing per-cell-type annotations inferred. We also generated a version of full-stack annotation in hg38 by lifting over the original annotation from hg19 to hg38. To do this, we first wrote the hg19 chromatin state annotation into .bed format such that each line corresponds to a 200bp interval. We then used the liftOver tool [80] with default parameters to generate the annotation in hg38. We did not annotate bases in hg38, if multiple bases in hg19 mapped to it. In total, there are 1,186,379 200-bp segments that were not mapped from hg19 to hg38, of which 98.7% fall into an assembly gap and 99.6% fall into the full-stack state primarily associated assembly gaps (GapArtf1) (**Supp. Fig. 50**). In hg38 on chr1-22, X, and Y, 92.9% of bases are annotated to a state, and that number increases to 97.1% when excluding assembly gaps. We verified that we saw highly similar state fold enrichments for similar annotations between hg19 and hg38 (**Supp. Fig. 18**). The sources of external annotations from hg38 are outlined in section “External annotation sources” below.

### Summary sets of experiments

To construct a summary visualization of the emission parameters with a reduced set of features that approximate the annotation from the full model, we applied a greedy search over the 1032 input datasets as described in **Supplementary Methods.** We applied this procedure to reduce the 1032 input datasets to 80 summary datasets.

### Identifying states with differential association of marks for individual tissue groups

For each state, we tested for combinations of the 8 most profiled marks, and 19 tissue groups previously defined [16], whether the emission probabilities of features associated with one chromatin mark and in one tissue group was significantly greater than those of features associated with the same mark and not in the tissue group. The eight marks that we tested were H3K9me3, H3K4me1, H3K4me3, H3K27me3, H3K36me3, H3K27ac, H3K9ac, and DNase. H3K27ac, H3K9ac and DNase were profiled in 98, 62 and 53 reference epigenomes, respectively, and the remaining five marks in 127 reference epigenomes. For tests involving H3K27ac, H3K9ac, and DNase, we excluded tissue groups for which there were no experiments. In total, there were 14,200 tests among 100 states, 8 chromatin marks and 19 tissue groups. For each combination of state, chromatin mark and tissue group being tested, we applied a one-sided Mann-Whitney test to test whether the emission probabilities of the state for the features associated with the tested mark in the tested tissue group are greater than those in other tissue groups. The Bonferroni-corrected p-value threshold based on a significance level of 0.05 to declare a test significant was 3.5e-6.

### Computing coefficients of variation across different tissue groups

For each state, we looked into the emission probabilities of experiments associated with six chromatin marks strongly associated with promoter and enhancer activities (DNase, H3K27ac, H3K4me1, H3K4me2, H3K4me3, H3K9ac). We grouped these experiments based on their associated chromatin mark and tissue groups, and calculated the average emission probabilities of experiments in each chromatin mark-tissue group combination. For each state and chromatin mark combination, we then calculated the coefficient of variation across different tissue groups, in terms of average emission probabilities from the previous step. For each group of states, we averaged the resulting coefficients of variation across states of the same group. The results show the average coefficients of variation of emission probabilities across different tissue groups for each state group-chromatin mark combination.

### Computing fold enrichments for other annotations

All overlap enrichments for external annotations were computed using the ChromHMM OverlapEnrichment command. We used the ‘-b 1’ flag, which specifies a binning resolution of the annotations. This ‘-b 1’ flag is necessary when computing enrichments based on the hg38 liftOver annotations, which no longer respects the 200bp segment coordinate intervals from hg19. Including this flag gives the same results when applied to annotations from hg19 with 200bp segments, though with extra computational costs. We also included the ‘-lowmem’ flag to specify the lower memory usage option. The ChromHMM command OverlapEnrichment computes fold enrichment between chromatin states and provided external annotations relative to a uniform genome-wide background distribution. More specifically, the fold enrichments are calculated as:

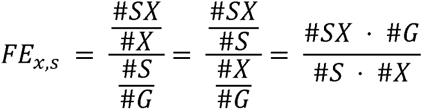

where

*FE_x,s_*: fold enrichment of state *s* in genomic context *x*

*#S*: number of genomic positions belonging to the state *S*

*#X*: number of genomic positions where genomic context *X* is present

*#SX*: number of genomic bins that overlap both state *S* and genomic context *X*

*#G*: number of genomic positions in the entire genome

### Enrichment and estimated probabilities of overlap with per-cell-type chromatin state annotations

We obtained per-cell type chromatin state annotations based on a 25-state ChromHMM model learned using concatenated approach for 127 reference epigenomes, which we will refer to as cell types for ease of presentation, from the Roadmap Epigenomics project [16, 39]. This model was trained based on observed and imputed data for 12-marks. We hereafter refer to this model as the CT-25-state model. As per the design of the concatenated approach of ChromHMM, CT-25-state model generates per-cell-type chromatin state annotations for each of the 127 cell types, and the 25 states’ characteristics are shared across 127 cell types. For each of these 127 cell types, we calculated overlap enrichments between the 100 full-stack states and the CT-25-states, resulting in 127 tables of size 100-by-25. We summarized this information by reporting, for each of the 100 full-stack states, and 127 cell types, the per-cell-type state in CT-25-state model that is maximally enriched, resulting in a 100-by-127 table (**Supp. Fig. 8**). We also provided detailed comments about the patterns of maximum-enriched-states observed across 127 cell types for each full-stack state in the **Supplementary Data File 3** to serve as a resource for future applications. We also reported, for each of the 100 full-stack states and each per-cell-type-25-state, the maximum and median values of fold enrichments across 127 cell types (**Supplementary Data File 3**).

In addition, we also estimated for each combination of (1) CT-25 state, (2) cell type group and (3) full-stack state, the probability that a genomic position being annotated as the corresponding full-stack state will overlap with the corresponding per-cell type state in a cell type from the corresponding cell group. The 19 groups of cell types were previously defined by the Roadmap Epigenomics Consortium [16]. To compute the target estimate probabilities, for each full-stack state, we sampled 100 genomic bins (each of length 200bp) that are assigned to that full-stack state. Second, in each of the 127 cell types, we report the frequency that the sampled regions of each full-stack state overlapped with each CT-25-state. We repeated such a process 21 times. We then calculated the average frequencies of overlap between each full-stack state and each CT-25-state, across 21 random samplings and across the cell types in each group (Example: Blood, ESC). This results in a table of size 100 full-stack states by 475 combinations of CT-25 states and 19 cell groups, with each cell showing the values of estimated probabilities (**Supp. Fig. 9**). These values, along with detailed comments about patterns of these overlap probabilities for each full-stack state, are available in **Supplementary Data File 3**.

### Receiver operator characteristic curve analysis for predicting external annotations

To evaluate how well the chromatin state annotations from different ChromHMM models can inform us about the position of external genomic annotations, we computed the Receiver Operator Characteristic (ROC) in a procedure as follows: First, we divided the genome into 200bp bins, and randomly partitioned 50% of the bins for training and the remaining 50% for testing. Second, we computed the enrichment of the target external annotation with each chromatin state on the training data, and ranked states in decreasing order of such enrichments. We used this ranking of states to iteratively add genomic bases assigned to the states as our predictions of bases that overlap the target annotation in the testing dataset. Based on the overlap of the predictions and the target annotation at each iteration, we plotted ROC curves and summarized the information by computing area under the ROC curves (AUROC).

### Per-cell-type annotations used to compare against full-stack annotations

In evaluating how predictive the full-stack model is at annotating external genomic elements, we compared the full-stack model to two sets of per-cell-type chromatin state annotations in terms of their ability to predict external annotations. One set of per-cell type annotations was the 18-state ChromHMM from Roadmap Epigenomic Project [16], which was based on a model trained using the concatenated approach and observed data of six chromatin marks (H3K4me1, H3K4me3, H3K9me3, H3K27ac, H3K27me3 and H3K36me3) in 98 cell types. In this model, we have a common set of state definitions across cell types, but unique state annotations for each cell type. The second set of per-cell type annotations were based on models learned independently in each of the 127 cell types. In learning these models, we partitioned the 1032 datasets used to learn the full-stack model into 127 subsets based on their associated cell type. For each of the 127 cell types, we applied ChromHMM to learn a 100-state per-cell-type ChromHMM model using only the observed data in the corresponding cell type. The number of states is similar to that in the full-stack model, to control for this variable in the evaluation. This process generates 127 models, each used to generate per-cell-type annotations in one cell type. The independent model learning approach of ChromHMM differs from the concatenated approach because the model parameters (state emission, transition, and initial probabilities) are different for different cell types, while state parameters in concatenated model are shared across cell-types. However, these two approaches both produce chromatin state annotations on a per-cell type basis. We learned these independent models with the same ChromHMM parameters as described above for the full-stack model, with the exception of using the ‘-init random’ flag to randomly initialize models’ parameters. Even when we specified the number of states to ChromHMM as 100, however, we note that due to the large number of states relative to the input tracks, for some of these models, fewer than 100 distinct states ended up being assigned to positions in the genome.

### Computing fine-mapped variant enrichment

To compute enrichment of full-stack states for phenotypically associated fine-mapped variants, we downloaded data on fine-mapped variants for 3052 traits from CAUSALdb [66]. Specifically we obtained posterior probabilities of variants being causal based on two fine-mapping methods, FINEMAP [68] and CAVIAR [67], which do not use epigenomic annotations as part of the fine mapping procedure. For each method and trait combination, we separately partitioned the provided set of potential causal variants into distinct loci. To form the distinct loci, we merged neighboring variants into the same loci until there was at least 1MB-gap between the two closest variants from different loci. Separately for each fine-mapping method, trait, and locus combination, we selected the single variant with the highest posterior probability of being causal. For each fine-mapping method, we took the union of variants across 3052 traits, and then calculated the fold enrichments for the union of these lead variants with stacked ChromHMM states relative to the enrichment with a background set of common variants from dbSNP build 151 (hg19). To do this, we separately computed the enrichments of both of these sets relative to a genome-wide background, and then divided the enrichment of the foreground set (lead fine-mapped variants) by the enrichment of the background set (common variants). The dbSNP variants were obtained from the UCSC genome browser [81].

### Computing structural variant enrichments

To compute enrichment of the full-stack states for structural variant enrichments, we obtained data of structural variants from [47]. We used the B38 call set, which was in hg38 and used for the analysis presented in [47]. We filtered out structural variants that did not pass the quality control criteria of [47]. We then separately considered structural variants annotated as either a deletion or a duplication, for which there were, 112,328 and 28,962 sites respectively.

Since the structural variants were defined in hg38, we computed their enrichment for ChromHMM state annotations from full-stack and per-cell-type concatenated models that were lifted over from hg19 to hg38, following the procedure outlined above. Next, we followed the enrichment analysis procedure outlined above to compare full-stack vs. per-cell-type chromatin state annotations’ power in recovering structural variants.

To compare the power of full-stack state annotations vs. per-cell-type state annotation frequency, we utilized the 15-state per-cell-type chromatin state annotation for 127 cell types (reference epigenomes) from Roadmap Epigenomics Consortium. We followed the analysis outlined in [47], for each of the 15 per-cell-type states, we annotated genomic positions based on the number of cell types in which the state is present (ranging from 0 to 127), resulting in 15 state frequency annotations per genomic position. We then applied the procedure above for each per-cell-type state to compare the predictive power of the state’s annotation frequency against the full-stack annotation.

### Computing enrichments with cancer-associated variants

We obtained data of somatic mutations associated with different types of cancer from COSMIC non-coding variants dataset v.88 in hg38 [69]. We selected from this dataset variants that were from whole-genome sequencing. We filtered out variants that overlap with any of the following: the hg38 black-listed regions from the ENCODE Data Analysis Center (DAC) [82], hg38 dbSNP (v151) set of common variants from the UCSC genome browser database, or regions annotated as coding sequence (‘CDS’) based on GENCODE v.30 hg38 [83] gene annotations. We decided to restrict this analysis to the four cancer types with most number of variants present in the dataset in hg38: liver (1,351,417), pancreas (500,930), haematopoietic and lymphoid tissue (354,501), and breast (323,751). We then lifted over these sets of variants from hg38 to hg19, resulting in 1,351,159, 500,798, 354,351, and 323,685, variants respectively. To obtain a background set of genomic locations for the enrichment analysis, we filtered from the genome the same set of hg38 annotations of black-listed regions, common variants, and coding sequences as we did for the foreground of COSMIC mutations. We then lifted over these remaining positions from hg38 to hg19 to obtain the background. We calculated the enrichment of chromatin states with cancer-associated variants by first calculating the enrichment values of chromatin states with filtered variants associated with each of the four cancer types, and the enrichment values with background set of genomic bases, all relative to the whole genome. We then divided the cancer-associated variant enrichment values by the background bases enrichments.

### Gene ontology enrichments

We calculated the gene ontology enrichments of genes being in proximity to each full-stack state annotation using GREAT [84]. For each full-stack state, we reported the top-GO terms with lowest FDR-corrected p-values. If multiple GO terms showed the same minimum p-values, we reported all of them. A full list of GO-terms associated with full-stack states are available in **Supplementary Data File 1**.

### External annotations sources

The sources for external annotations for enrichments analyses, not given above, were as follows (all download links are listed in **Supplementary Data File 5**):

⍰ Annotations of CpG islands, exon, gene bodies (exons and introns), transcription start (TSS), and transcription end sites (TES), 2kb windows surrounding TSSs (TSS2kb) in hg19 and hg38 were RefSeq annotations included in ChromHMM (v1.18) and originally based on annotations obtained from the UCSC genome browser on July 26th 2015.
⍰ Lamina associated domains were for human embryonic lung fibroblasts that were included in ChromHMM (1.18), which were lifted over to hg19 from hg18 positions originally provided by [38].
⍰ Annotations of assembly gaps in hg19 and hg38 were obtained from the UCSC genome browser and correspond to the Gap track.
⍰ Coordinates of zinc finger genes correspond to non-overlapping coordinates from GENCODE’s hg19 gene annotation, v30 [83]. ZNF named genes were those whose gene named contained ‘ZNF’. The list of C2H2-type genes were from https://www.genenames.org/.
⍰ Annotations of coding sequences in hg19 and hg38 correspond to coordinates of genes whose feature type is ‘CDS’ from GENCODE’s hg19 and hg38 gene annotation, v30 [83].
⍰ Annotations of pseudogenes in hg19 and hg38 correspond to coordinates of genes whose gene type or transcript type contained ‘pseudogene’ from GENCODE’s hg19 and hg38 gene annotation, v30 [83].
⍰ Annotations of repeat elements were obtained from UCSC genome browser RepeatMasker hg19 tracks.
⍰ Per-cell-type concatenated ChromHMM chromatin state annotations were obtained from the Roadmap Epigenomics Consortium through http://compbio.mit.edu/roadmap [16]. These include data of the 18-state models based on observed data and the 25-state chromatin model based on imputed data for 98 and 127 reference epigenomes, respectively.
⍰ CTCF- per-cell-type chromatin states were based on the ChromHMM chromatin state annotations for six human cell types (GM12878, H1ESC, Helas3, Hepg2, Huvec, K562) for a 25-state model from the ENCODE integrative analysis [22, 31]. We extracted coordinates of regions annotated to the ‘Ctcf’ and ‘CtcfO’, both associated with CTCF signal and limited histone mark signal.
⍰ Blacklisted regions were those provided by the ENCODE Data Analysis Center (DAC) for hg19 and hg38 [82].
⍰ ConsHMM conservation state annotations for human (hg19) were those from [46].
⍰ Annotations of human genetic variants and their allele frequency were from GNOMAD v2.1.1 [62]. The dataset includes 229 million SNVs and 33 million indels from 15,708 genomes of unrelated individuals, which are aligned against the GRCg37/hg19 reference.
⍰ GWAS catalog variants were obtained from the NHGRI-EBI Catalog, accessed on December 5th, 2016 [63].
⍰ Coordinates of CpG sites profiled across cell types were obtained from DNA Methylation data in Roadmap Epigenomic portal.
⍰ Data of G/C content at 5bp resolution from UCSC Genome Browser, file hg19.gc5Base.txt.gz.
⍰ Data of binding regions of proteins of the polycomb repressive complexes were downloaded from the ENCODE portal [85]. Download links are listed in **Supplementary Data File 5.**

### Analysis of gene expression across states

To analyze the relationship between gene expression and the full-stack states, we downloaded gene expression data from the Roadmap Epigenomics Consortium [16]. Specifically, we downloaded a matrix of gene expression values, in RPKM (Reads Per Kilobase Million), for protein coding genes for 56 reference epigenomes that were among the 127 used as part of the full-stack model. In total, we obtained expression values for 19,795 Ensembl protein coding genes.

The gene expression data was obtained from (https://egg2.wustl.edu/roadmap/data/byDataType/rna/expression/57epigenomes.exon.RPKM.pc.gz). We also obtained the corresponding genomic coordinates for these genes from (https://egg2.wustl.edu/roadmap/data/byDataType/rna/expression/Ensembl_v65.Gencode_v10.ENSG.gene_info). For this analysis, we filtered out genes that are not classified as protein-coding. We transformed the gene expression values by adding a pseudo-count of 1 to the raw counts in RPKM, and taking the log of the resulting values.

For each full-stack-state and 56 reference epigenomes, we calculated the average gene expression of all genes overlapping with the state, taking into account the genes’ length. For each gene *g* we denote its length *L_g_* and expression *E_g_*. We let *s_i_* denote the state assigned at the 200-bp bin *i* and *g*, denote the set of genes overlapping the 200bp bin *i*. Let *B_s_* denote the set of 200bp bins that are assigned to state *s*. The average normalized expression with state *s* then becomes:

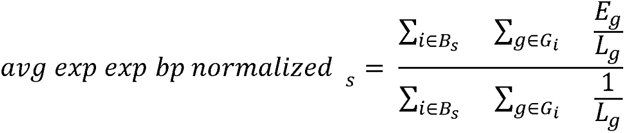

reference epigenomes. We used the BEDTools [86] *bedtools intersect* command to obtain the chromatin state assignments for 200bp segments that totally or partially overlap with any gene. To obtain average gene expressions of a state in a cell group as presented in **Fig. 3C**, we averaged the reported bp-normalized average gene expressions of the corresponding state across cell types within the group.

We also analyzed average gene expression values for each state as a function of the position of the state annotations relative to TSS, following a procedure similar to what was used previously [3]. We first identified a gene’s outer transcription start site (TSS) based on the reported coordinates of the gene and strand in the gene annotation file noted above. For each 200bp bin that is within 25kb upstream or downstream of an annotated TSS, including those that directly overlap with an annotated TSS, we determined the assigned full-stack state at this bin, and the position of the bin relative to those TSSs. Bins directly overlapping an annotated TSS were at position 0. If the gene was on the positive strand, the segments’ genomic coordinates lower than the TSSs’ correspond to upstream regions at negative points (minimum value: -250000), while genomic coordinates higher than the TSSs’ correspond to downstream regions at positive points (maximum value: 25000). If the gene is on the negative strand, the upstream and downstream positions are reversed. For each state and each 200-bp bin position relative to TSS, we determined the subset of genes where there is a 200bp bin annotated to that state at that position relative to their TSSs, and calculated their average expression. This produces a 100-by-251 table for one reference epigenome, corresponding to the number of full-stack states and 200-bp segments intersecting the 50kb windows surrounding genes’ TSSs and one segment directly overlapping the TSSs. We then smoothed the averaged expression data spatially by applying a sliding window with a window size of 21, i.e. each segment’s smoothed gene expression is the average of data in that segment and 21 surrounding genomic segments. Data of average gene expression in the first and last 10 segments within the 50kb window are not included in the window of smoothed data. We averaged results of 56 tables corresponding to 56 reference epigenomes as the final output from this procedure.

### Computing average DNA methylation levels

The DNA methylation analysis was conducted based on Whole Genome Bisulfite Sequencing data from Roadmap Epigenomics [16]. The fraction DNA methylation values was obtained from https://egg2.wustl.edu/roadmap/data/byDataType/dnamethylation/WGBS/FractionalMethylation.tar.gz. For each combination of 37 reference epigenomes with DNA methylation available and 100-full-stack states, the average fractional DNA methylation in that reference epigenome was computed for all CpG bases with a non-missing DNA methylation value overlapping the full-stack state annotation.

### Computing enrichment for bases prioritized by variant prioritization scores

To compute state enrichments for bases prioritized by different variant prioritization scores, we followed the approach of [46]. We obtained coordinates of bases containing prioritized variants based on 14 different methods as processed and described in [46]. The scores were Eigen and Eigen-PC version 1.1, funSeq2 version 2.1.6, and CADD v1.4, REMM, FIRE, fitCons, CDTS, LINSIGHT, FATHMM-XF, GERP++, phastCons, phyloP and DANN [48,50–61]. For 12 of the 14 scores, we separately considered prioritized variants genome-wide and in non-coding regions only. Two of the variant prioritization scores, LINSIGHT and FunSeq2, were defined only in the non-coding regions, so these scores were only used in the non-coding region analysis. As described in [46], the regions included in the non-coding analysis were defined as the bases where both LINSIGHT and FunSeq2 provided scores, which was 90.4% of the genome. For both the non-coding and whole genome analysis we computed the enrichment for bases ranked in the top 1%, 5% or 10% using the variant prioritization scores. We note that because of ties in some scores, the score-threshold above which we classified the bases as prioritized was chosen to be as close as possible to the target percentage (1%, 5% or 10%). We also note that if there were any bases with missing values for any particular score, then that base was assigned with the minimum values of such scores.

Enrichment values for the whole genome were computed as described above with the OverlapEnrichment command from ChromHMM. For computing enrichments restricted to non-coding regions, we first calculated enrichment of the non-coding prioritized variants relative to the whole genome and the enrichment of non-coding regions as defined above relative to the whole genome. We then divided these two enrichment values to obtain the enrichment of prioritized non-coding variants within non-coding regions.

### Data availability

Full-stack chromatin state annotation of the genome is available at https://github.com/ernstlab/full_stack_ChromHMM_annotations. An updated version of ChromHMM is available at https://ernstlab.biolchem.ucla.edu/ChromHMM/. All links to download publicly available data for analyses in this paper are listed in **Supplementary Data File**.

## Supporting information

All supplementary table SD1-6

## Acknowledgements

We thank Adriana Arneson for helping us in collecting data of analyses involving prioritized variants, repeat classes and GWAS catalog variants. We thank Hector Corrada Bravo, Amin Haghani, and Steve Horvath, Caesar Li, and Ake Lu for helpful discussions related to the manuscript. We acknowledge funding from US National Institute of Health (DP1DA044371, U01MH105578, UH3NS104095); US National Science Foundation (1254200, 2125664); Kure-IT award from Kure It cancer research, a Rose Hills Innovator Award, and the UCLA Jonsson Comprehensive Cancer Center and Eli and Edythe Broad Center of Regenerative Medicine and Stem Cell Research Ablon Scholars Program.

## Ethics Declarations

The authors announce no conflicts of interests.

---

**Supplementary Figure 1:**
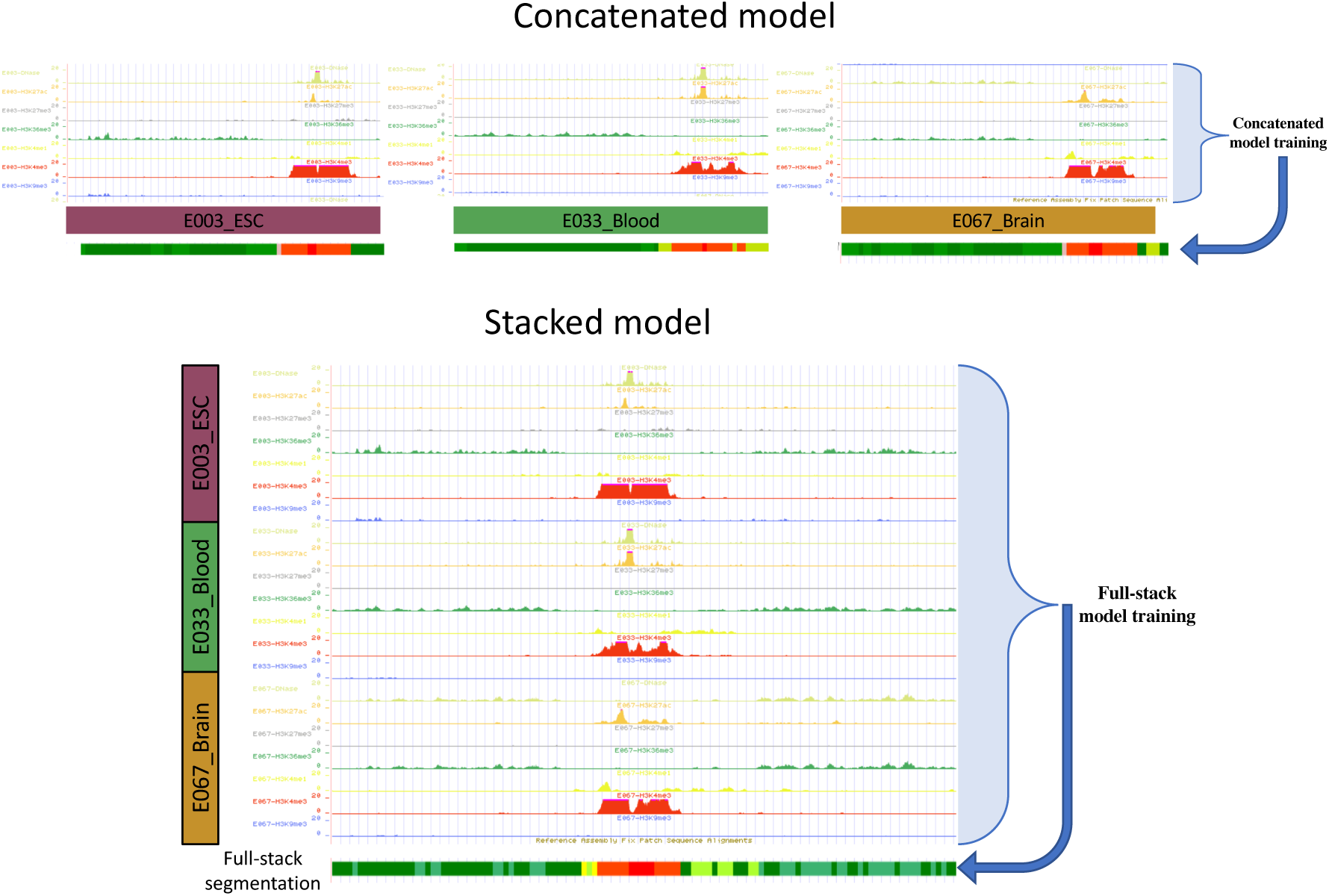
Illustration of concatenated model training vs. stacked model training. The top of the figure illustrates the concatenated modeling approach where a chromatin state annotation is produced for each cell type based on the data in that cell type using a common set of chromatin state definitions. In contrast, the stacked modeling approach produces a single chromatin annotation of the genome based on all the data.

**Supplementary Figure 2:**
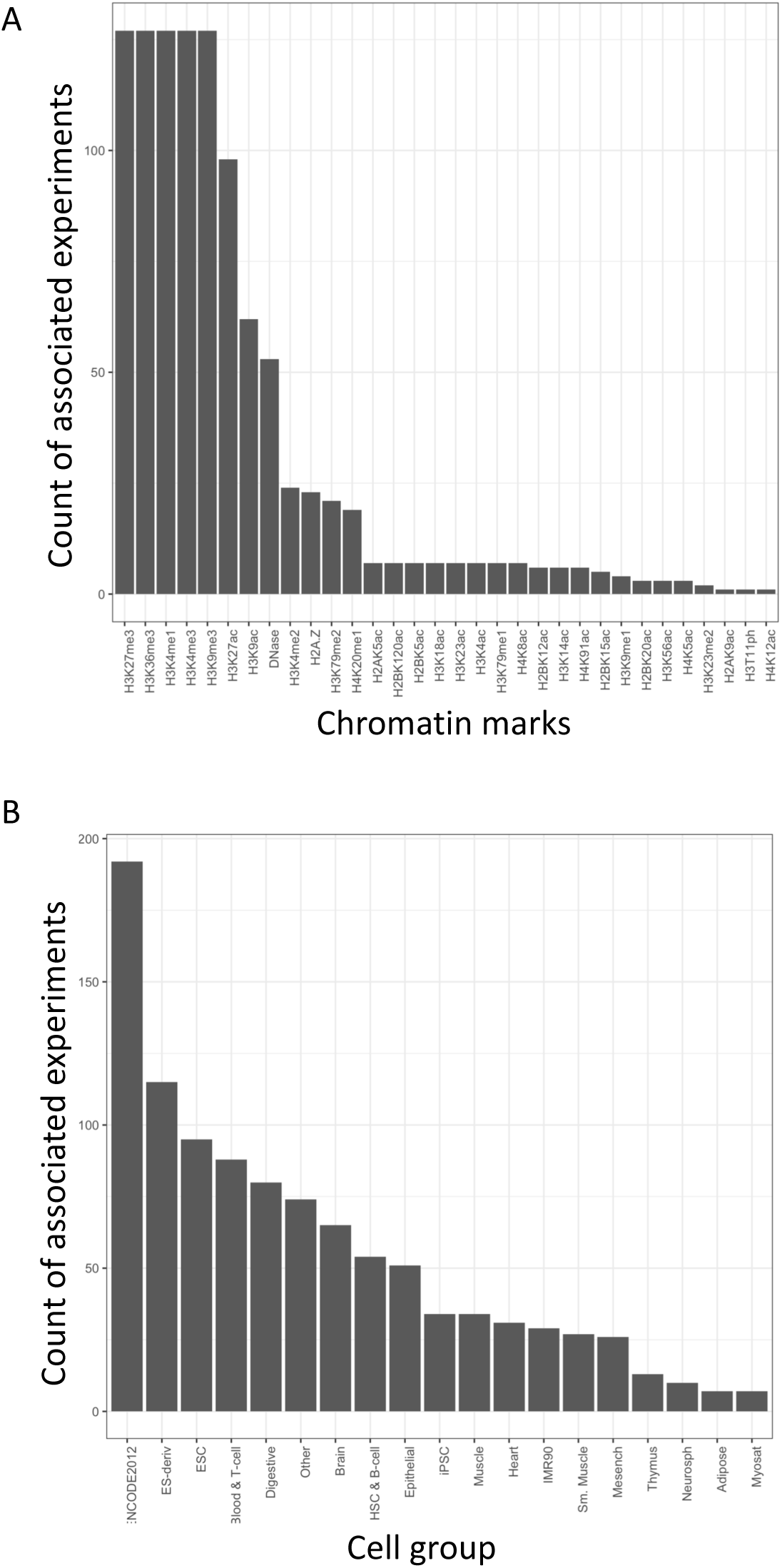
Mark and tissue group distribution of the input data tracks. **(A)** Counts of input tracks associated with different chromatin marks. There are five marks that were profiled in all 127 reference epigenomes, while some marks, largely acetylation marks, were profiled in few reference epigenomes. In total there were 1032 input tracks, including 53 DNase-seq experiments and 979 Chip-seq experiments. **(B)** Count of input tracks associated with different tissue groups previously defined (Kundaje et al., 2015).

**Supplementary Figure 3:**
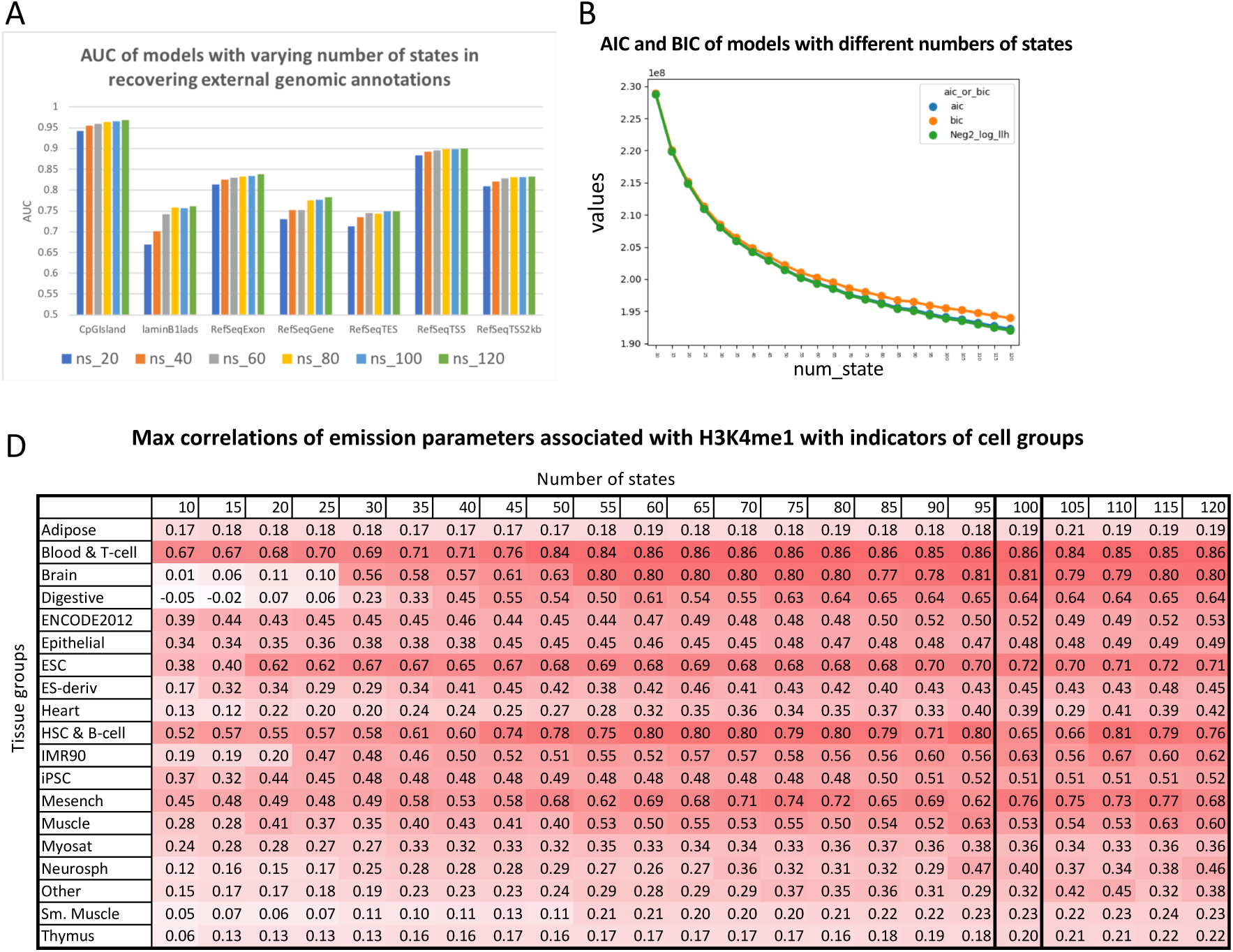
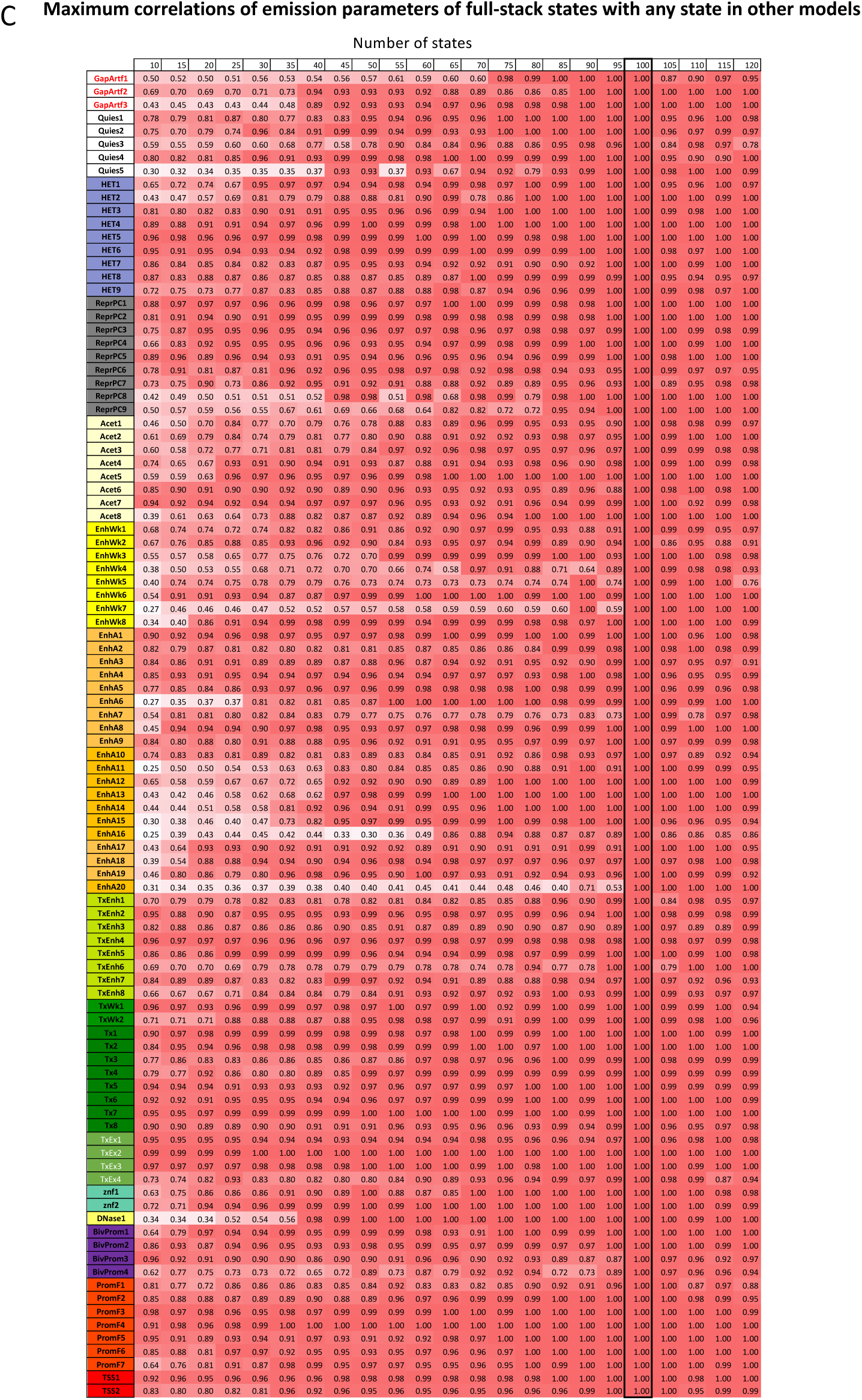
Evaluation of full-stack model’s number of states. **(A)** AUCs of full-stack models with varying number of states in recovering external genomic annotations. The figure shows the AUC of full-stack models with 20, 40, 60, 80, 100, and 120 states at predicting the genomic locations of multiple different external genomic annotations (CpG Islands, lamina associated domains (laminB1lads), Exon, Gene body, TES, TSS, and TSS2kb regions) (**Methods**). As the number of chromatin states increases, the AUC increases, but the level of the AUC increases diminishes. **(B)** The estimated AIC-BIC curves for models with the number of states ranging from 10 to 120 (5 states apart). We calculated the AIC and BIC based on ChromHMM’s output reporting the log-likelihood of observed data for 300 1-Mb regions. Neg2_log_llh: -2 * negative log likelihood of observed data. (**C**) Maximum correlations of emission parameters between each state in the 100-state model and any state for each other model. This is output from ChromHMM’s CompareModels command. Rows correspond to the states of the 100-state model. Columns correspond to models with varying numbers of states. Values are the maximum correlation of any state from the model in the column (with varying number of states) with the state from the 100-state in the row. The 100-state model is boxed. (**D**) Maximum correlations of emission parameters associated with H3K4me1 (an enhancer mark available in all cell types) and the binary vector indicating whether the cell type associated with an emission parameter is in a tissue group (1) or not (0). The rows correspond to different tissue groups from Roadmap Epigenomics Consortium (Roadmap Epigenomics Consortium et al, Nature 2015). The columns correspond to different models with varying numbers of states. The values show the maximum correlations mentioned above across all states within a model. The 100-state model is boxed.

**Supplementary Figure 4:**
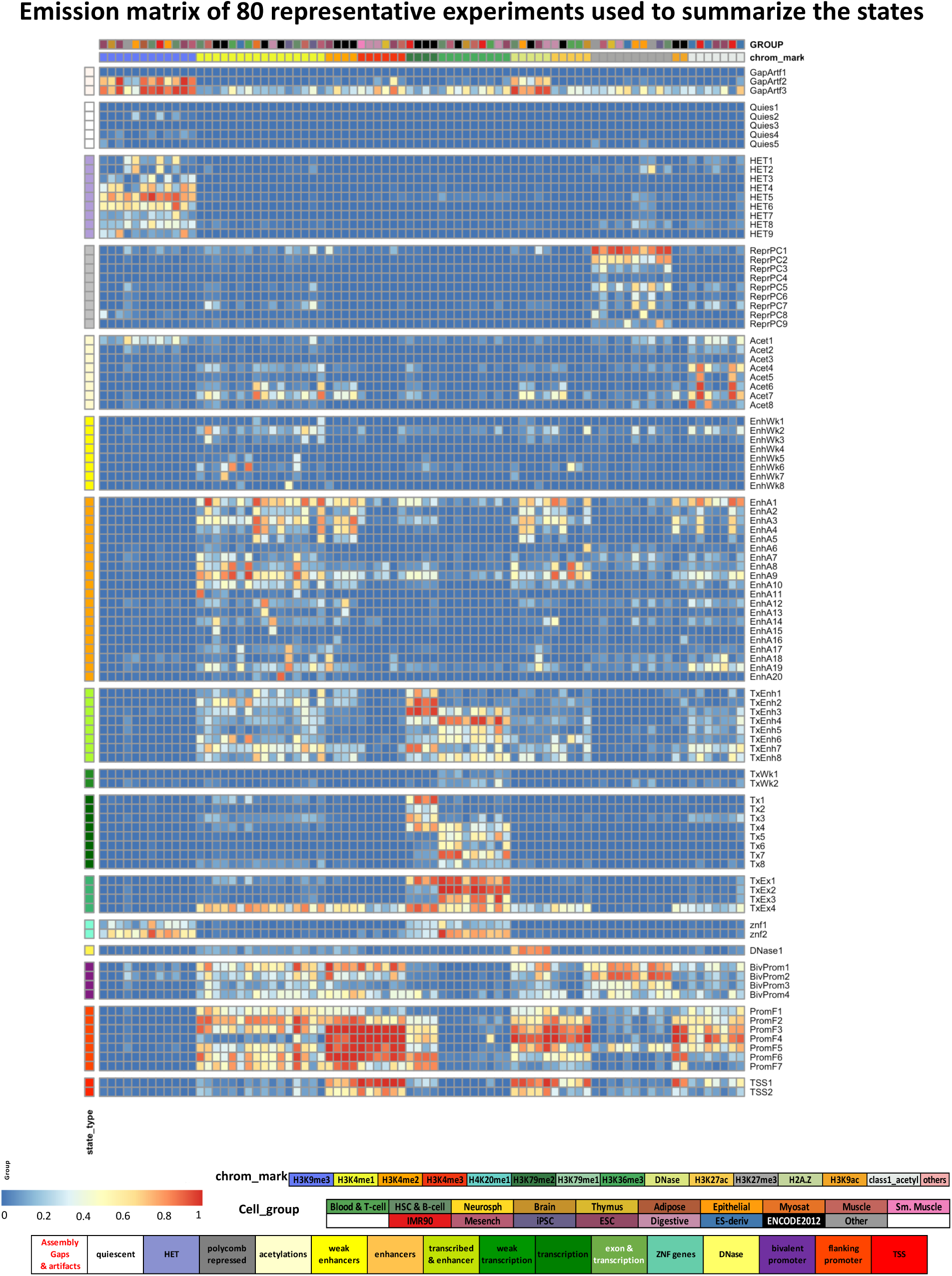
Emission probabilities of 80 experiments chosen to summarize the full-stack model. Each row in the heatmap corresponds to a full-stack state. Each of the 80 columns corresponds to an experiment that has been chosen to represent the space of 1032 experiments. These experiments were chosen through a greedy search of features that optimize prediction of the full-stack annotation using Naïve Bayes with the selected features (**Methods, Supplementary Methods**). For each state and each experiment, the heatmap gives the probability within the state of observing a binary present call for the experiment’s signal. States are displayed in 16 groups as in Fig. 2A. Color legends for the emission values, the state groups, chromatin mark, and tissue group are shown at the bottom.

**Supplementary Figure 5:**
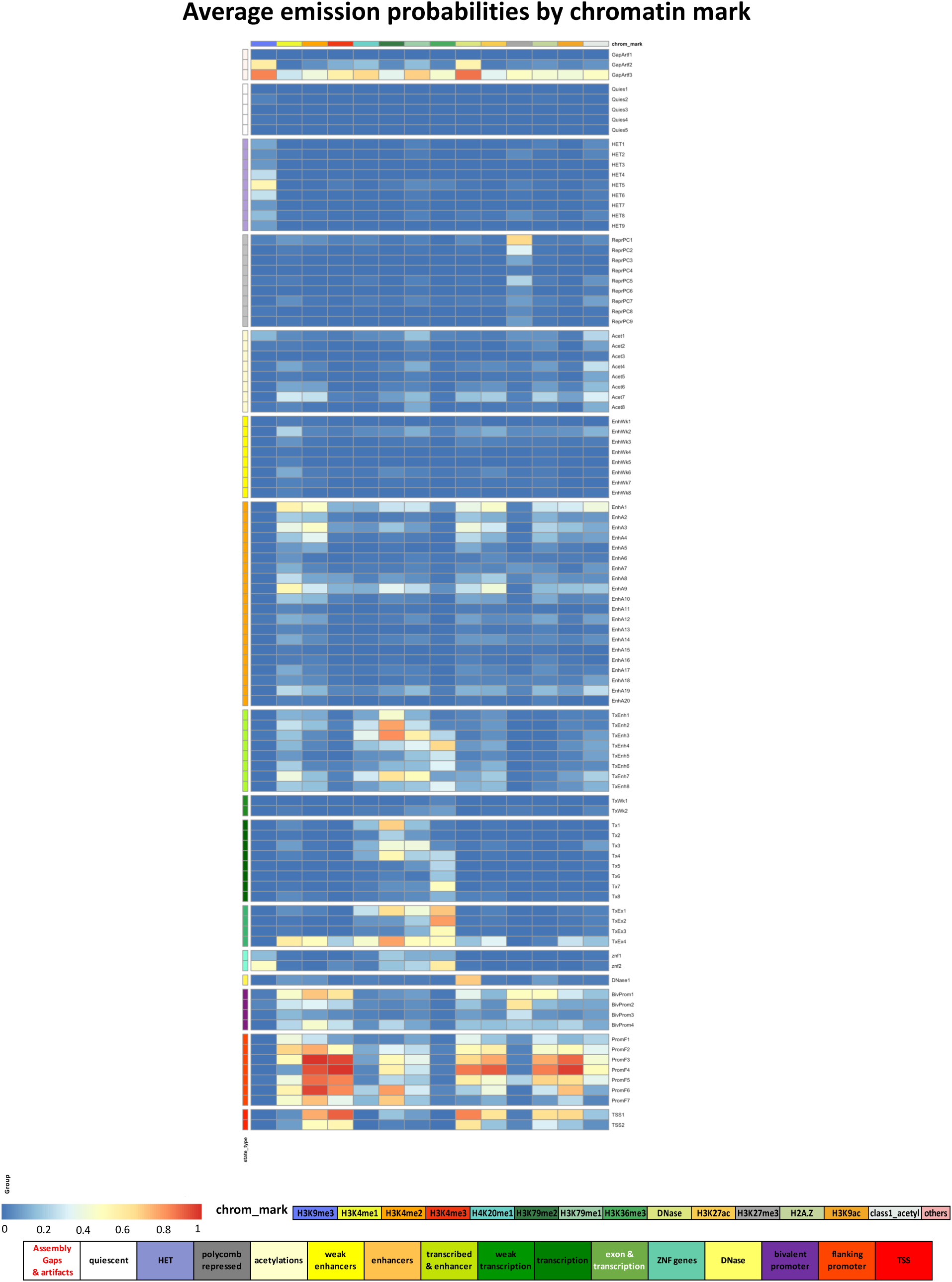
Full-stack states emission probabilities, averaged by chromatin marks. Each column corresponds to an individual chromatin mark or the group of acetylation marks. The heatmap shows for each state the average emission probabilities of experiments associated with each chromatin mark or with the group of acetylations. Color legends for the emission values, the state groups, and chromatin mark are shown at the bottom.

**Supplementary Figure 6:**
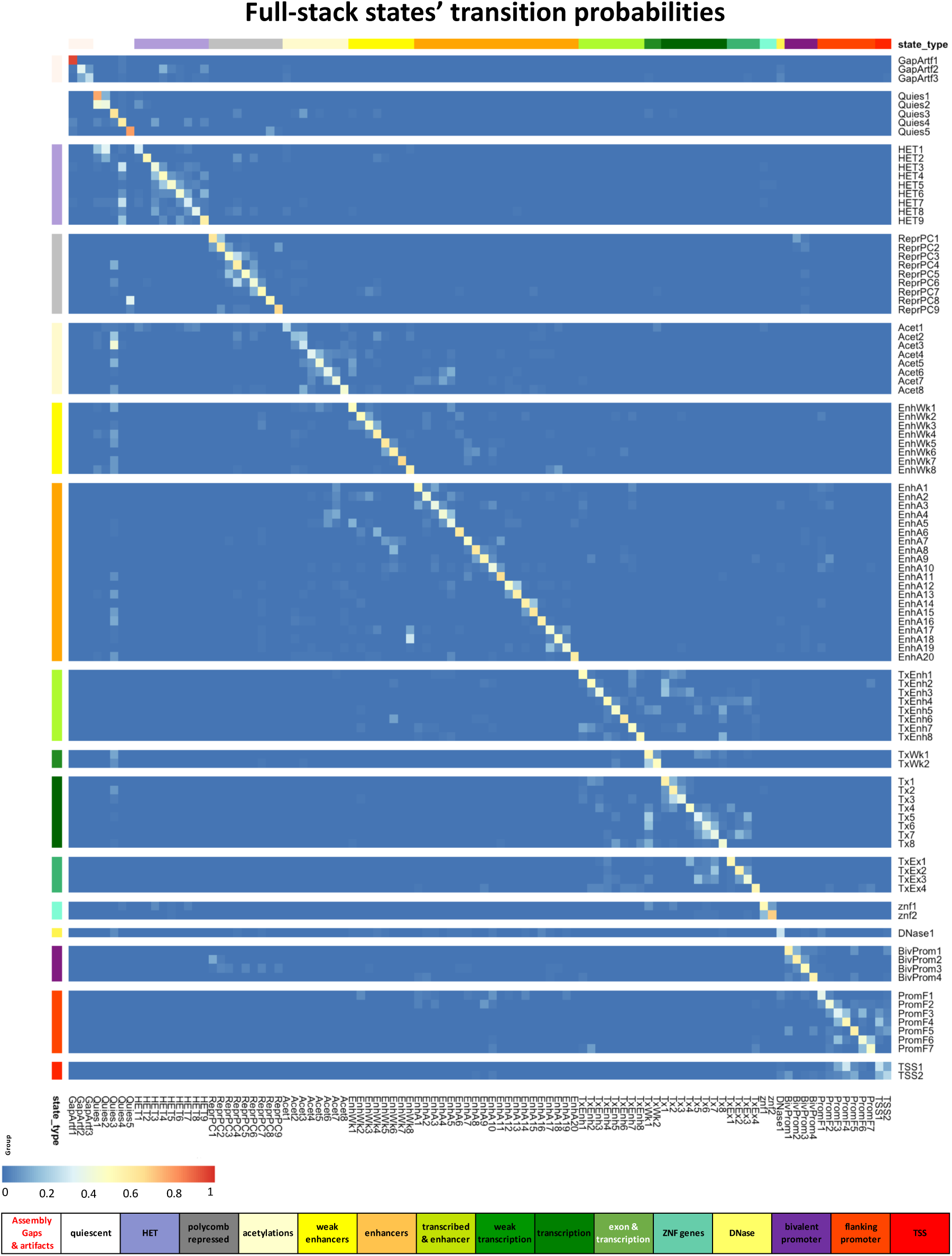
Full-stack states transition probabilities. Each row and each column correspond to a full-stack state, ordered based on their associated state group. The heatmap shows for each state assigned at a current genomic position (rows) the probabilities of transitioning to another state (columns) at the subsequent genomic position. Color legends for the emission values, the state groups are shown at the bottom.

**Supplementary Figure 7:**
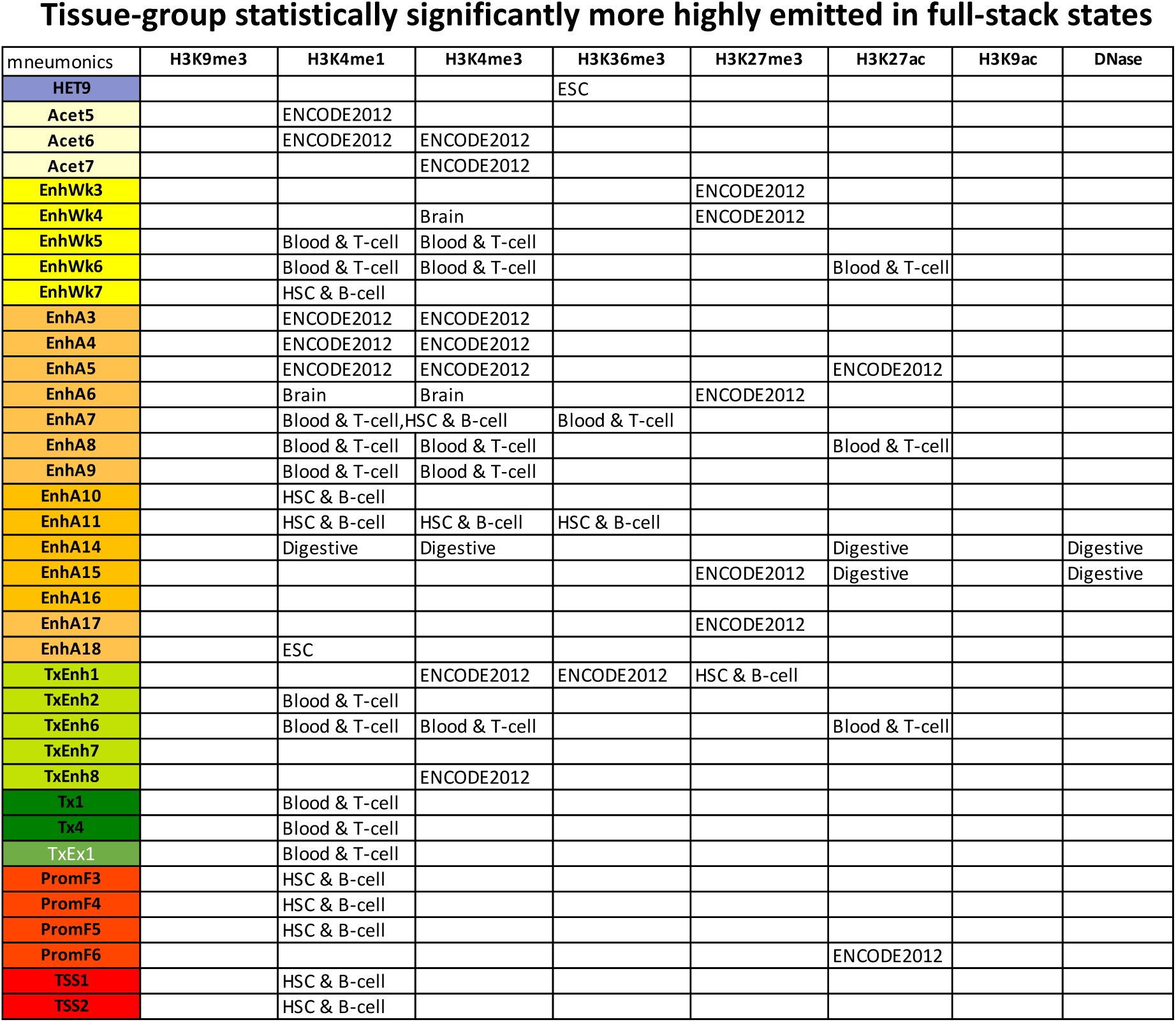
Statistically significant tissue—group specificity in full-stack states. The columns correspond to the eight most frequently profiled chromatin marks (H3K9me3, H3K4me1, H3K4me3, H3K36me3, H3K27me3, H3K27ac, H3K9ac, and DNase I hypersensitivity). The rows correspond to states that for at least one chromatin mark show statistically significant higher emission probabilities for one tissue group compared to others (**Methods**). Statistical significance is based on one-sided Mann-Whitney tests at a Bonferroni-corrected p-value threshold of 3.5e-6. The entries in the grid shows the tissue groups reaching significance for each chromatin mark-full-stack state combination.

**Supplementary Figure 8:**
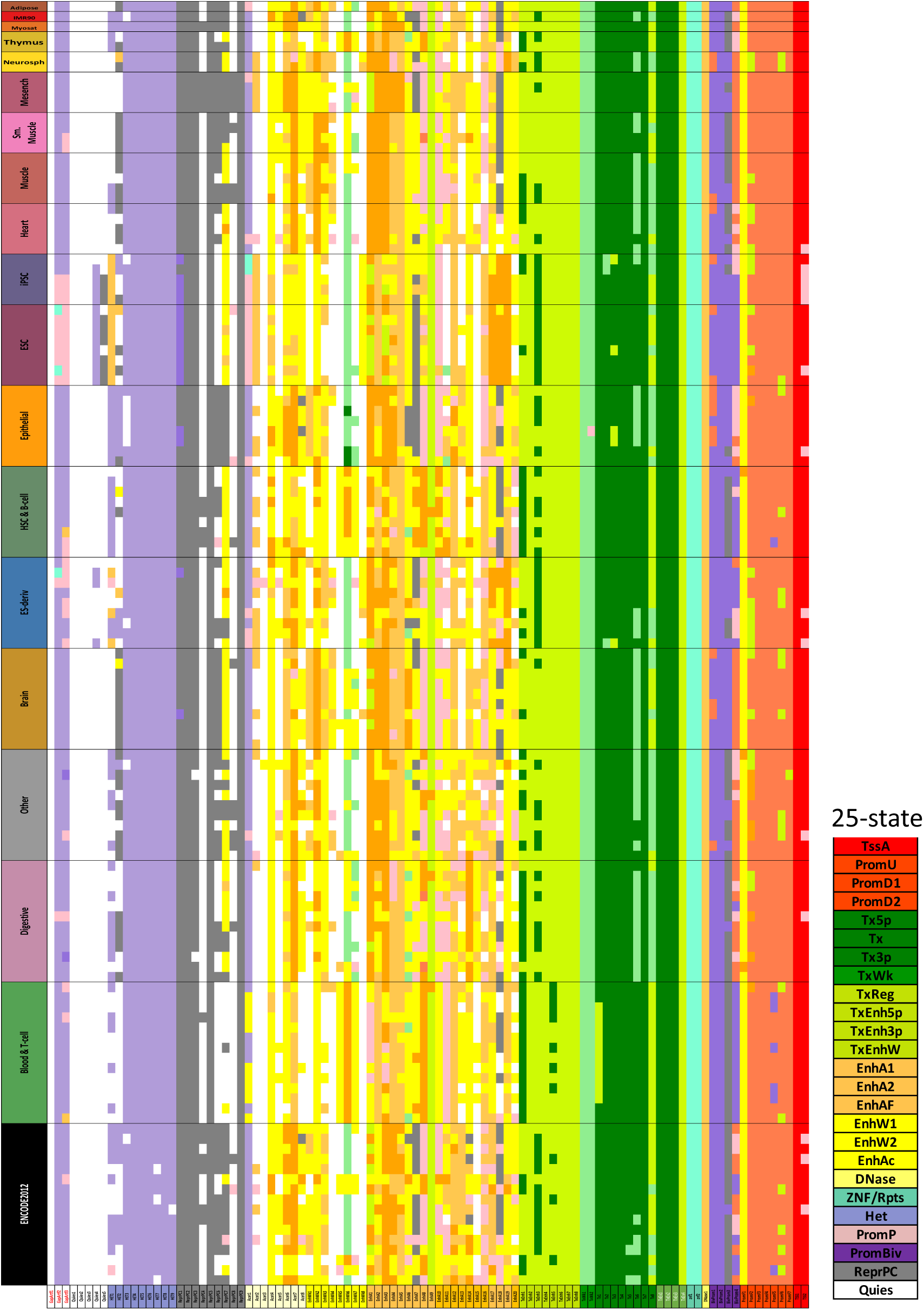
Full-stack states maximum-enrichments with annotated chromatin states in 127 reference epigenomes. Each row corresponds to one of 127 reference epigenomes from the Roadmap Epigenomics Consortium (**Methods**). Each column corresponds to a state of the full-stack model. Each color entry corresponds to a reference epigenome-full-stack state combination. The color corresponds to the chromatin state from the 25-state model annotating the respective reference epigenome that is most enriched with the respective full-stack state. The figure highlights how some full-stack states are maximally enriched with the same per-cell-type chromatin states across all the reference epigenomes; for example, states ZNF1 and ZNF2 are maximally enriched with ZNF Gene state in all 127 reference epigenomes’ 25-state per-cell-type annotation. At the same time, other full-stack states are enriched for distinct per-cell-type states, for example state EnhA8-- characterized as a blood enhancer state based on emission probabilities-- is most enriched with activate/flanking enhancer in cell types of the groups Blood&Tcell, HSC&B-cell, while being most enriched with poised promoter and weak enhancer states in other cell types. Detailed description of each full-stack state enrichment patterns with per-cell-type states can be found in **Supplementary Data 3**.

**Supplementary Figure 9:**
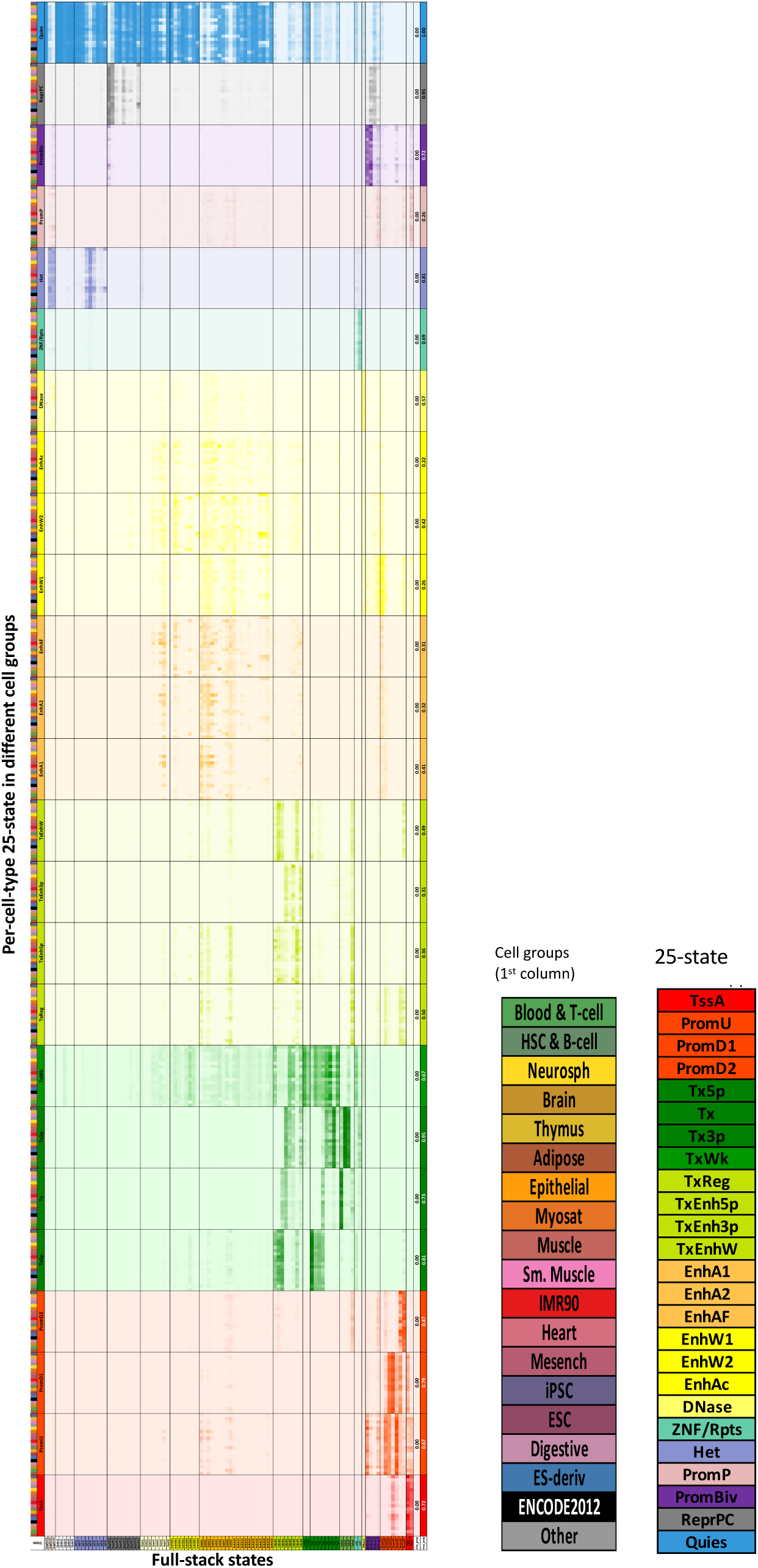
Estimated probabilities of per-cell-type chromatin states overlapping with full-stack states. The figure shows estimated probabilities of per-cell-type chromatin state assignments overlapping with full-stack state annotations conditioned on the cell group of the per-cell-type annotations. This figure is also provided as an excel file in **Supp. Data File 3** where it is accompanied with detailed comments about each full-stack state. The figure is based on a 25-state per-cell type chromatin state model (Ernst & Kellis, 2015), and 19-previously defined tissue groups for the 127 reference epigenomes (Kundaje et al., 2015). Each row corresponds to a combination of per-cell type state (among 25 states) and tissue group, as denoted in the first two columns and legend. Rows corresponding to the same per-cell-type state are grouped together. The first two columns show the colors of tissue groups and per-cell-type state, respectively, as indicated in legends on the right, and matching with the colors in **Supp. Fig. 8**, except we changed per-cell-type quiescent 25-state from white to blue for better visibility. The 100 following columns correspond to 100 full-stack states. Values in the heatmap correspond to the estimated probability a genomic position annotated as a full-stack state (column) is also annotated as a per-cell-type state in a cell type from the corresponding tissue group (row) (**Methods**). The last two columns show the minimum and maximum probabilities observed for each per-cell type state for any combination of tissue group and full-stack state. The heatmap colors correspond to the 25-state’s colors and are scaled such that the maximum probability values in each block are colored darkest (as seen in the right most column). The figure complements **Supp. Fig. 8** in providing information on how each full-stack state can correspond to different 25-per-cell type states in different groups of cells, hence stratifying full-stack states’ characteristics in more details. For example, full-stack state ReprPC8 shows high probabilities of overlapping ReprPC state in ESC-related cell groups (ESC, iPSC, and ES-derived), and quiescent state in other cell groups.

**Supplementary Figure 10:**
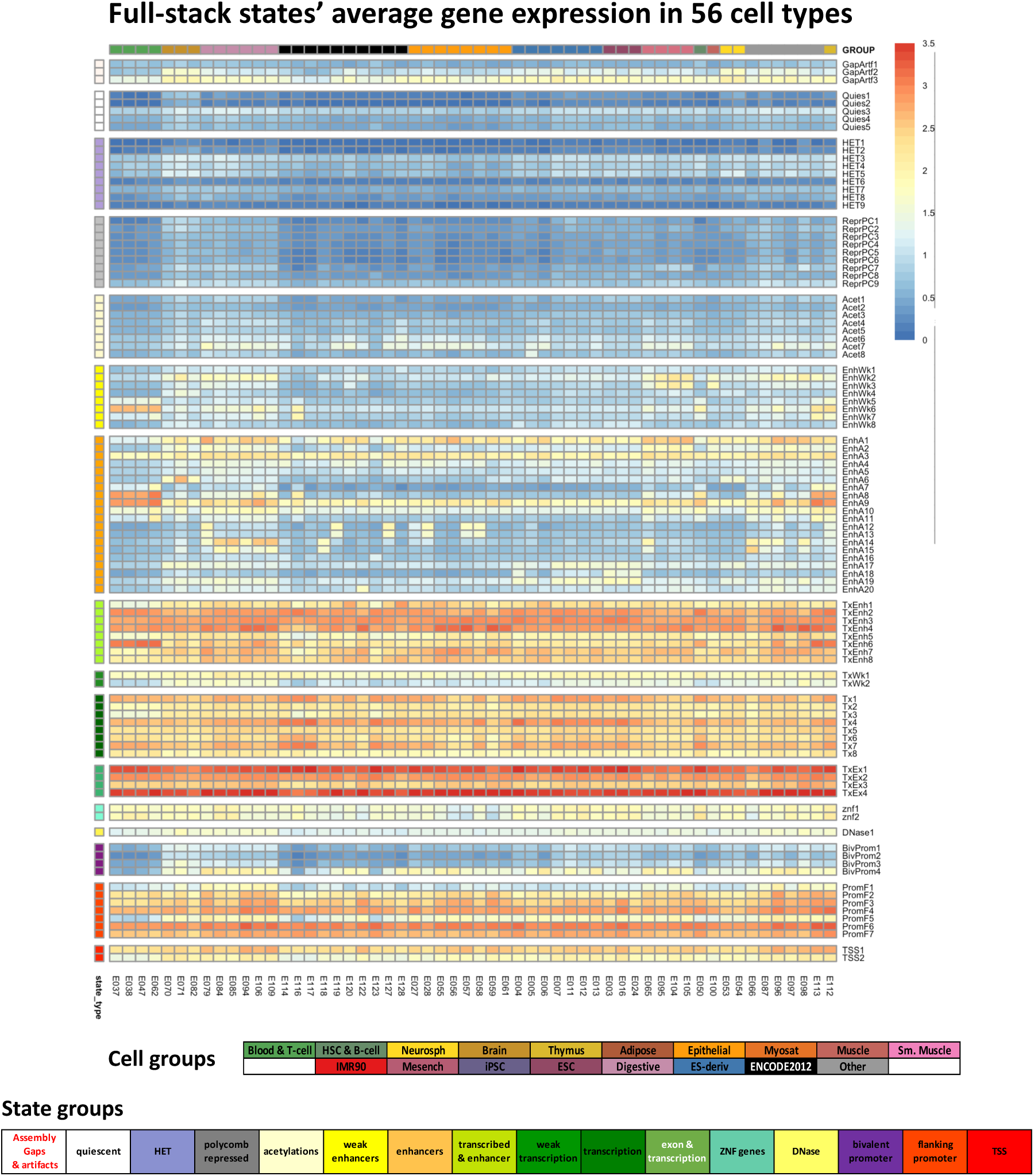
Full-stack states’ average gene expression in different cell types. Each row corresponds to one of the 100 full-stack states grouped into state groups as indicated by the legend at the bottom. Each column corresponds to one of 56 cell types whose gene expression data were available from Roadmap Epigenomics (Kundaje et al., 2015). The columns are grouped based on their associated tissue group as indicated by the legend at the bottom. Each column shows the average expression of genes in the respective cell type that overlap with each full-stack state, weighted by the extent of the overlap and the gene length (**Methods**). The figure highlights how states in the transcription and exon group show consistently high gene expression across all cell types, while cell-type-specific enhancer states tend to show higher gene expression in the cell types corresponding to those states.

**Supplementary Figure 11:**
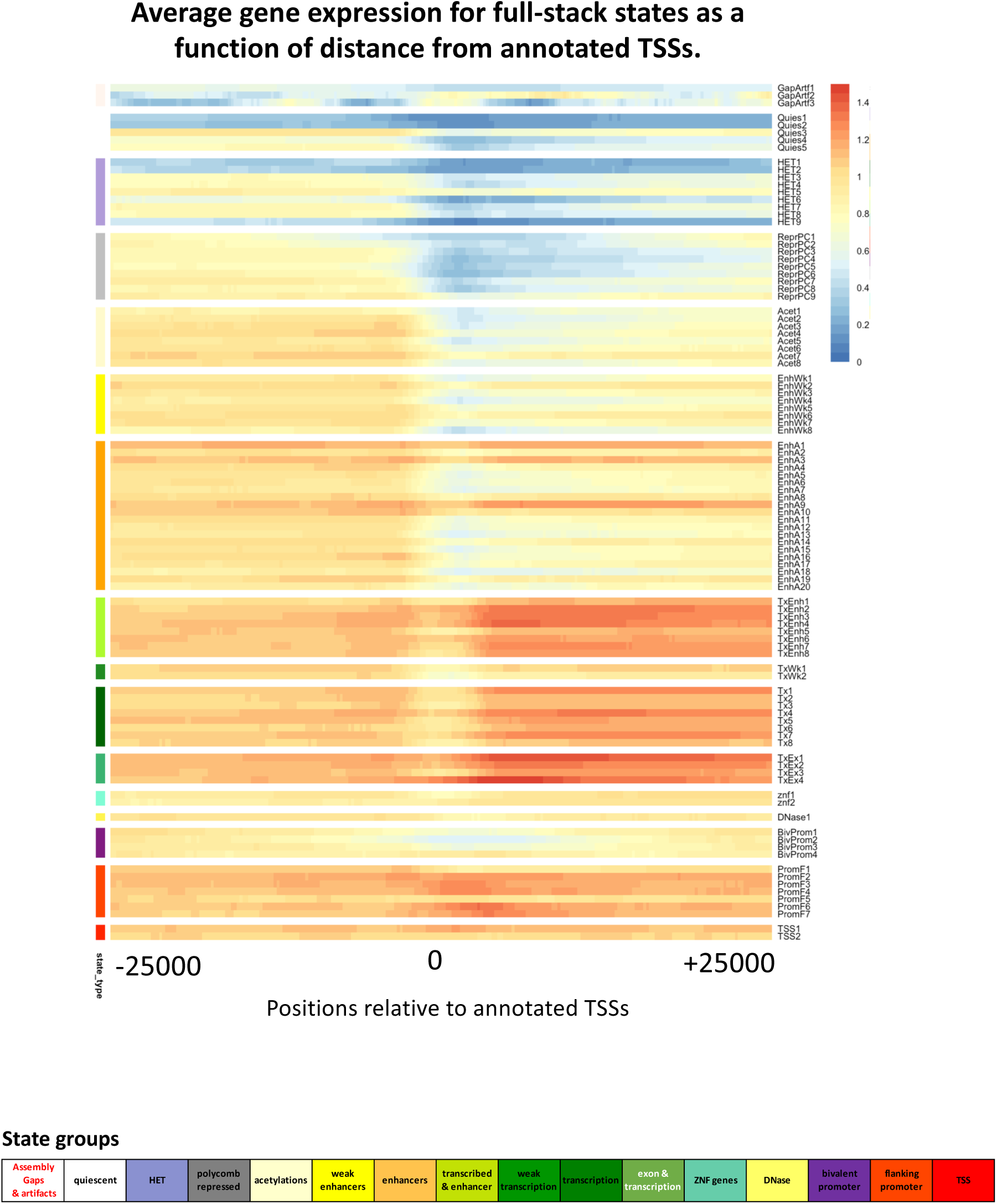
Full-stack states’ average gene expression as a function of distance from TSS. Each row corresponds to a full-stack state. Each column corresponds to a 200-bp bin within 25kb relative to annotated TSS, such that TSS is at position 0. Positions downstream of TSS in the direction of transcription have positive coordinate values, and those upstream have negative values. The heatmap shows for each state and position relative to the TSS, the average expression, across 56 cell types, of genes that have the state annotation at such position relative to the TSS (**Methods**). The figure highlights that states in the transcription group tend to have higher gene expression compared to other states, and the average gene expression is usually larger toward the downstream of genes. The figure also shows that for TSS and flanking promoter states, the average gene expression is relatively higher around the TSS compared to other positions.

**Supplementary Figure 12:**
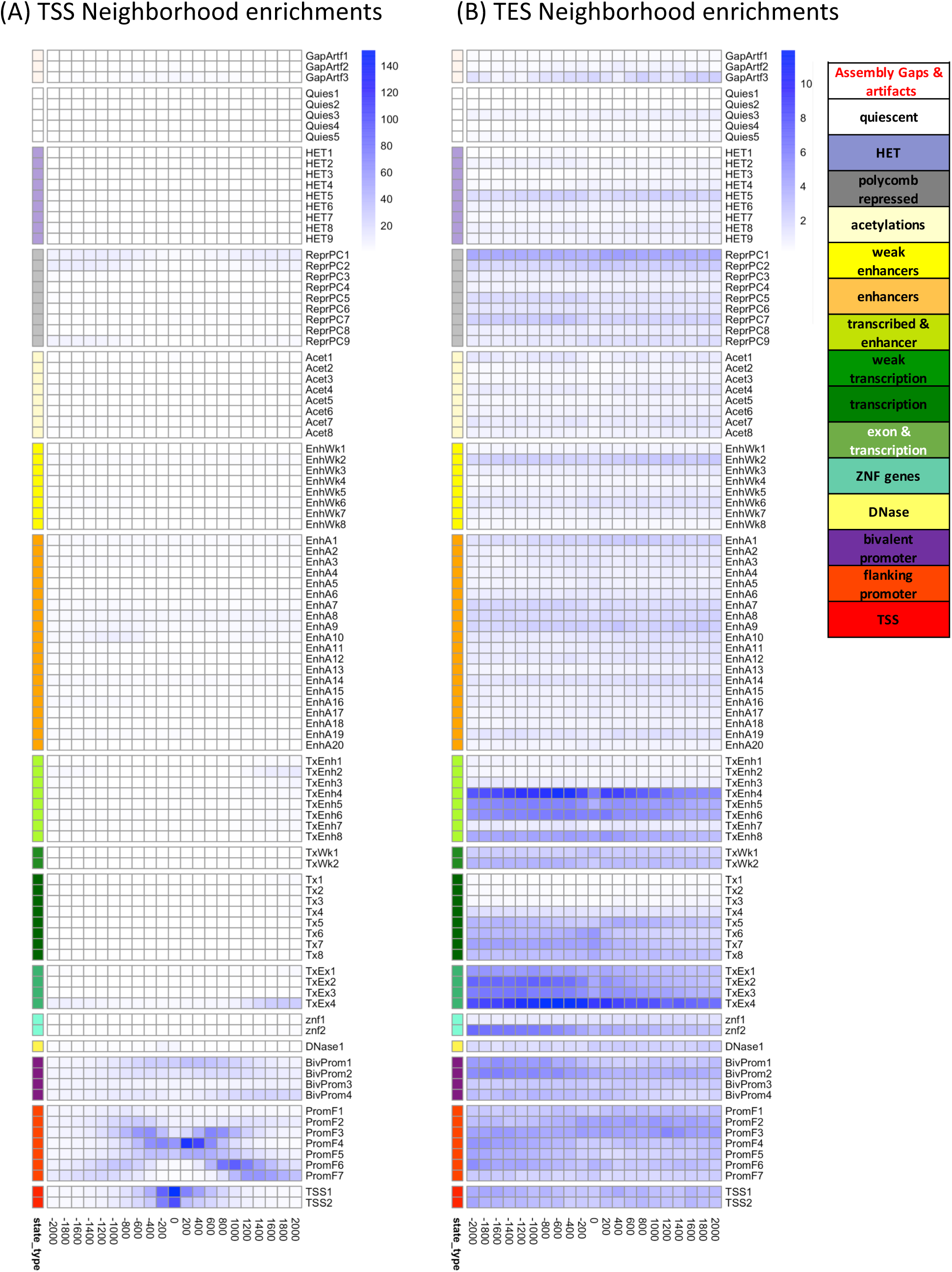
Positional enrichments of full-stack states. The figure shows positional fold enrichments for positions within 2kb of annotated **(a)** transcription start sites (TSS) and **(b)** transcription end sites (TES). Each column corresponds to one 200bp window as indicated at bottom. Positive coordinate values represent the number of bases downstream in the 5’ to 3’ direction of transcription, while negative values represent the number of bases upstream. Enrichments are calculated based on a genome-wide background. Color scale of enrichments is indicated at right for each panel. State groups’ color legends are shown at right.

**Supplementary Figure 13:**
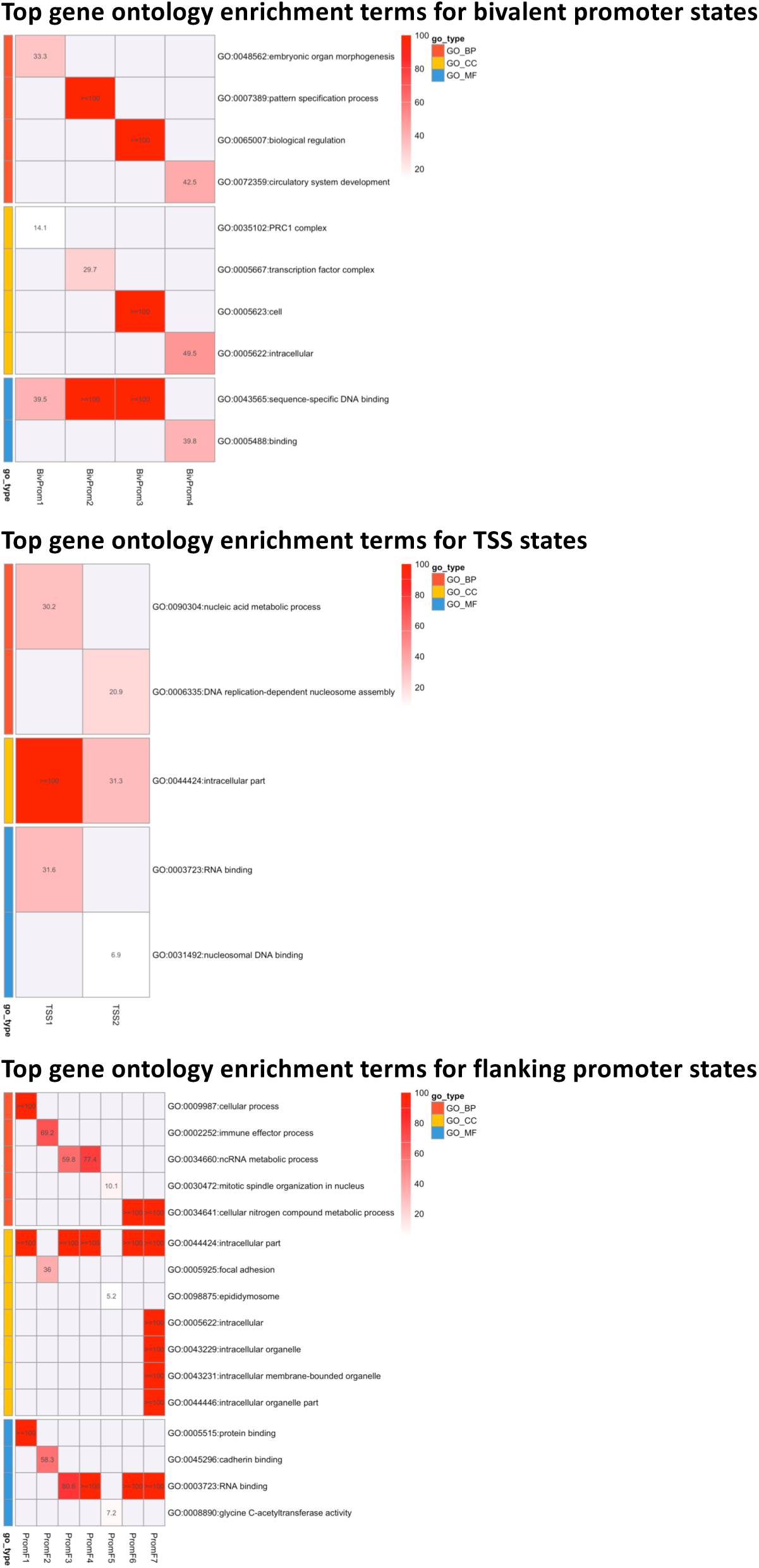
Top GO terms for states in promoter-associated states. In each subpanel, each column corresponds to a full-stack state, each row corresponds to a GO term, and each cells show the negative log of FDR-corrected p values of enrichments for states and GO terms. Only terms that are a top enriched GO terms with at least one full-stack states are shown. For each full-stack state, only the associated top GO term’s negative log q-values are shown and colored according to the color scales on the right. The most enriched GO- terms for all full-stack states are provided in **Supplementary Data File 1**.

**Supplementary Figure 14:**
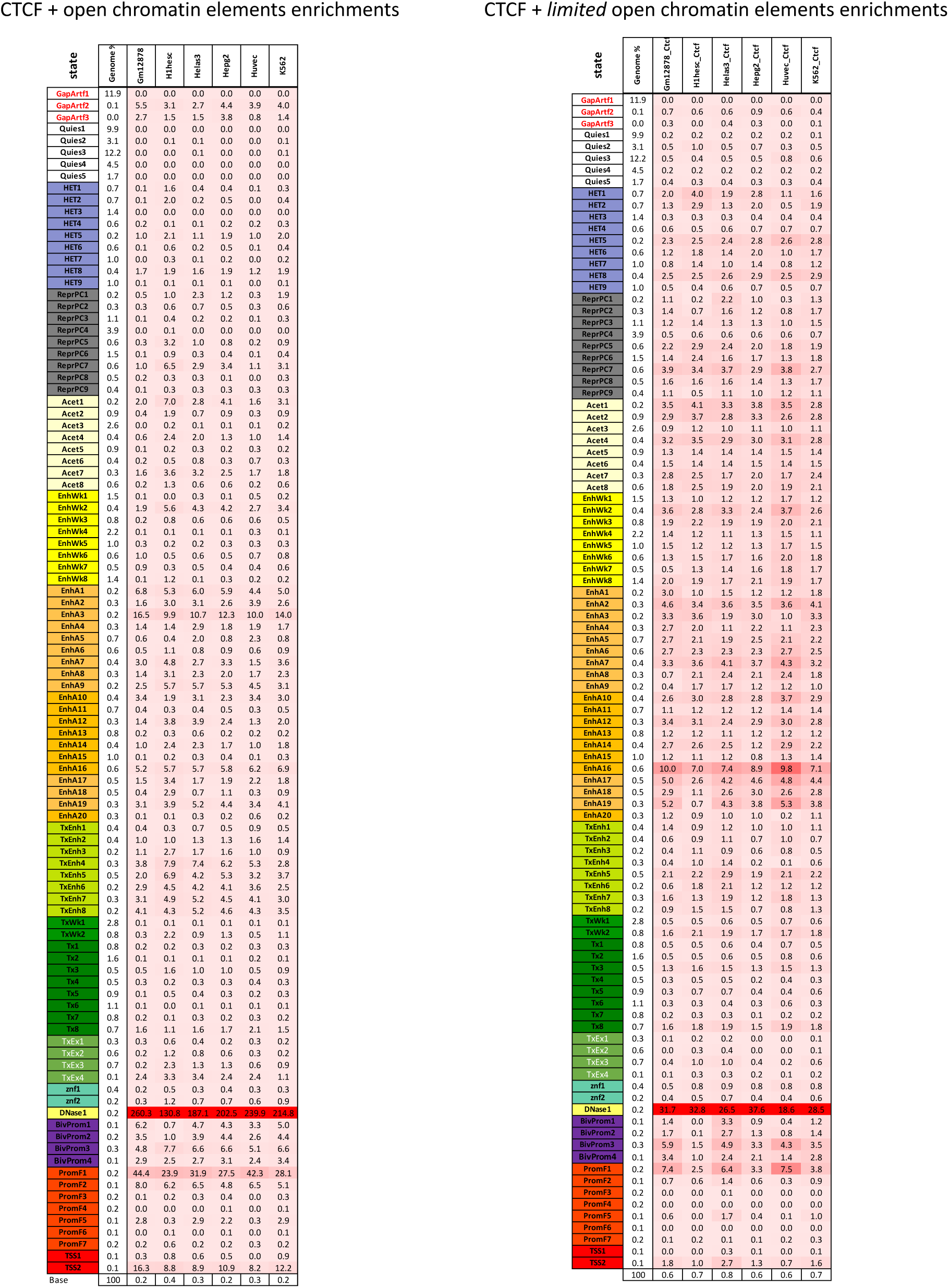
Full-stack states enrichments with CTCF associated chromatin states. **(A)** The heatmap shows enrichment values for the full-stack states (rows) and a chromatin state that corresponds to CTCF with open chromatin and limited histone modification signal from per-cell-type annotations in six different cell types (columns). CTCF signals were included as input for training these per-cell-type chromatin state models (**Methods**). Coloring of enrichments is column specific. **(B)** Similar to (A), except showing enrichments for a state associated with CTCF signal with limited open chromatin and limited histone modification signals.

**Supplementary Figure 15:**
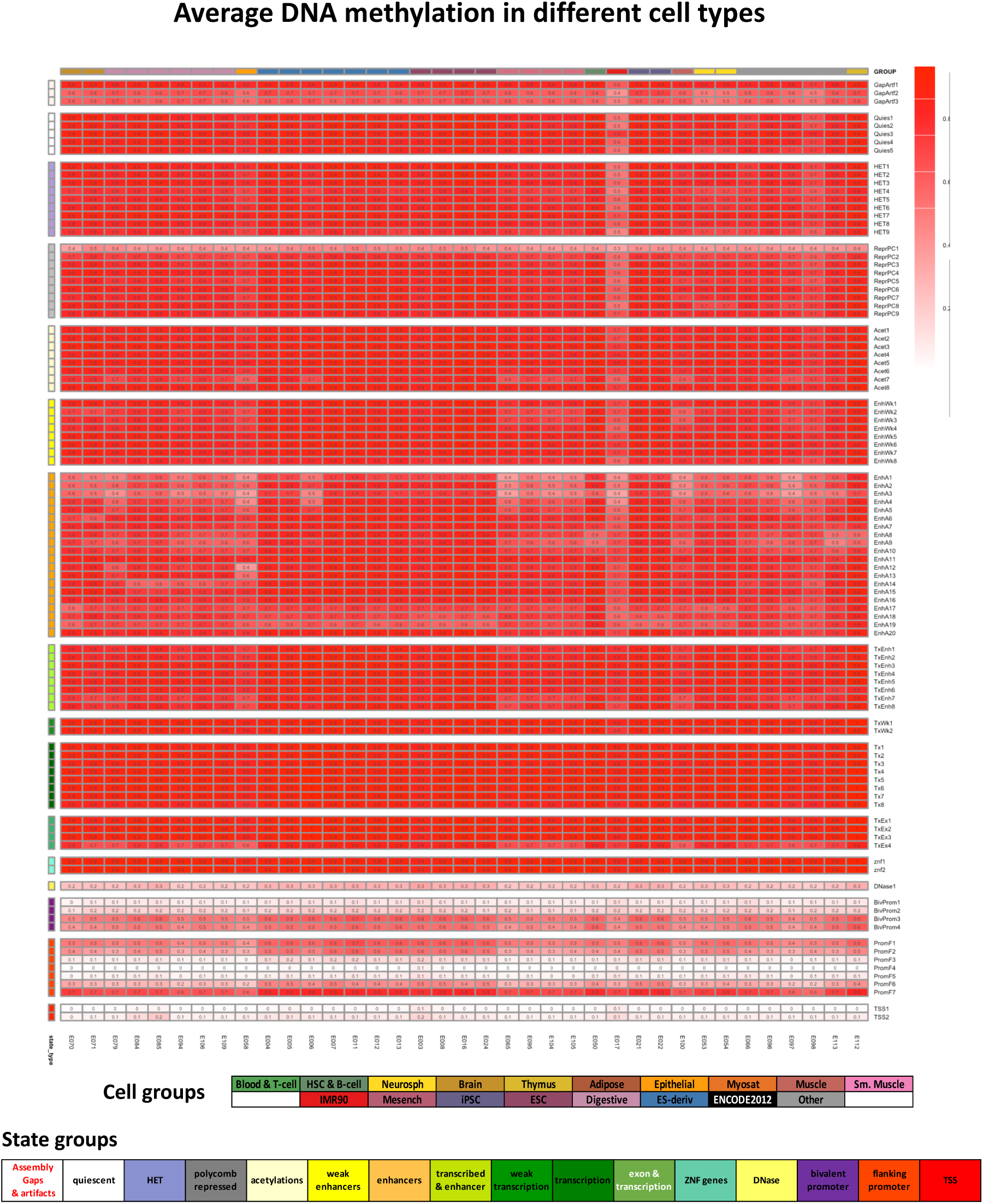
Full-stack states’ average DNA methylation in different cell types. Each row corresponds to one of the 100 full-stack states grouped into state groups as indicated by the legend at the bottom. Each column corresponds to one cell type whose DNA methylation data were available from Roadmap Epigenomics. The columns are grouped based on their associated tissue group as indicated by the legend at the bottom. Each column shows the average DNA methylation level in the respective cell type that overlaps with each full-stack state (**Methods**). Among promoters-associated states, those most enriched with CpG islands also show lowest average DNA methylation levels (Fig. 3A), consistent with expectation (Jones & Takai, 2001; Weber et al., 2007).

**Supplementary Figure 16:**
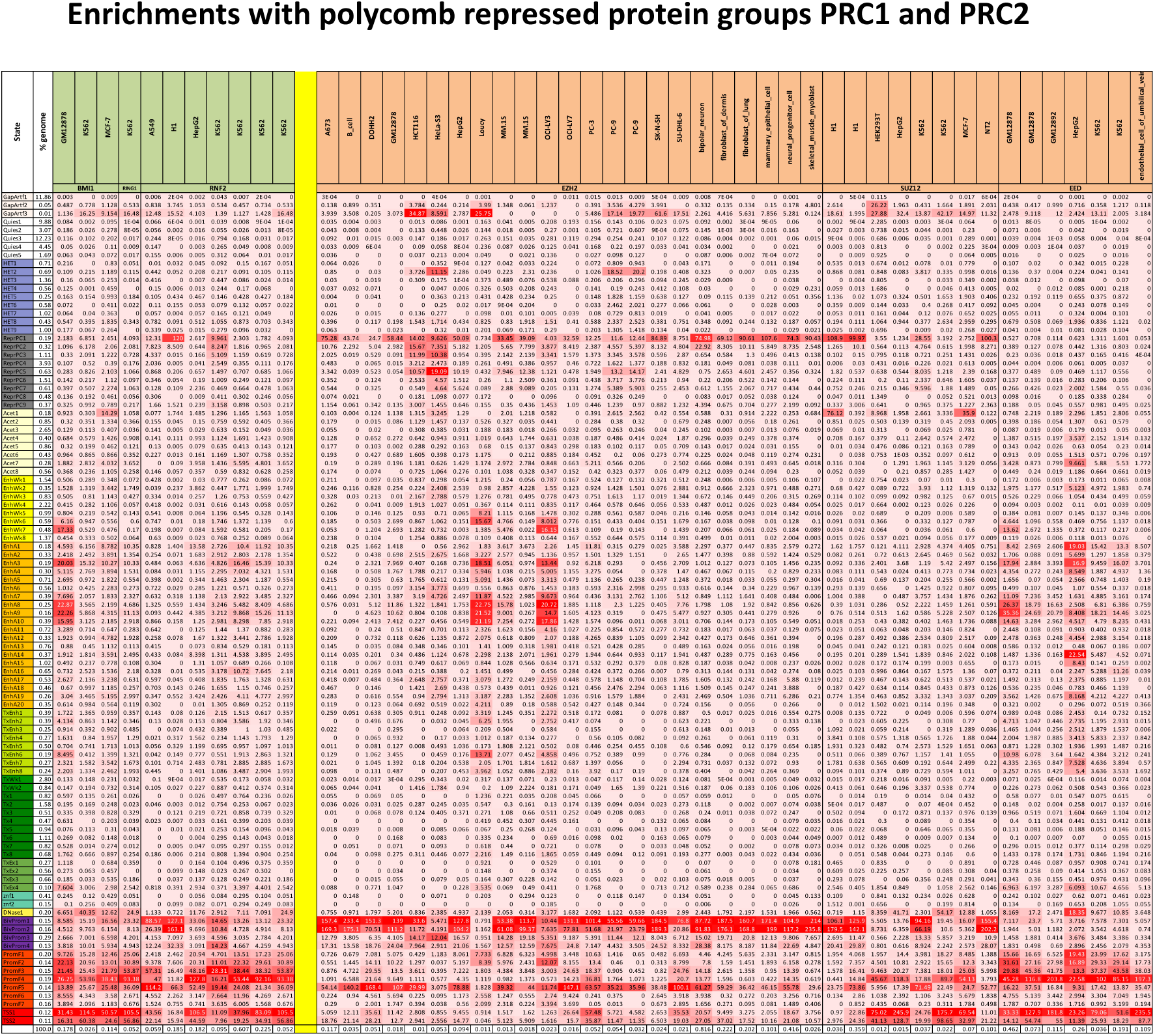
Full-stack states enrichments with Polycomb Repressive protein complexes PRC1 (green column headers) and PRC2 (orange column headers). The heatmap shows enrichment values for the full-stack states (rows) and the binding sites of subunits of PRC1 and PRC2 in different cell types (columns) that were available from ENCODE project (**Methods**). Coloring of enrichments is column specific with highest and lowest enrichment values in each column are colored red and white, respectively. The first two columns show state names, and their percentage of genome coverage. The last row shows the percentage of genome coverage for each type of PRCs.

**Supplementary Figure 17:**
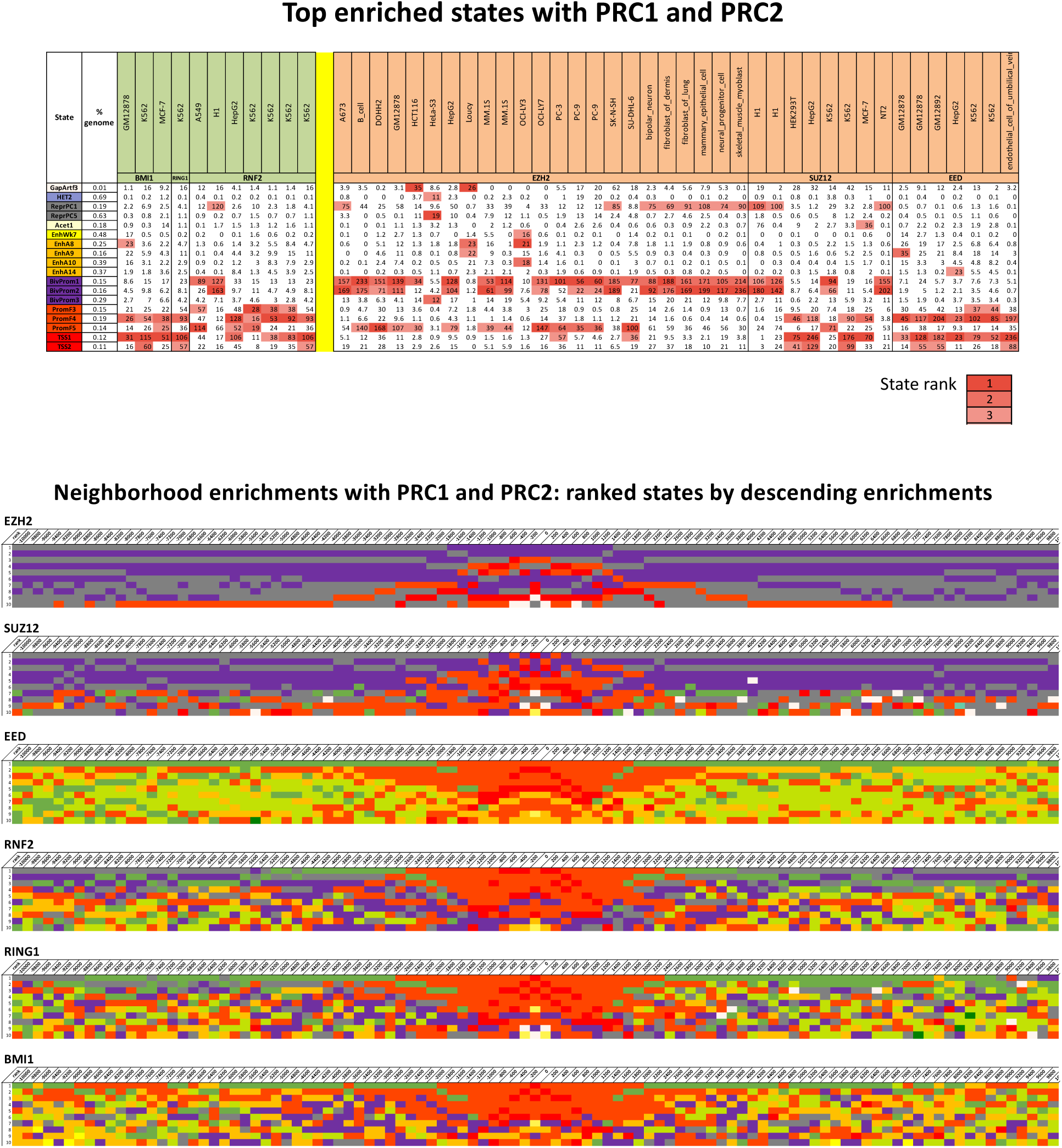
Enrichments of selected full-stack states with PRC1 and PRC2. **(A)** A subset of **Supp. Fig. 16** showing fold enrichment of full-stack states for binding sites of polycomb repressive protein complexes in different cell types from ENCODE (Methods). The column names highlighted in green (PRC1) and orange (PRC2) show the subunits of PRCs (BMI1, RING1 and RNF2 in PRC1, and EZH2, SUZ12, EED in PRC2) and the cell types where the PRCs were profiled in the second and first rows of column names, respectively. Each corresponding row corresponds to a state, and only states that were among the top three with greatest enrichments for at least one category of PRC complexes are shown. Top enrichment values are colored red based on the rank of the state for each score as indicated in the color legend at the bottom. Some states that show strong consensus across cell types in enrichments with PRC1 and PRC2 include ReprPC1, BivProm1-2, PromF4-5, TSS1-2. ReprPC1 and BivProm1-2 all show strong signals of H3K27me3. **(B**) Neighborhood enrichments of full-stack states with binding sites of PRC1 and PRC2 complexes. In each subpanel, each column corresponds to a 200-bp bin across the 20,000-bp regions overlapping and surrounding annotated PRC1&2 subunit complexes. Within each column, the top 10 states most enriched at the corresponding 200-bp position (within the 20,000bp window) are shown, in descending order of enrichments, and colored based on the state groups as presented throughout the paper. **Supp. Data. File 2** accompanies this figure to show full state names and rankings.

**Supplementary Figure 18:**
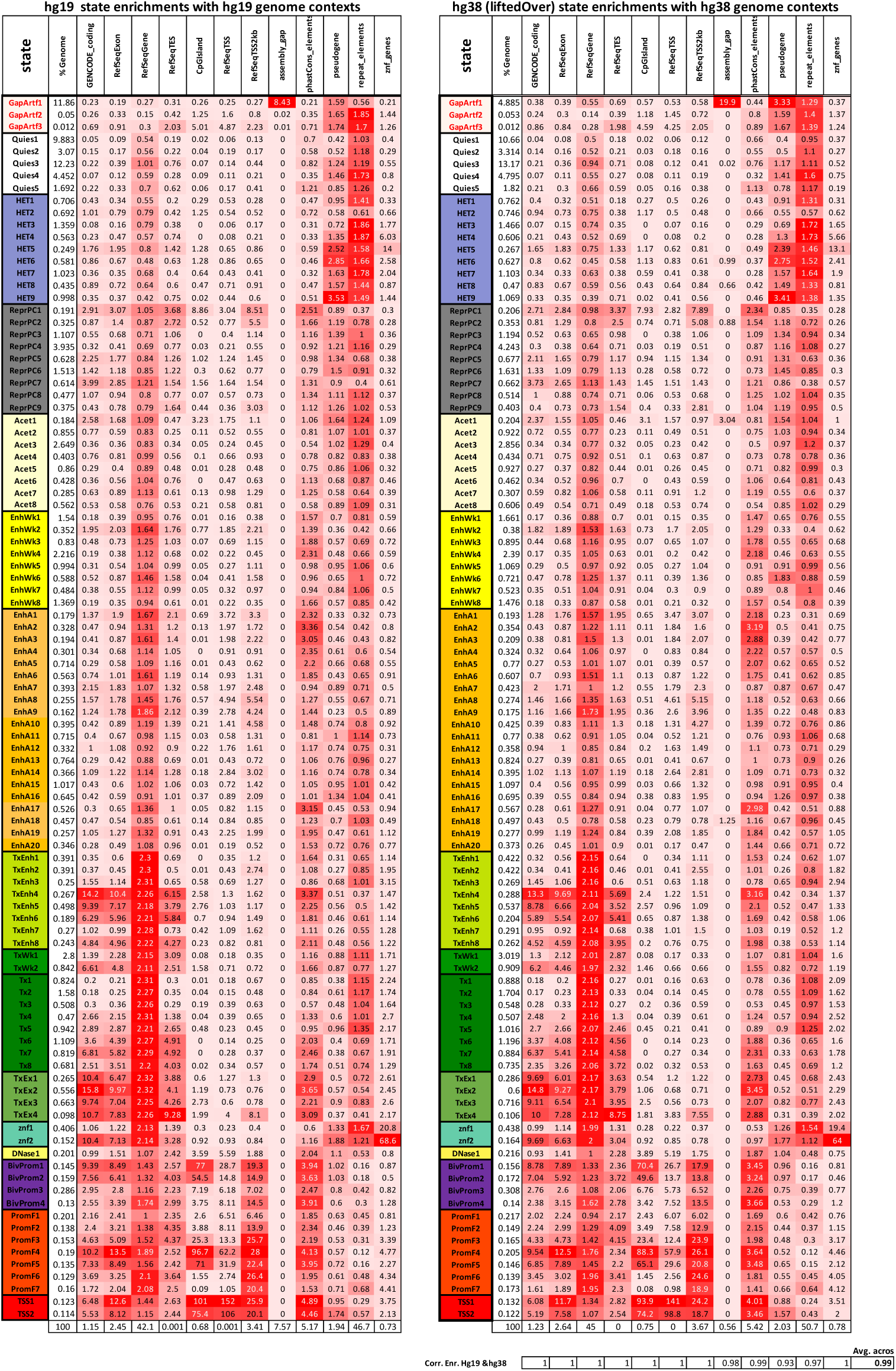
Comparison of hg19 and hg38 full-stack states enrichments with annotated genomic contexts. The heatmaps show enrichment values for the full-stack states (rows) and different external genome annotations from Fig. 3A (columns) in hg19 (A) and hg38 (B) (**Methods**). Panel **(A)** is similar to Fig. 3A, but we present it here for better comparison with the hg38 enrichment heatmap. Results in **(B)** are based on (1) lifting over the full-stack annotation from hg19 to hg38 (**Methods**), and (2) doing enrichment analysis with annotated genome contexts derived from various databases in hg38 (**Methods**). In each heatmap, coloring of enrichments is column specific with highest and lowest enrichment values in each column are colored red and white, respectively. The first two columns of each heatmap show state labels and their percentage of genome coverage. The last row of each heatmap shows the percentage of genome coverage for each type of genome contexts. Below the heatmap in **(B)** is the correlation of the enrichments across states based on hg19 and hg38 for each corresponding annotation column as well as the average of them.

**Supplementary Figure 19:**
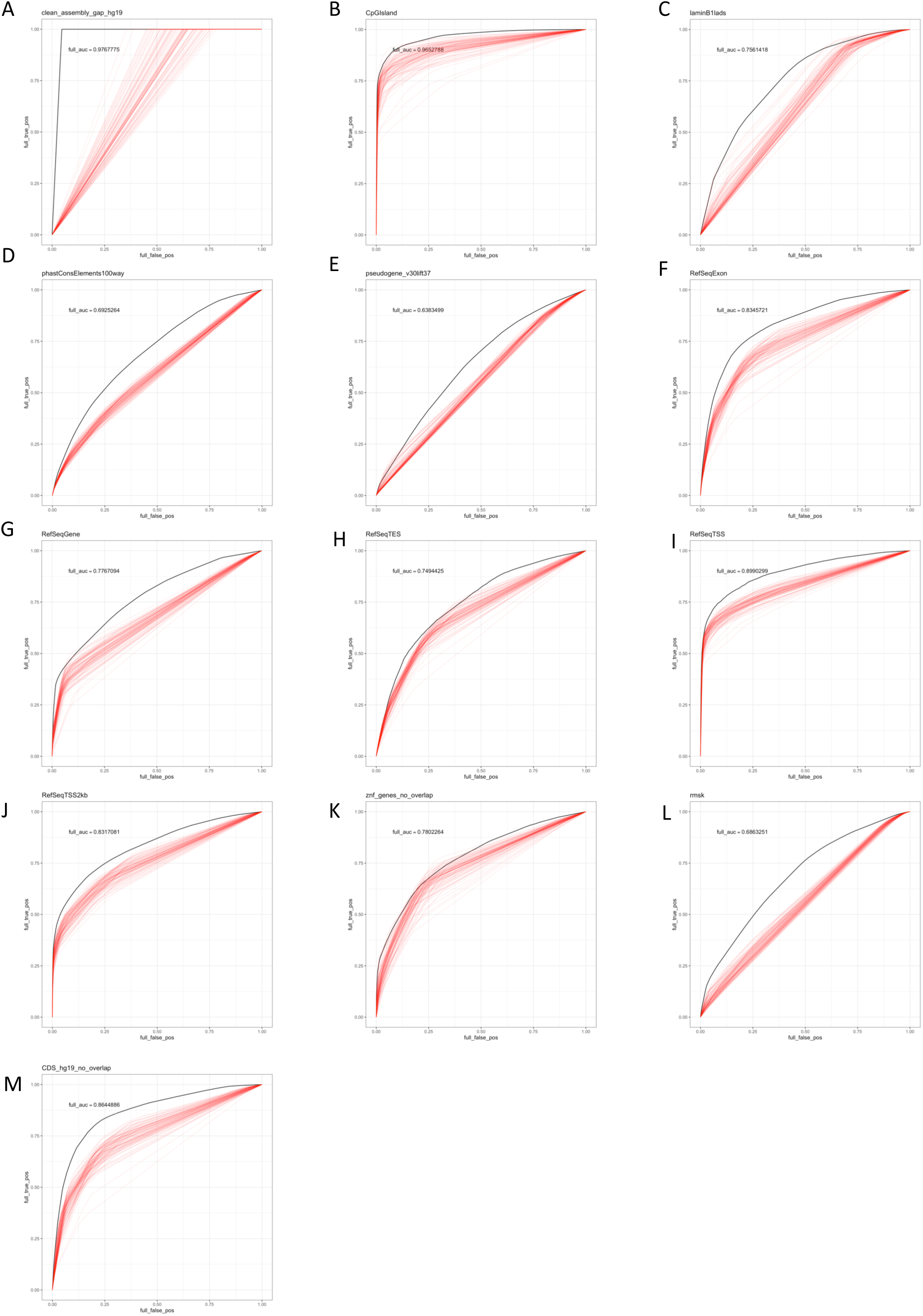
ROC comparison of full-stack model annotations and the 18-state per-cell-type concatenated model annotations for predicting various external genomic annotations. Each panel shows the ROC curves from using the full-stack model annotations and 98 per-cell-type chromatin state annotations from a concatenated model to predict different external genomic annotations (**Methods**). The per-cell-type annotations are from a previously learned 18-state concatenated model (Kundaje et al., 2015). The full-stack annotations’ ROC curves are in black, and 98 per-cell-type annotations’ ROCs are in red. The respective genomic contexts for panels A-M are assembly gaps, CpG Islands, lamina associated domains (laminB1lads), phastCons elements, pseudogenes, exons, gene bodies, transcription end sites (TES), transcription start sites (TSS), 2kb regions surrounding transcription start sites (TSS2kb), ZNF genes, repeat elements in UCSC Genome Browser’s repeatMasker track and coding sequences.

**Supplementary Figure 20:**
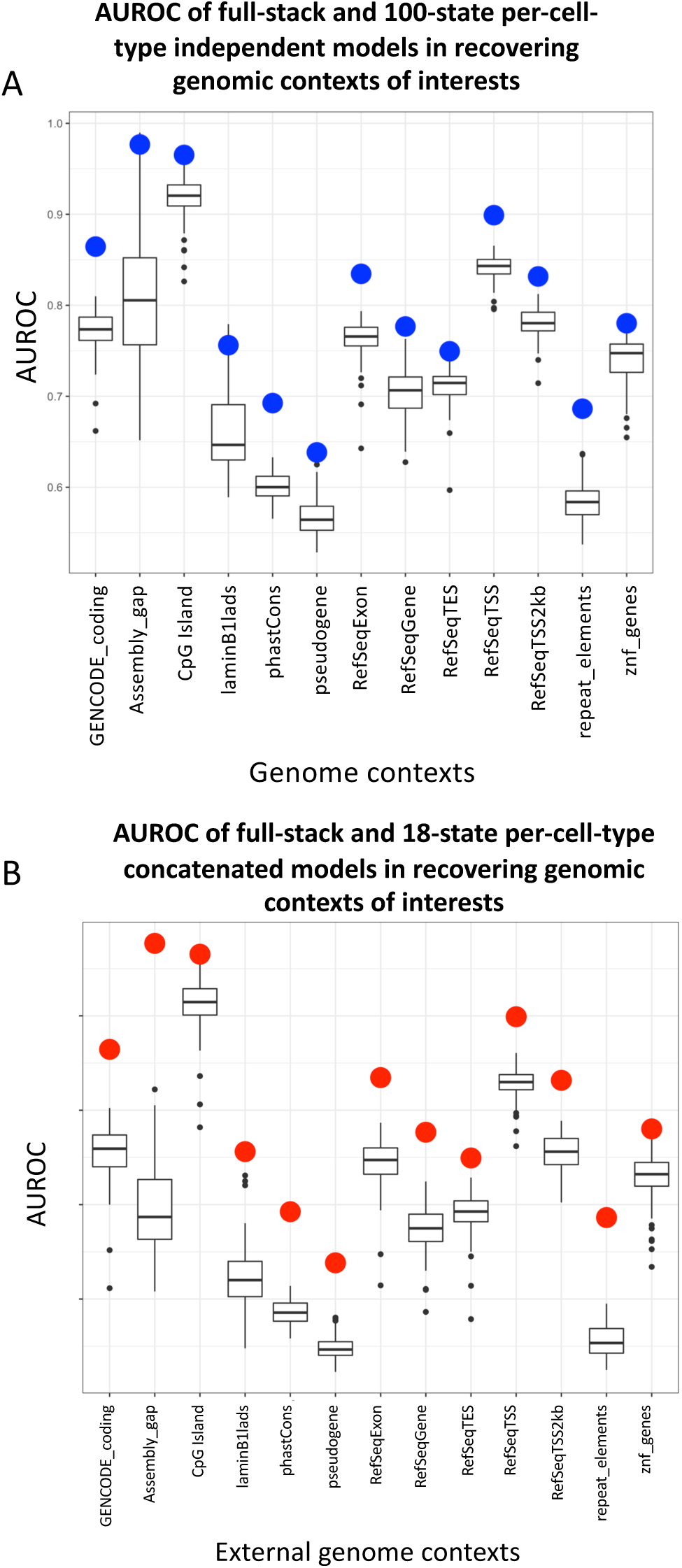
AUROC comparison of the full-stacked and the per-cell-type chromatin state annotations at predicting various external genomics annotations. **(A)** AUROC values for ROC curves in Supplementary Fig. 19. The x-axis represents different genomic contexts. The box-plots show AUROC of the 127 100-state per-cell-type annotations based on models learned independently in 127 cell types at predicting locations of the external annotations. The blue dots show the AUROC for the full-stack chromatin state annotations. **(B)** Similar to **(A),** but showing AUROC values for ROC curves in **Supplementary Fig. 21**. but the boxplots show the AUROC values for 98 18-state per-cell-type annotations based on concatenated models in 98 cell types. The red dots show the AUROC for the full-stack chromatin state annotations.

**Supplementary Figure 21:**
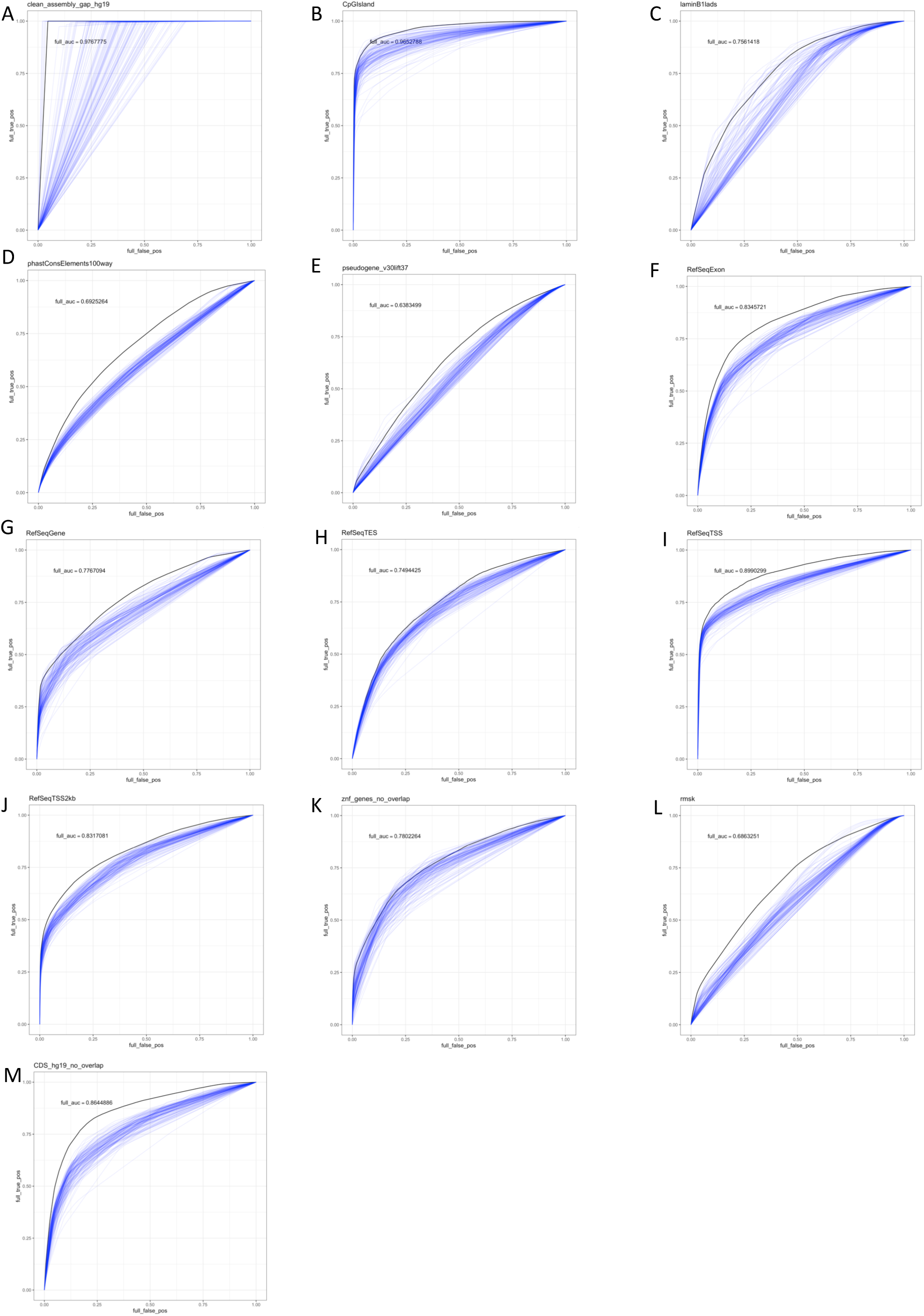
ROC comparison of full-stack model annotations and the 100-state per-cell-type-independent model annotations for predicting various external genomic annotations. Each panel shows the ROC curves from using the full-stack model annotations and the 127 per-cell-type-independent model chromatin state annotations to predict different external genomic annotations (**Methods**). The per-cell-type-independent models were 100 state models learned separately using all available data from each cell type. The full-stack annotations’ ROC curves are in black, and per-cell-type-independent annotations’ ROCs are in blue. The respective genomic contexts for panels A-M are assembly gaps, CpG Islands, lamina associated domains (laminB1lads), PhastCons elements, pseudogenes, exons, gene bodies, transcription end sites (TES), transcription start sites (TSS), 2kb regions surrounding transcription start sites (TSS2kb), ZNF genes, repeat elements from all classes and families in UCSC Genome Browser’s repeatMasker track and coding sequences.

**Supplementary Figure 22:**
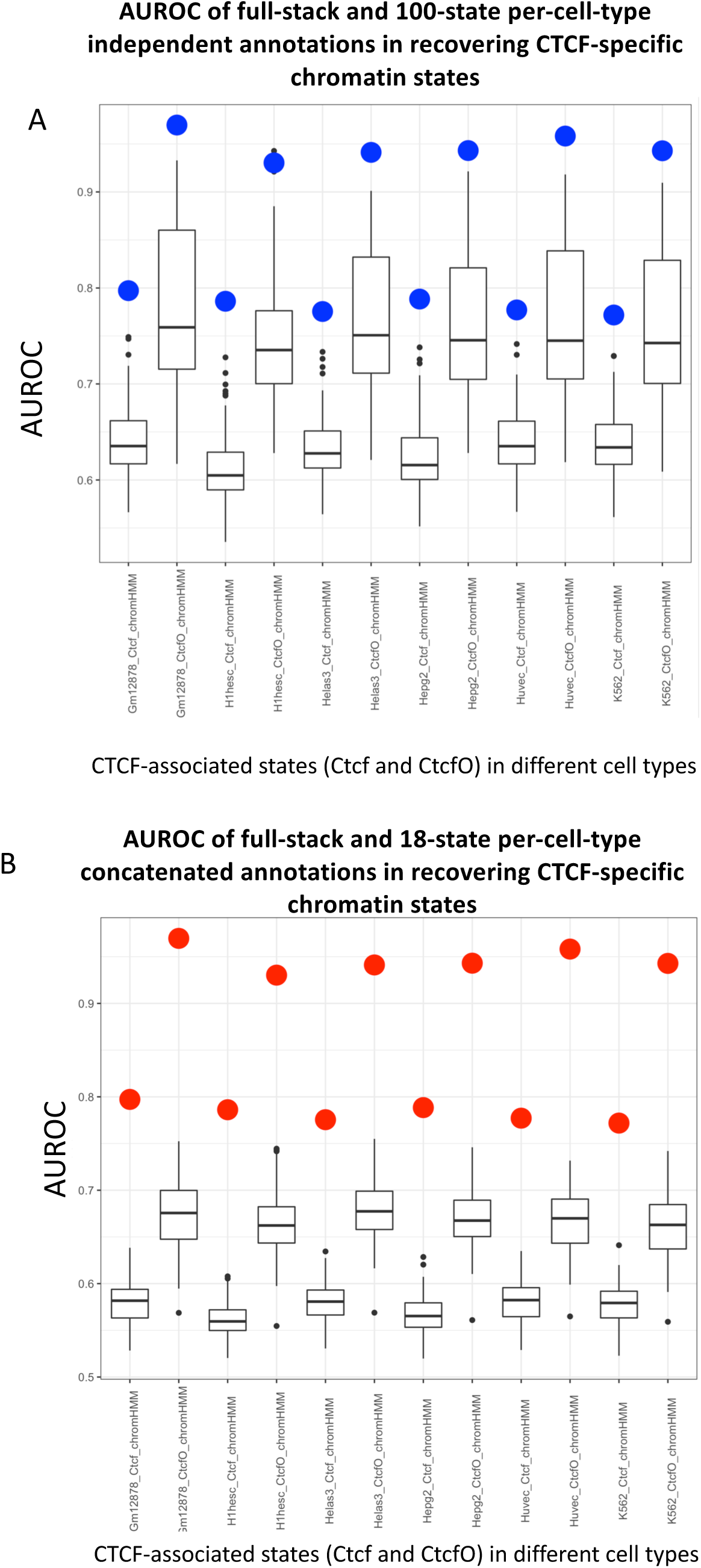
AUROC comparison of the full-stack and per-cell-type chromatin state annotations at predicting CTCF specific per-cell-type chromatin states. Box-plots show AUROC of **(A)** 127 100-state per-cell-type – independent and **(B)** 98 18-state per-cell-type-concatenated model annotations, which did not include CTCF, at predicting bases in sets of per-cell-type CTCF-associated chromatin states. In both panels, the x-axis represents sets of chromatin states associated with CTCF signal and limited histone modification signal in one of six cell types from a previously published chromatin state model that included CTCF (Hoffman et al., 2013) (**Methods**). CtcfO corresponds to a state that also had open chromatin signals, while state Ctcf lacked those signals. The dots colored **(A)** blue and **(B)** red show the AUROC for the full-stack chromatin state annotations, which were not trained using CTCF signals data.

**Supplementary Figure 23:**
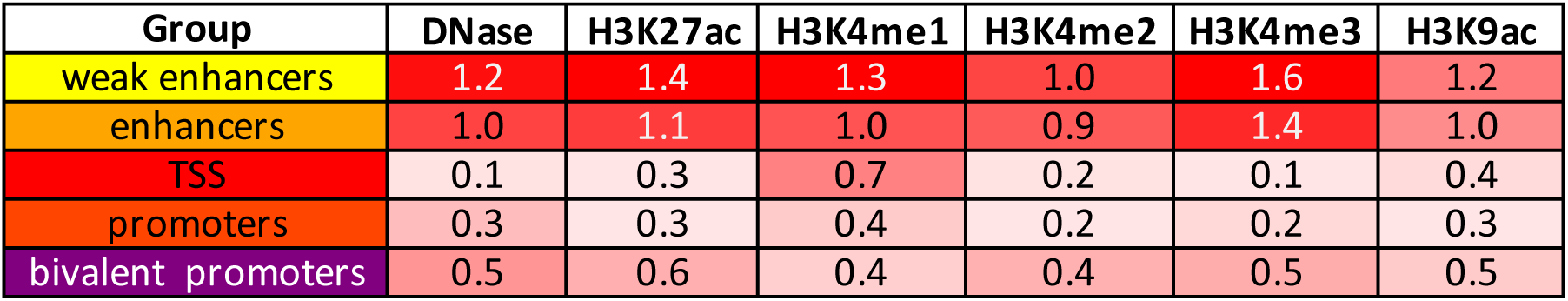
Coefficient of variations of emission probabilities across different cell groups. Average coefficient of variations for the five enhancer and promoter state groups of full-stack states (rows) and six chromatin marks that are associated with enhancer and promoter activities. For a mark and state group combination, the coefficient of variation for the mark emission was computed separately for each state and then averaged among states in the group. The enhancer and weak enhancer group showed greater than two-fold higher coefficient of variations compared to the promoter group.

**Supplementary Figure 24:**
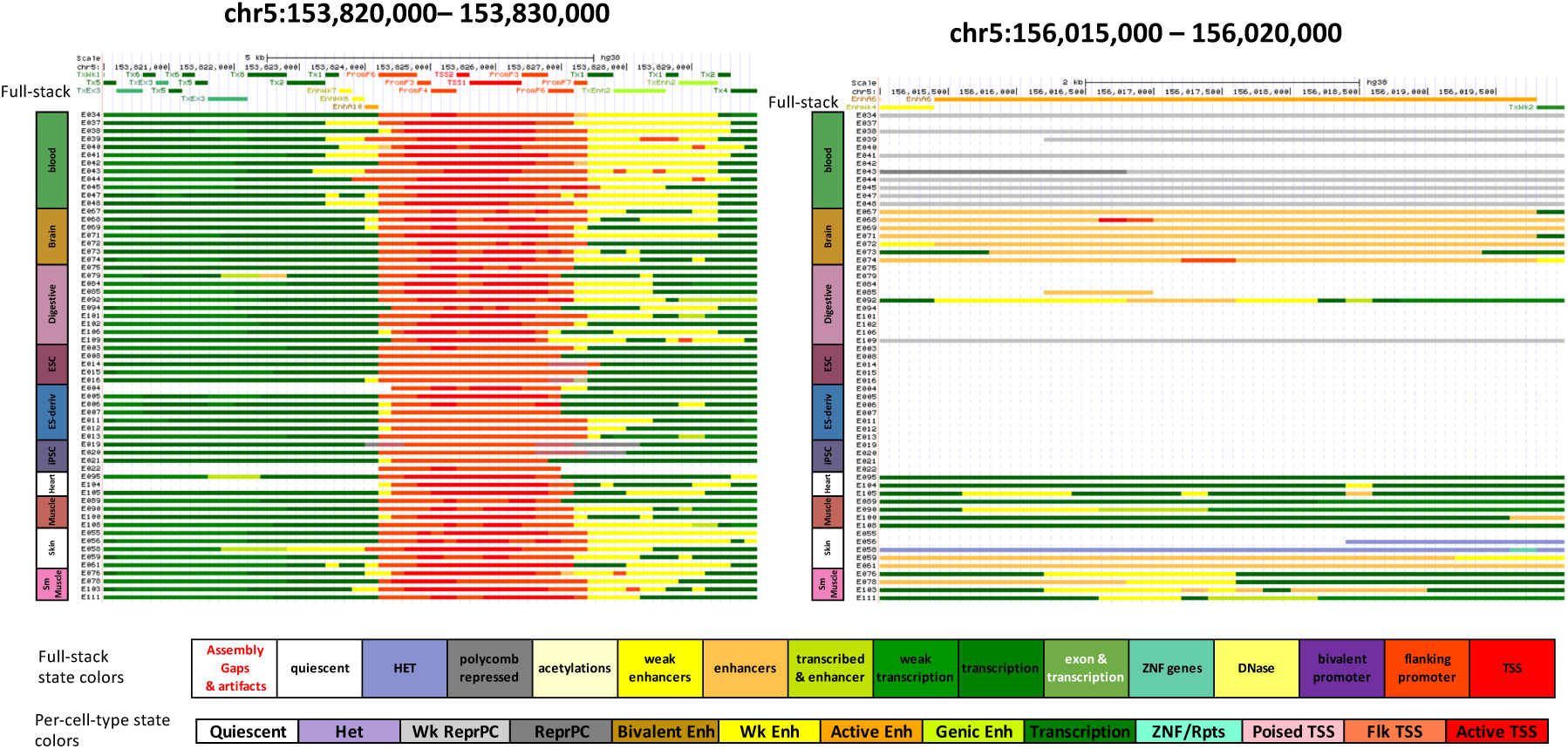
Illustration of the full-stack annotations. Two loci representing regions that are in transcribed and active promoter states across cell types (left), and in an enhancer state specifically in brain (right). The loci correspond to those presented in Fig. 1. The top track shows the full-stack state annotations. The following tracks show per-cell-type-concatenated annotations from 18-state models based on imputed data (Kundaje et al., 2015). The cell types are ordered based on their associated cell groups. A color legend for the states is shown along the bottom.

**Supplementary Figure 25:**
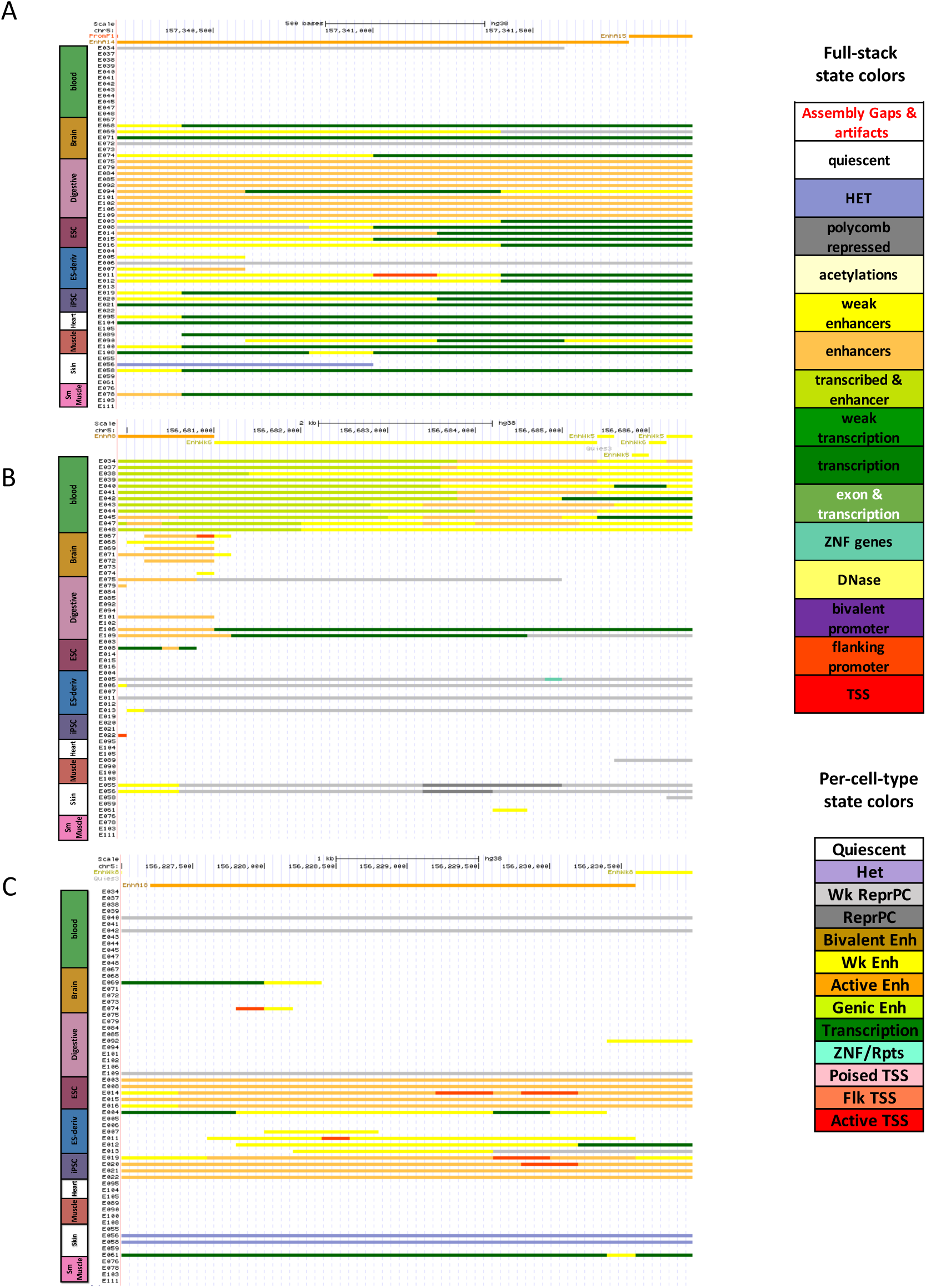
Illustration of full-stack cell-type-specific enhancer states. (A-C) The first track in each panel demonstrates the full-stack state annotation. Each of the following tracks show chromatin state annotations from a 18-state per-cell-type-concatenated model (Kundaje et al., 2015). The individual reference epigenomes IDs and their tissue groups are labeled on left. The chromatin state coloring is labeled on right. **(A)** A genomic region (chr5:157340200-157342000) annotated to an active enhancer state in digestive cells in the full-stack model (EnhA14). **(B)** A genomic region (chr5:156679900-156686500) annotated to blood enhancer states in the full-stack model (EnhWk6 and EnhA8). **(C)** A genomic region (chr5:156227000-156231000) annotated as an ESC/iPSC-specific enhancer state in the full-stack model (state EnhA18).

**Supplementary Figure 26:**
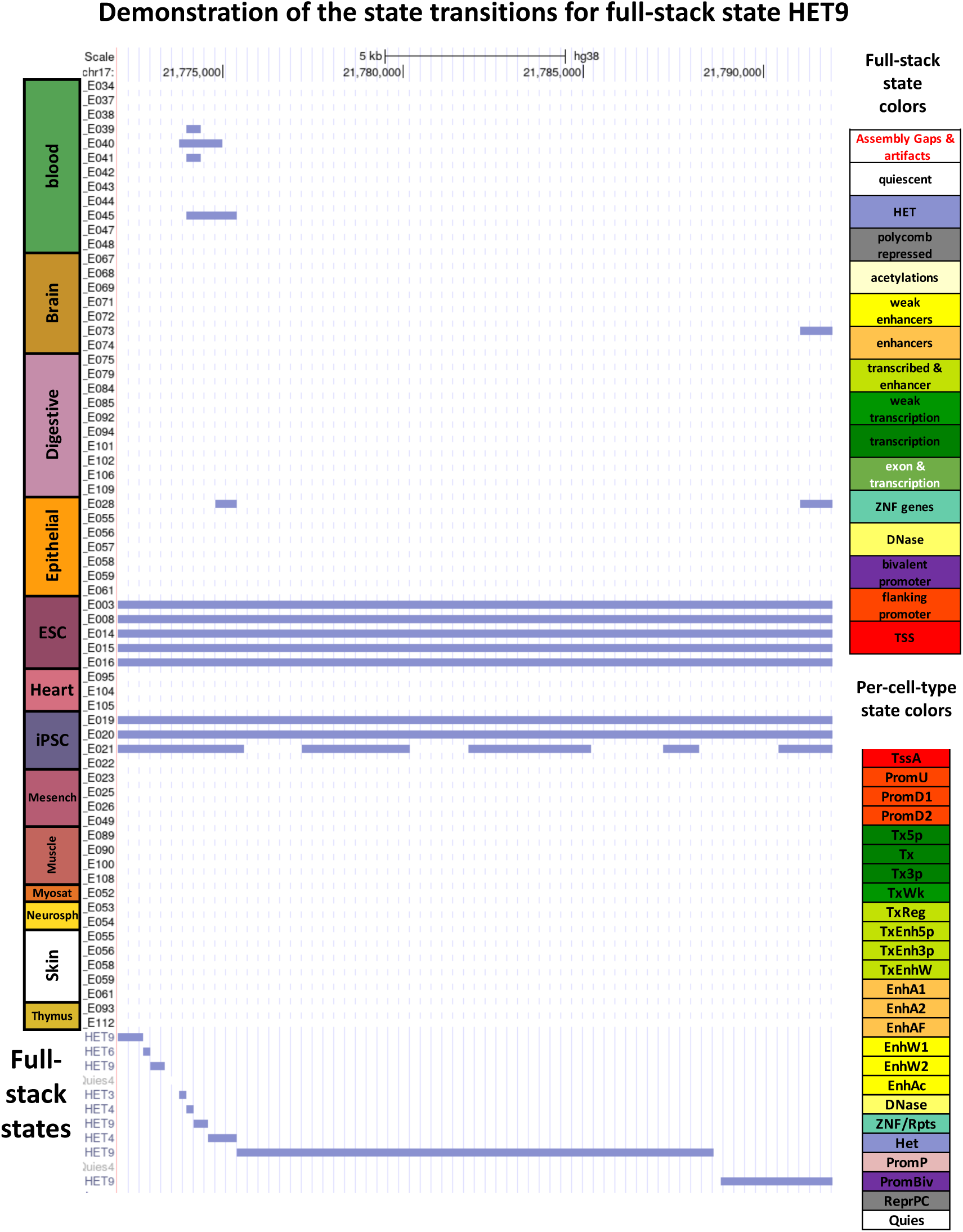
Illustration of full-stack heterochromatin state HET9. The figure captures the per-cell-type-concatenated chromatin state maps for various reference epigenomes, and the corresponding full-stack chromatin state maps at region chr17:21772099-21791900. The first 66 tracks show chromatin state annotations from a 25-state concatenated model for 66 reference epigenomes (equivalently, in this paper, cell types) (Ernst & Kellis, 2015). The individual reference epigenomes IDs and their tissue groups are labeled on the left. The chromatin state colors are explained on the right. The last track, shown in full mode to display all state labels on the right, corresponds to the full-stack chromatin state map at this region. State HET9 is characterized, based on our analysis, as an ESC-group-related heterochromatin state (Fig. 2C**, Supp. Fig. 8-9, Supplementary Data 1,3**). Detailed characterizations of all full-stack states are in Supplementary Data.

**Supplementary Figure 27:**
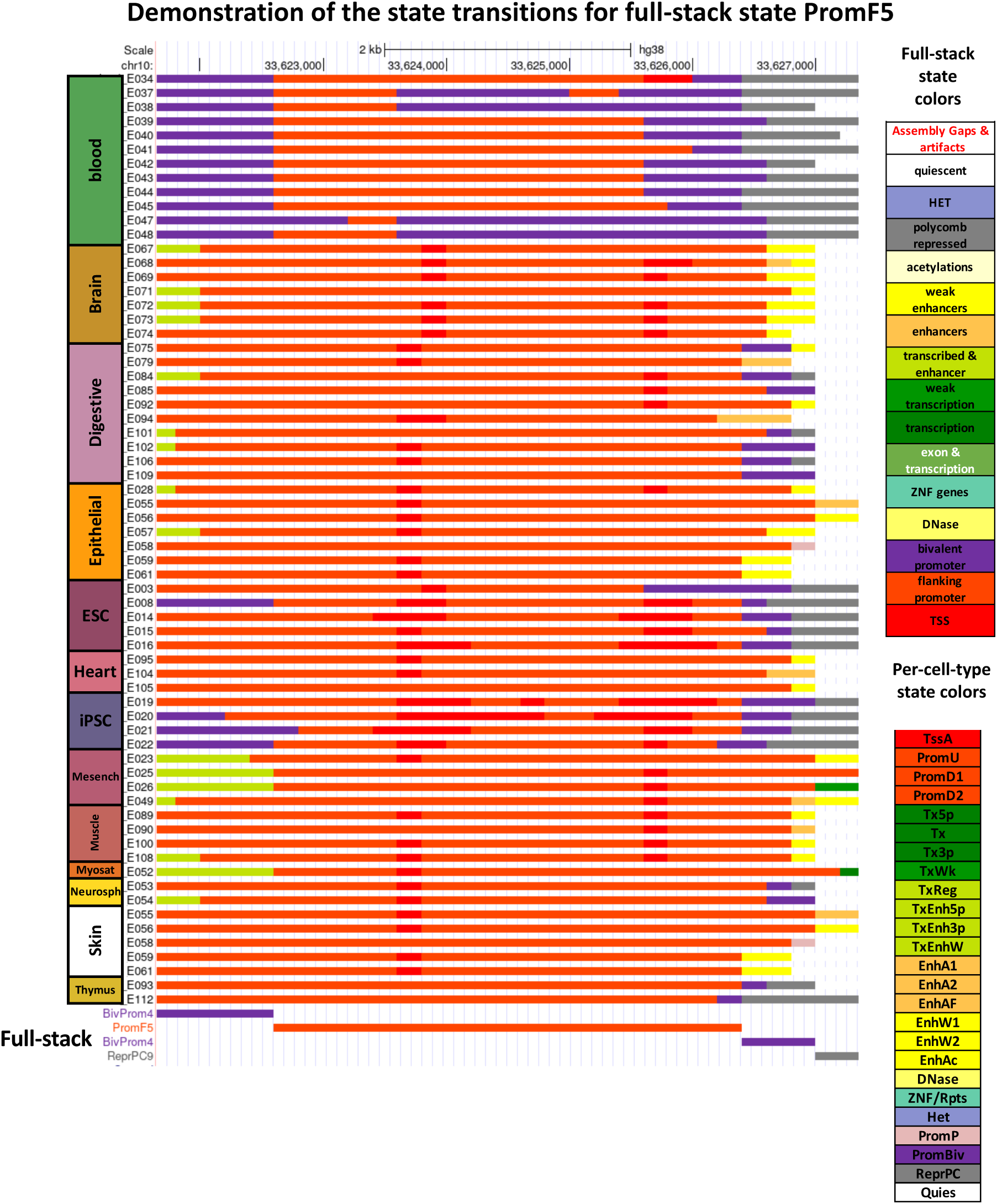
Illustration of full-stack flanking promoter state PromF5. The figure captures the per-cell-type-concatenated chromatin state maps for various reference epigenomes, and the corresponding full-stack chromatin state maps at region chr10:33621649-33627350. The first 66 tracks show chromatin state annotations from a 25-state concatenated model for 66 reference epigenomes (equivalently, in this paper, cell types) (Ernst & Kellis, 2015). The individual reference epigenomes IDs and their tissue groups are labeled on the left. The chromatin state coloring is labeled on the right. The last track, shown in full mode to display all state labels on the left, corresponds to the full-stack chromatin state map at this region. State PromF5 is characterized, based on our various analyses as state frequently found at flanking promoter regions with some upstream bias, and sometimes, this state overlaps with regions of bivalent promoters in Blood-related and ESC- related groups (Blood & T cells, HSC & B cells, ESC, iPSC and ES-deriv) (**Supp. Fig. 8-9, Supplementary Data 1,3**). Detailed characterization of all full-stack states are in **Supplementary Data 1-3**.

**Supplementary Figure 28:**
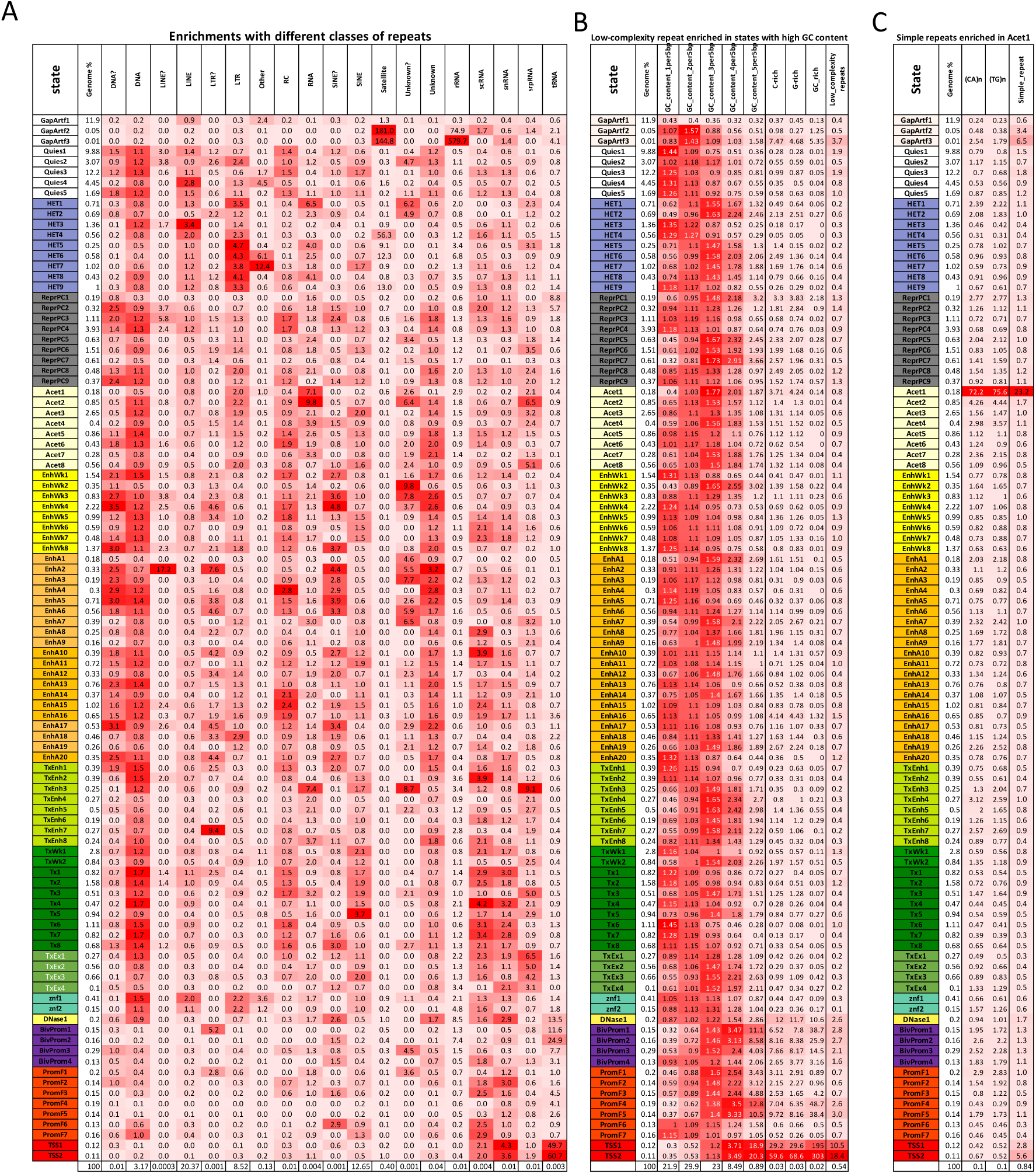
Full stack states enrichments with RepeatMasker classes of repeats (A), low-complexity repeats and GC content (B), simple repeats (C). This is an extended version of Fig. 4C. In each panel, the rows correspond to full-stack states. The second column reports the percentage of the genome that each full-stack state occupies. In **(A)**, columns 3-21 correspond to different repeat classes. In **(B)**, columns 3-7 correspond to 5bp windows in the genome are stratified by the number of G/C bases in them, columns 8-10 correspond to regions enriched with C-rich, G-rich, and GC-rich low complexity sequences, respectively, and column 11 shows enrichments for all low complexity sequences from RepeatMasker. States TSS1-2 are most enriched with Low complexity repeat class, which is consistent with these states having a high enrichment (19-20 fold) for windows in which all bases are a G or C. In **(C)**, columns 3-4 correspond to simple repeats of repeated (CA) and (TG) sequences, and column 5 shows enrichments for all simple repeats. State Acet1 is most enriched with simple repeats and this enrichment is mostly driven by enrichments with repeated CA and TG dinucleotides. In each panel, the values in all columns except the first and second columns correspond to fold enrichment for different repeat contexts in the full-stack states. Values are colored on a column-specific color scale. The last row gives the percentage of the genome that each repeat class occupies.

**Supplementary Figure 29:**
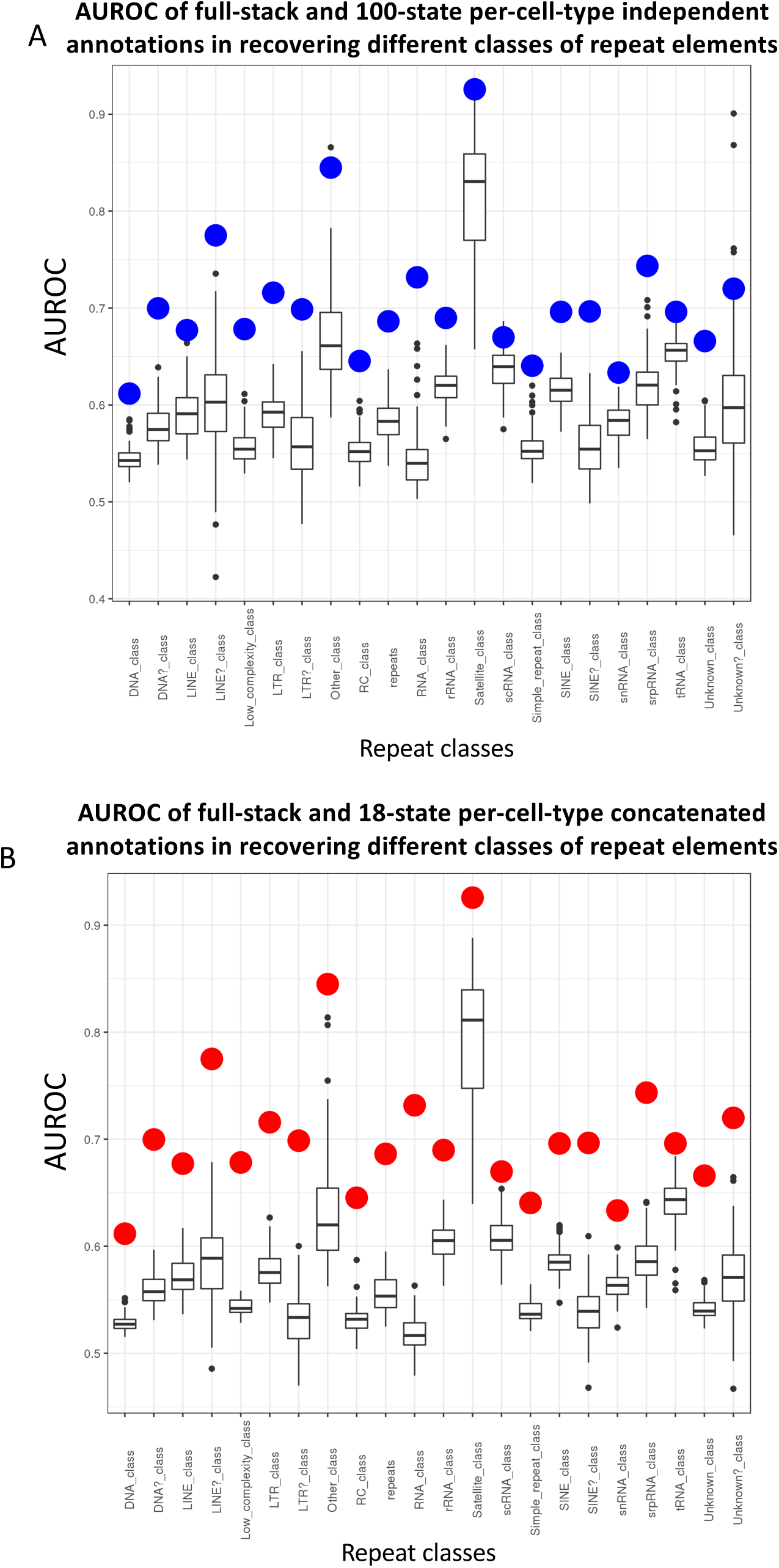
AUROC comparison of the full-stack and per-cell-type chromatin state annotations at predicting different classes of repeat elements. Box-plots showing AUROC of the **(A)** 127 100-state per-cell-type annotations from independent models and **(B)** 98 18-state per-cell-type annotations from a concatenated model at predicting bases in different repeat classes labeled on the x-axis. The dots colored **(A)** blue and **(B)** red show the AUROC for the full-stack chromatin state annotations.

**Supplementary Figure 30:**
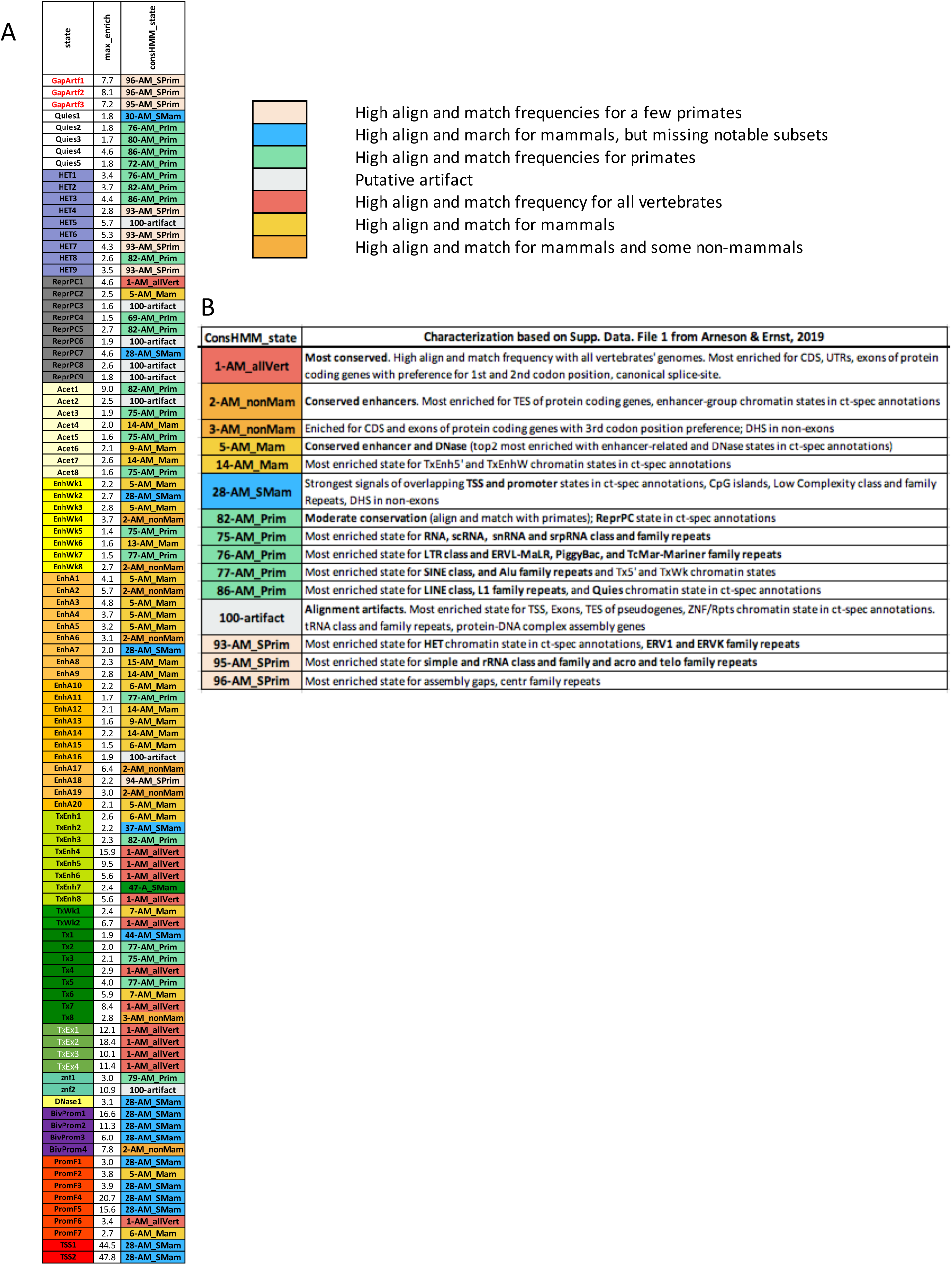
Full-stack states and maximum enriched ConsHMM state. The first column gives the label of the full-stack states. The second column shows the maximum fold enrichment for any ConsHMM state defined to annotate nucleotides based on sequence conservation patterns (Arneson & Ernst, 2019) (**Methods**). The third column shows the ConsHMM state that has the highest fold-enrichment in each full-stack state. One notable ConsHMM state is state 1 (1-AM_allVert), representing regions with high probabilities of aligning and matching the human reference genome for all vertebrates and the most enriched for exons. Full-stack states in the transcription-exon group (TxEx) are all maximally enriched with ConsHMM state 1. Another notable ConsHMM state, state 28 (28-AM_SMam), was the ConsHMM most strongly enriched for overlapping annotated TSS. Consistent with this, this state is also the maximum-enriched ConsHMM state in many full-stack states in TSS and Promoter flanking groups.

**Supplementary Figure 31:**
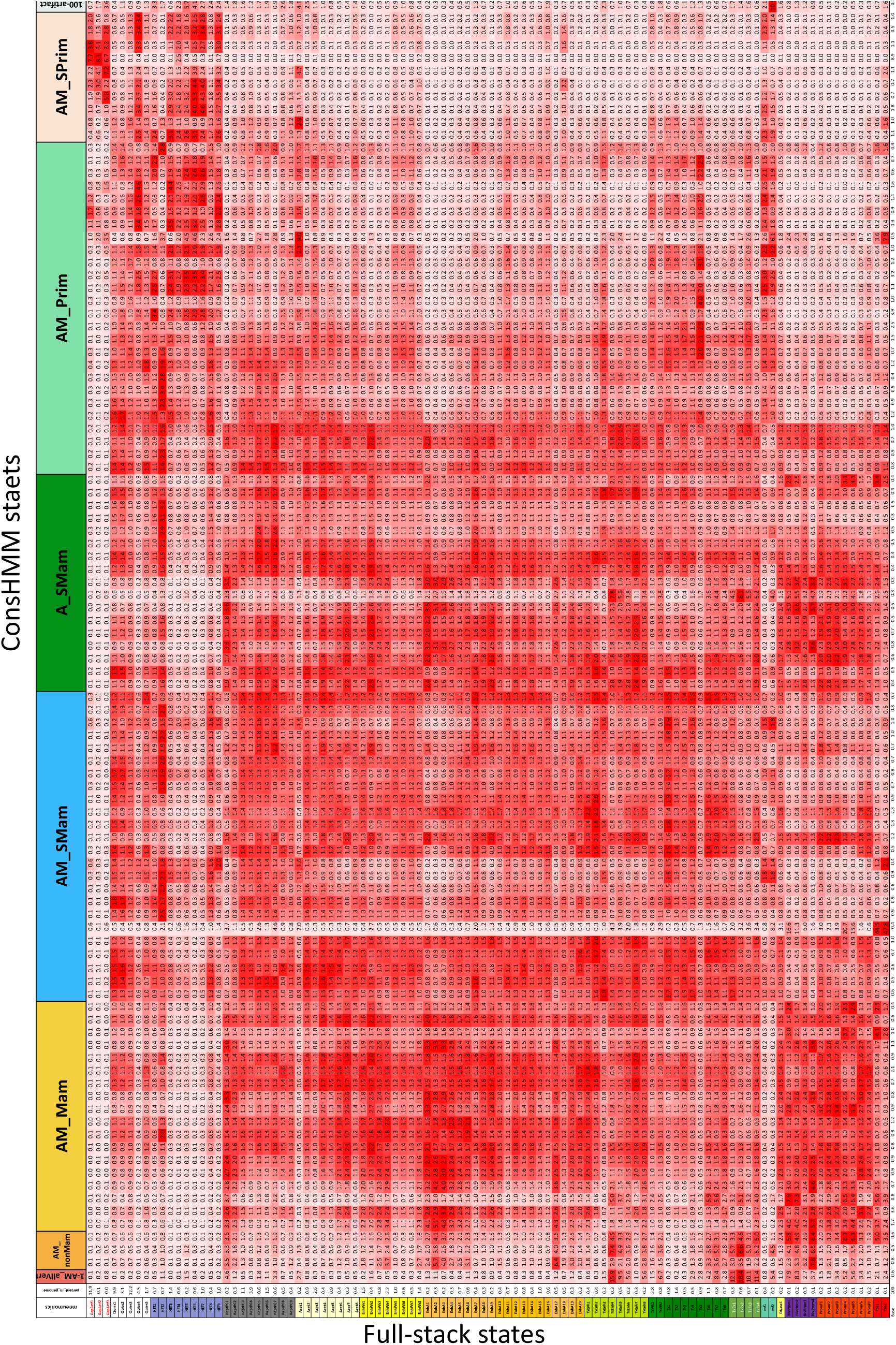
Enrichment of all full-stack states for ConsHMM states. The figure shows the enrichment of each full-stack state for each ConsHMM state from a 100-state model based on a 100-way vertebrate alignment (Arneson & Ernst, 2019). Rows correspond to different full-stack states. The header row gives the ConsHMM state labels, where ConsHMM states are placed in groups previously defined based on their patterns of sequence alignment with other vertebrates (Arneson & Ernst, 2019), colored as in **Supplementary Fig. 30**. The second column shows the percentage of the genome that each full-stack state falls into. Each of the remaining columns corresponds to one ConsHMM state. Values in the columns are colored on a column specific coloring scale. The last line in the heatmap gives the percentage of the genome that is covered by each ConsHMM state.

**Supplementary Figure 32:**
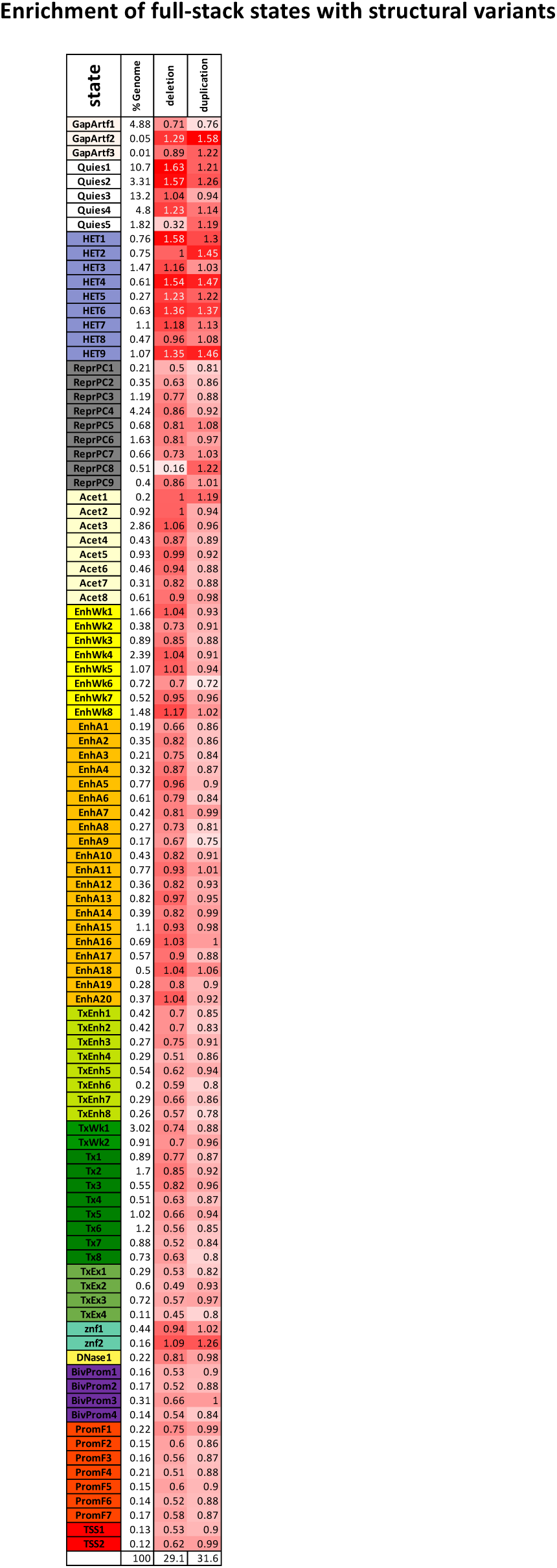
Full-stack states enrichments with structural variants. The rows correspond to full-stack states. The first column presents the state labels, the second presents the percentage of the genome in hg38 that each state occupies, and the last two columns correspond to two different types of structural variants: deletions and duplications. The values correspond to the full-stack states’ fold enrichment for the structural variant type. Values are colored on a column-specific color scale. The last row gives the percentage of the genome that each type of structural variants occupies.

**Supplementary Figure 33:**
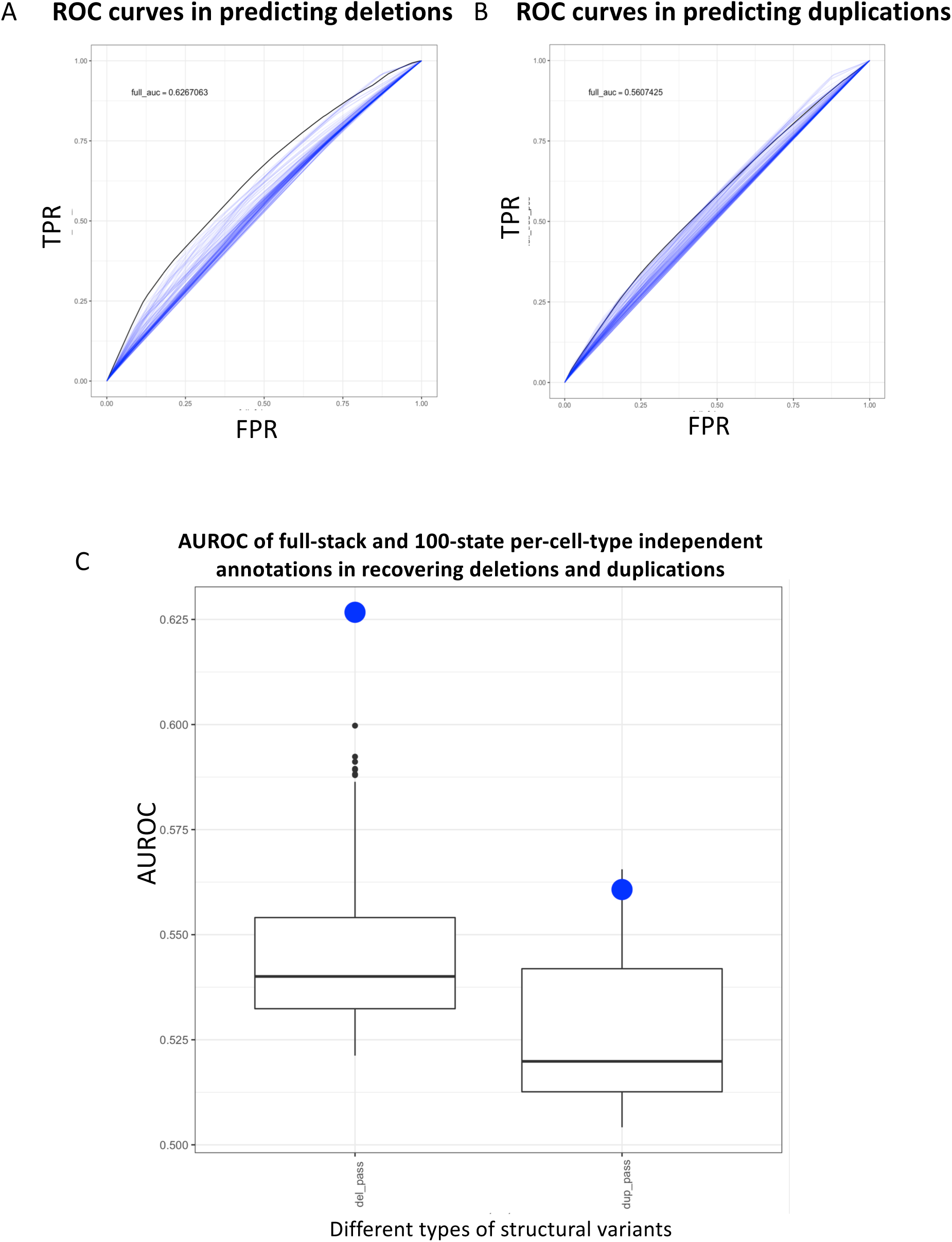
Comparison of full-stack model annotations and the 100-state per-cell-type-independent model annotations in predicting structural variants of type deletions and duplications. **(A)** ROC curves for the full-stack model and the 127 100-state per-cell-type-independent models’ chromatin state annotations at predicting bases covered by deletions (**Methods**). The full-stack model’s annotation ROC curve is in black and the 127 100-state per-cell-type-independent models’ annotation ROCs are shown in blue. **(B)** Similar plot as (A), but for duplications. **(C)** Comparison of the AUROC in predicting structural variants. The x-axis represents different types of structural variants. The box-plots show AUROC for 127 100-state per-cell-type-independent models’ in predicting deletions and duplications. The blue dots show the AUROC of the full-stack chromatin state annotation.

**Supplementary Figure 34:**
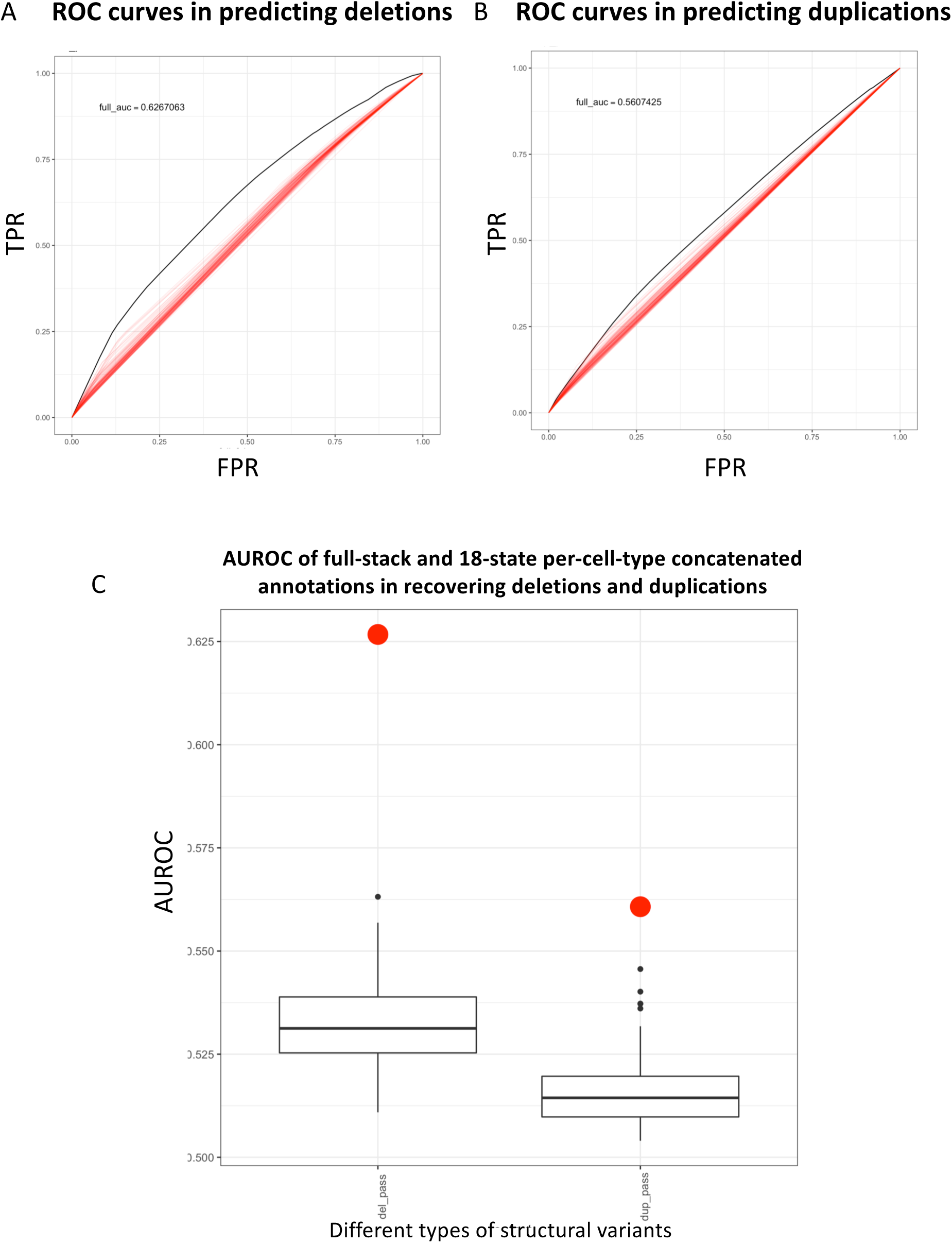
Compare full-stack model annotations and 18-state per-cell-type-concatenated model annotations in predicting structural variants of type deletions and duplications. **(A)** ROC curves for the full-stack model and 98 per-cell-type-concatenated models’ chromatin state annotations at predicting bases covered by deletion (**Methods)**. The full-stack model’s annotation ROC curve is in black and the 98 18-state per-cell-type annotations from concatenated models ROCs are shown in red. (**B)** Similar plot as (A), but for duplications. **(C)** Comparison of the AUROC in predicting structural variants. The x-axis represents different types of structural variants. The box-plots show AUROC of 98 18-state per-cell-type-concatenated models’ in predicting deletions and duplications. The red dots show the AUROC of the full-stack chromatin state annotations in predicting bases in each type of structural variant.

**Supplementary Figure 35:**
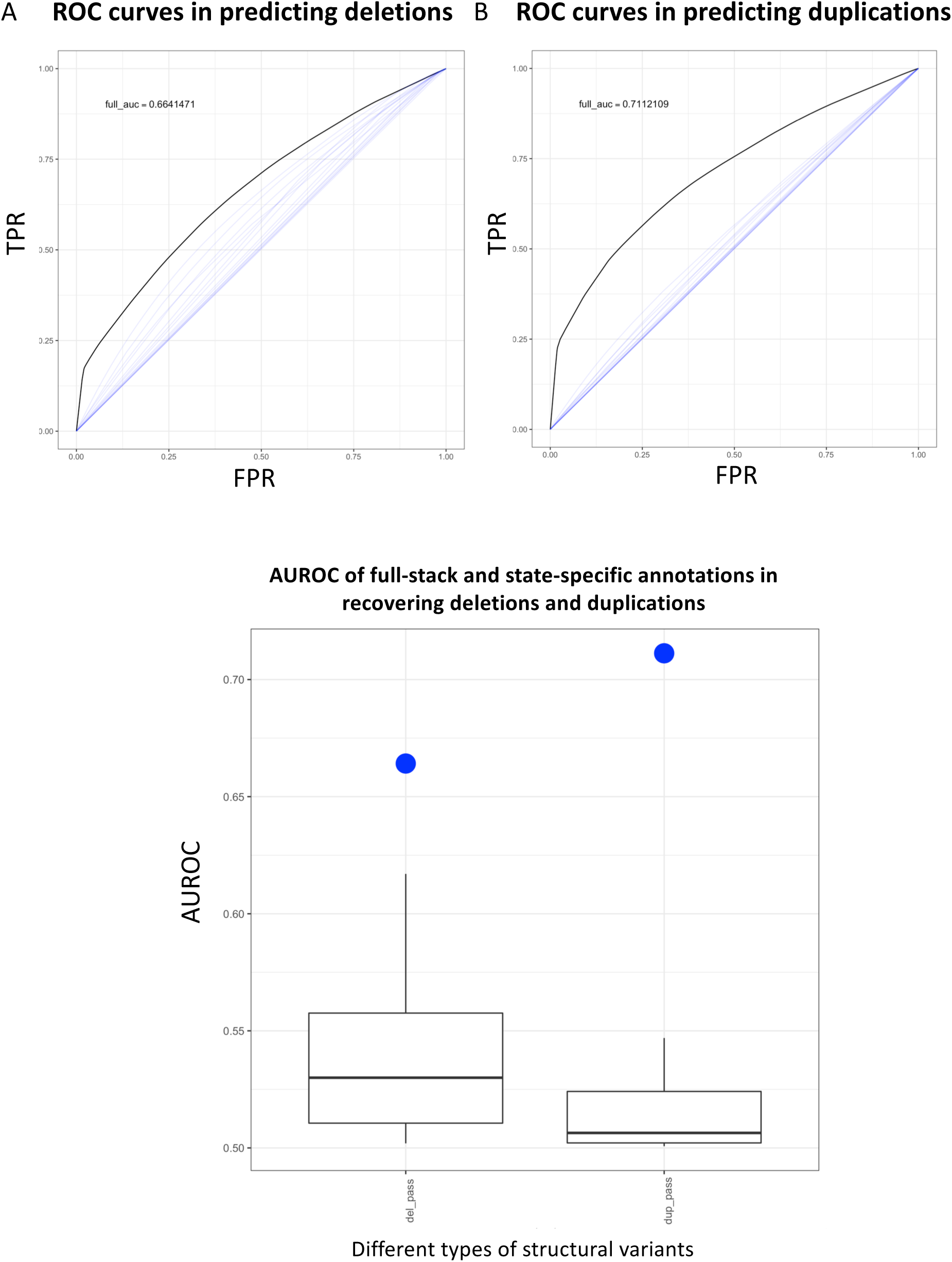
Compare Full-stack states vs. state-specific annotations in predicting structural variants of types deletions and duplications. We followed the procedure outlined in (Abel et al., 2020) to compute the enrichments between annotations associated with one chromatin state and structural variants. In particular, we utilized 15-state chromatin state annotation for 127 reference epigenomes from Roadmap Epigenomics Consortium. Then, for each of the 15 states, we stratified genomic positions based on the number of cell types in which the state is present (ranging from 0 to 127), resulting in 15 state-specific models’ annotations (**Methods**). **(A)** ROC curves for the full-stack model and 15 state-specific models’ annotations at predicting bases covered by deletions (**Methods**) The full-stack model’s ROC curve is in black, and state-specific models’ ROCs are shown in blue. (**B)** Similar plot as (A), but for duplications. **(C)** Comparison of the AUROC in predicting structural variants. The x-axis represents different types of structural variants. The box-plots show AUROC of 15 state-specific models’ annotation in predicting deletions and duplications. The blue dots show the AUROC of the full-stack chromatin state annotations in predicting respective types of structural variants.

**Supplementary Figure 36:**
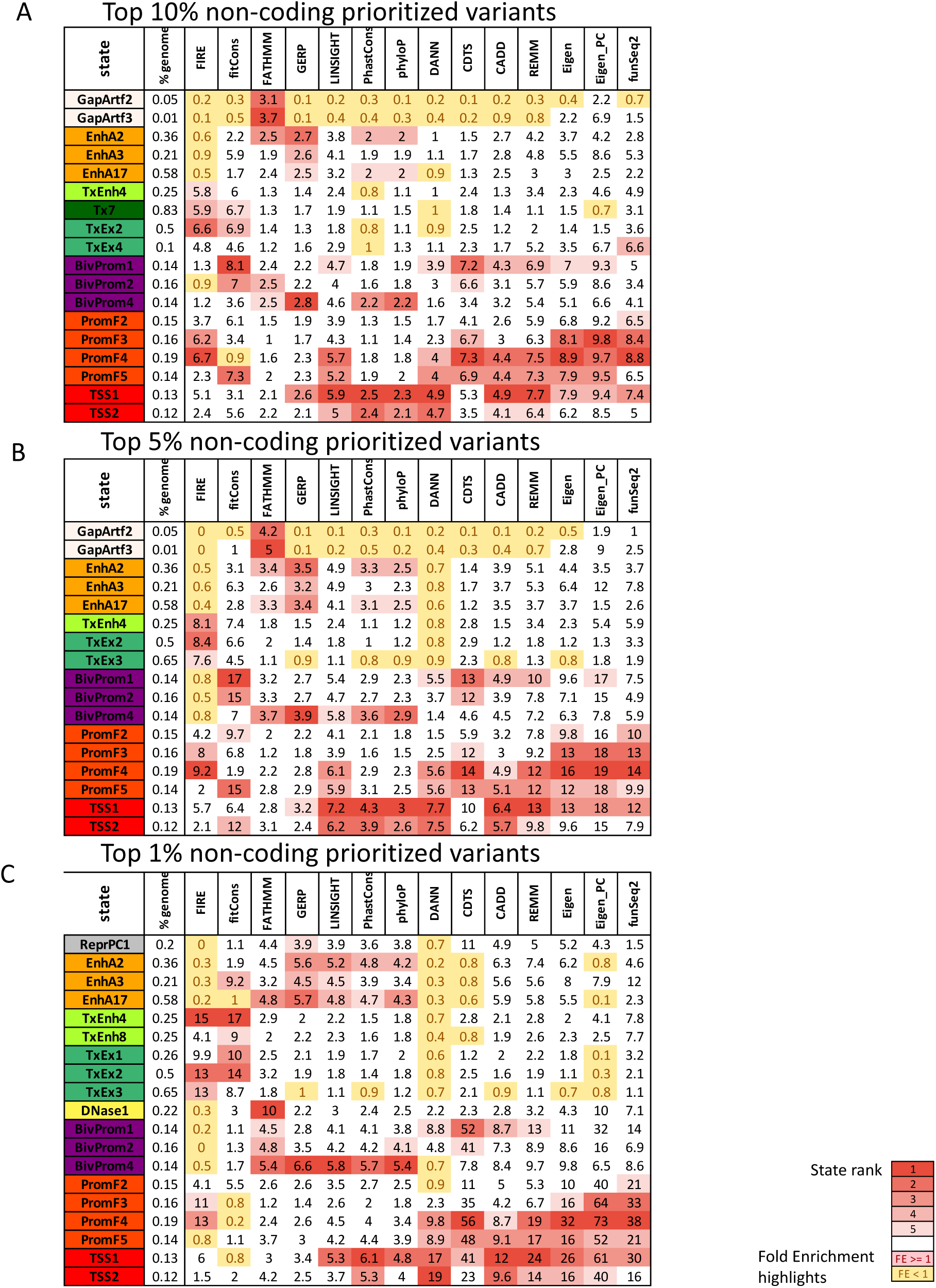
Enrichment of selected full-stack states with prioritized variants. Extended version of figure Fig. 5C showing fold enrichment of full-stack states for genomic bases prioritized in the **(A)** top 10% **(B)** top 5%, and **(C)** top 1% among non-coding bases by 14-different variant prioritization scores previously curated in (Arneson & Ernst, 2019) (Methods). Only states that were among the top five with greatest enrichments for at least one score are shown. Top enrichment values are colored red based on the rank of the state for each score as indicated in the color legend at the bottom. Depletions are shown in yellow.

**Supplementary Figure 37:**
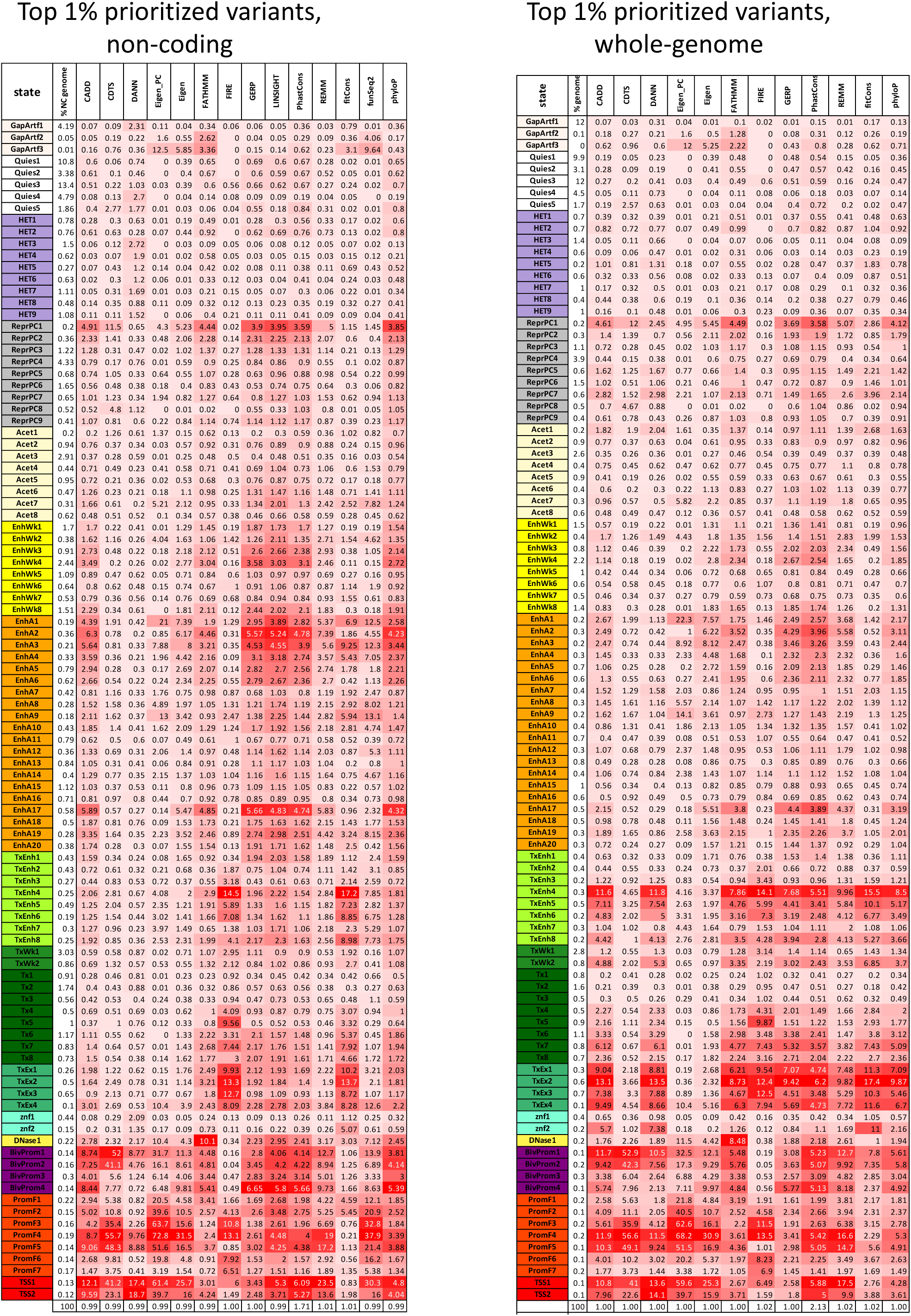
Enrichment of all full-stack states for top 1% bases prioritized by variant prioritization scores. Extended version of figure Fig. 5C showing the enrichment values of all full-stack states for genomic bases prioritized in the top 1% prioritized bases **(A)** in non-coding genome, and **(B)** genome-wide, by various variant prioritization scores. Coloring of enrichments is column specific. The second column in each heatmap, to the right of the state labels, shows the percentage of the background region (non-coding genome in **(A)** and whole genome in **(B)**) that each full-stack state covers. The last line in both heatmaps gives the actual percentage of the background region that is covered by each set of prioritized variants, which can differ from 1% exactly because of how ties of prioritization scores among bases were handled.

**Supplementary Figure 38:**
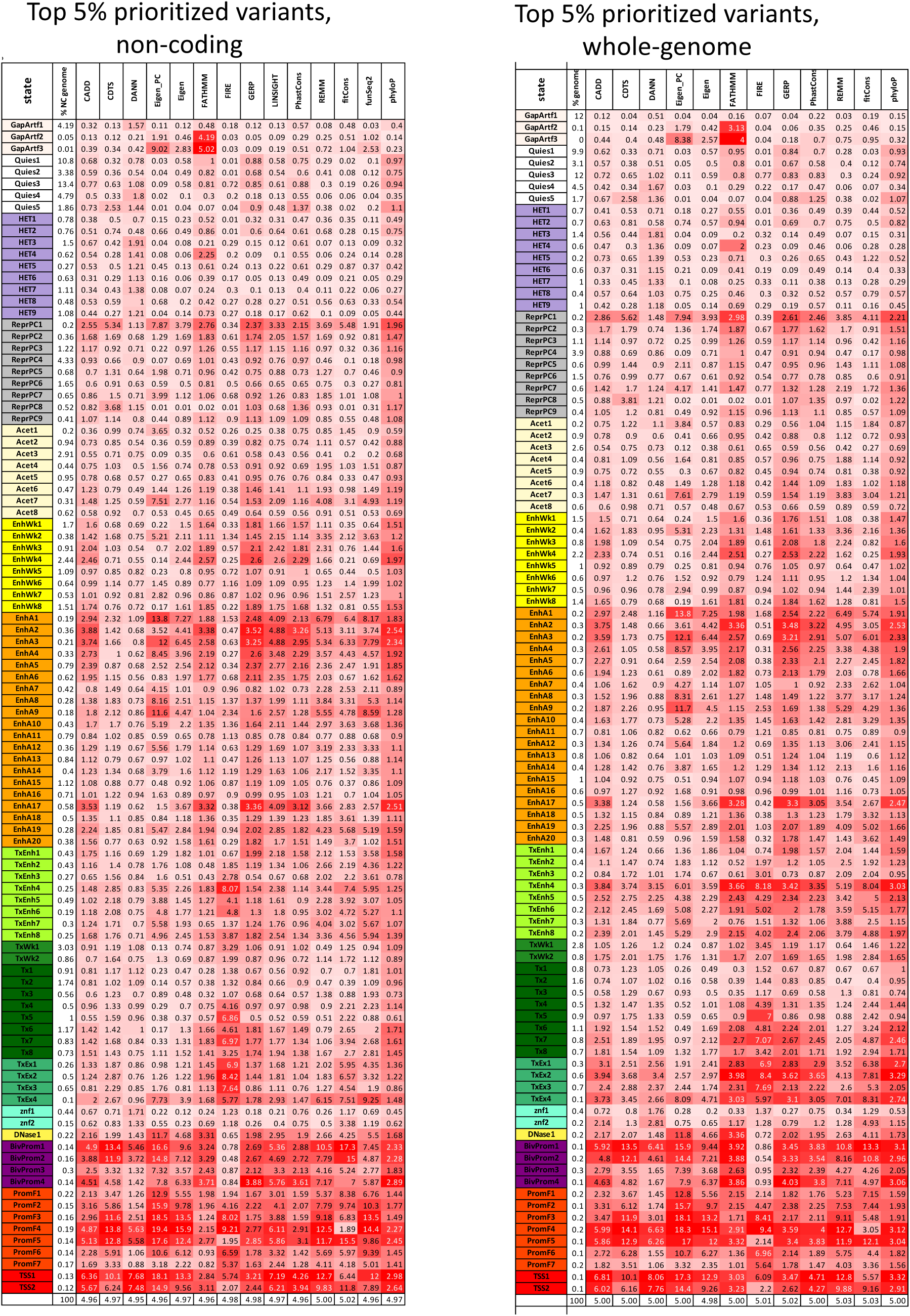
Enrichment of all full-stack states for top 5% bases prioritized by variant prioritization scores. Analogous to **Supplementary Fig. 37**, but for top 5% prioritized bases.

**Supplementary Figure 39:**
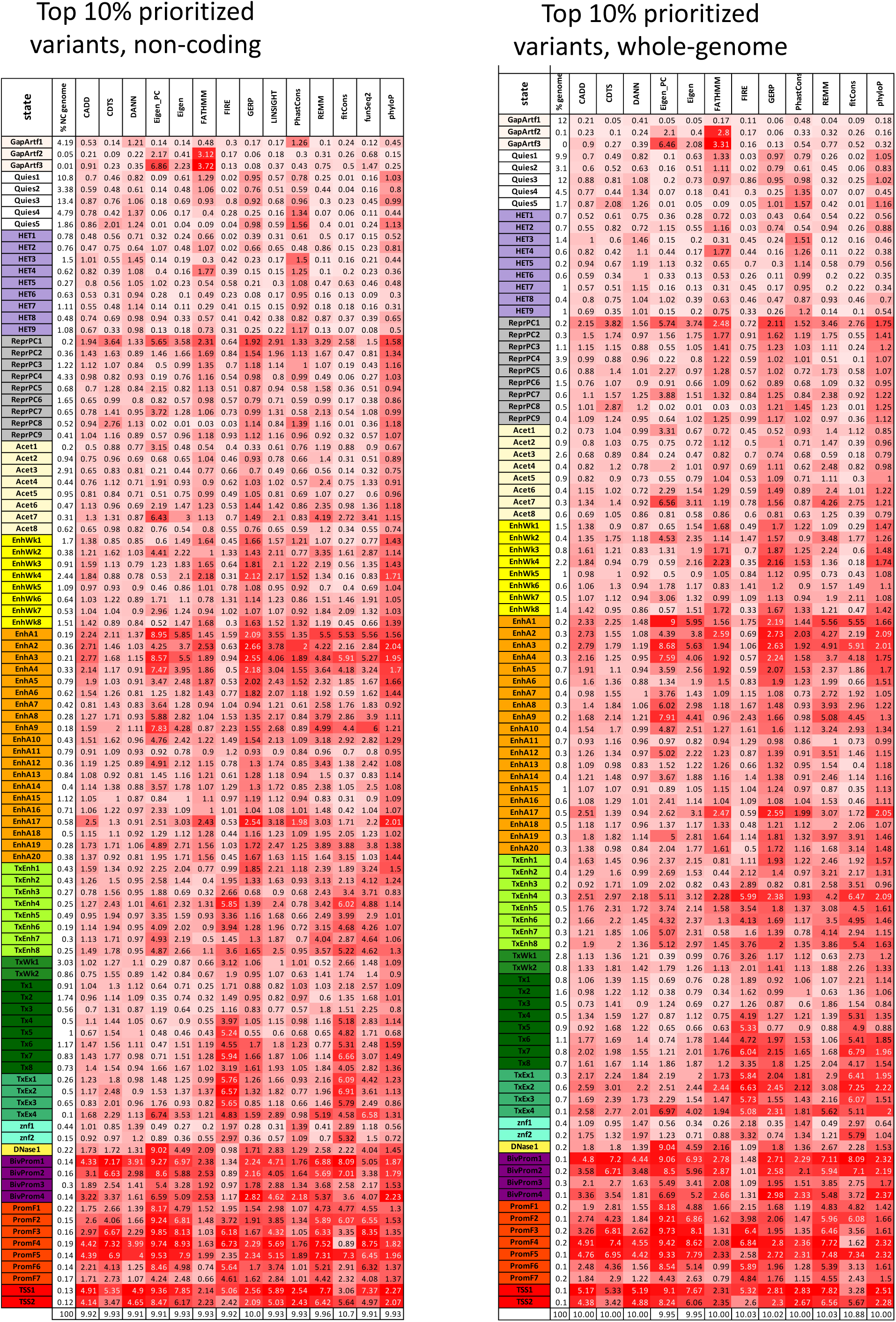
Enrichment of all full-stack states for top 10% bases prioritized by variant prioritization scores. Analogous to **Supplementary Fig. 37** and **Supplementary Fig. 38**, but for top 10% prioritized bases.

**Supplementary Figure 40:**
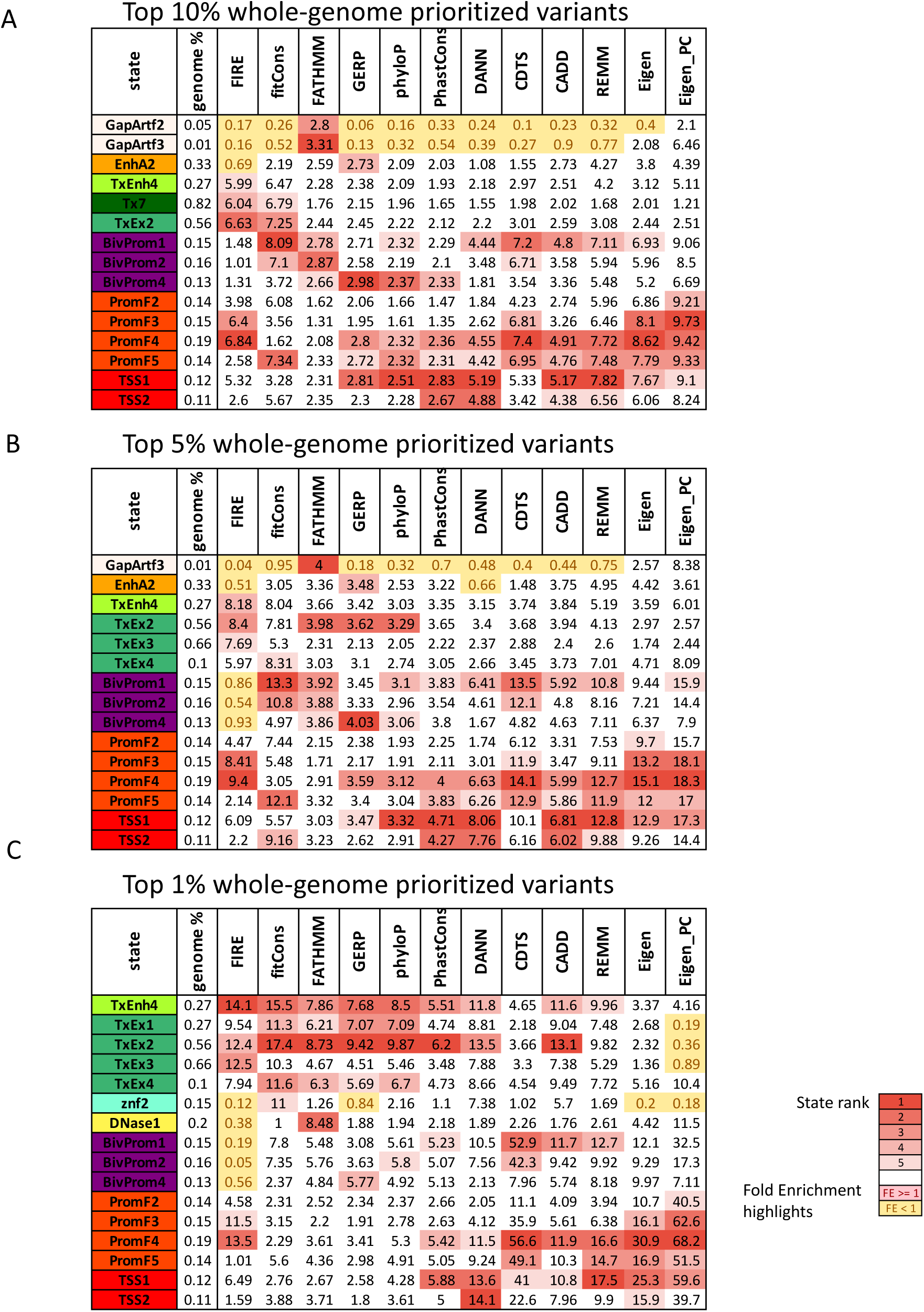
Full-stack states enrichment with bases prioritized by variant prioritization scores. A similar figure to **Supplementary Fig. 36**, except showing the top enriched states for prioritized variants from the whole genome not restricted to non-coding regions. Fold enrichment of full-stack states for genomic bases prioritized in the **(A)** top 10% **(B)** top 5%, and **(C)** top 1% bases by 12-different variant prioritization scores (**Methods**). Only states that were among the five with greatest enrichment for at least one score are shown. Top enrichment values are colored red based on the rank of the state for the score as indicated in the color legend at the bottom. Depletions are shown in yellow.

**Supplementary Figure 41:**
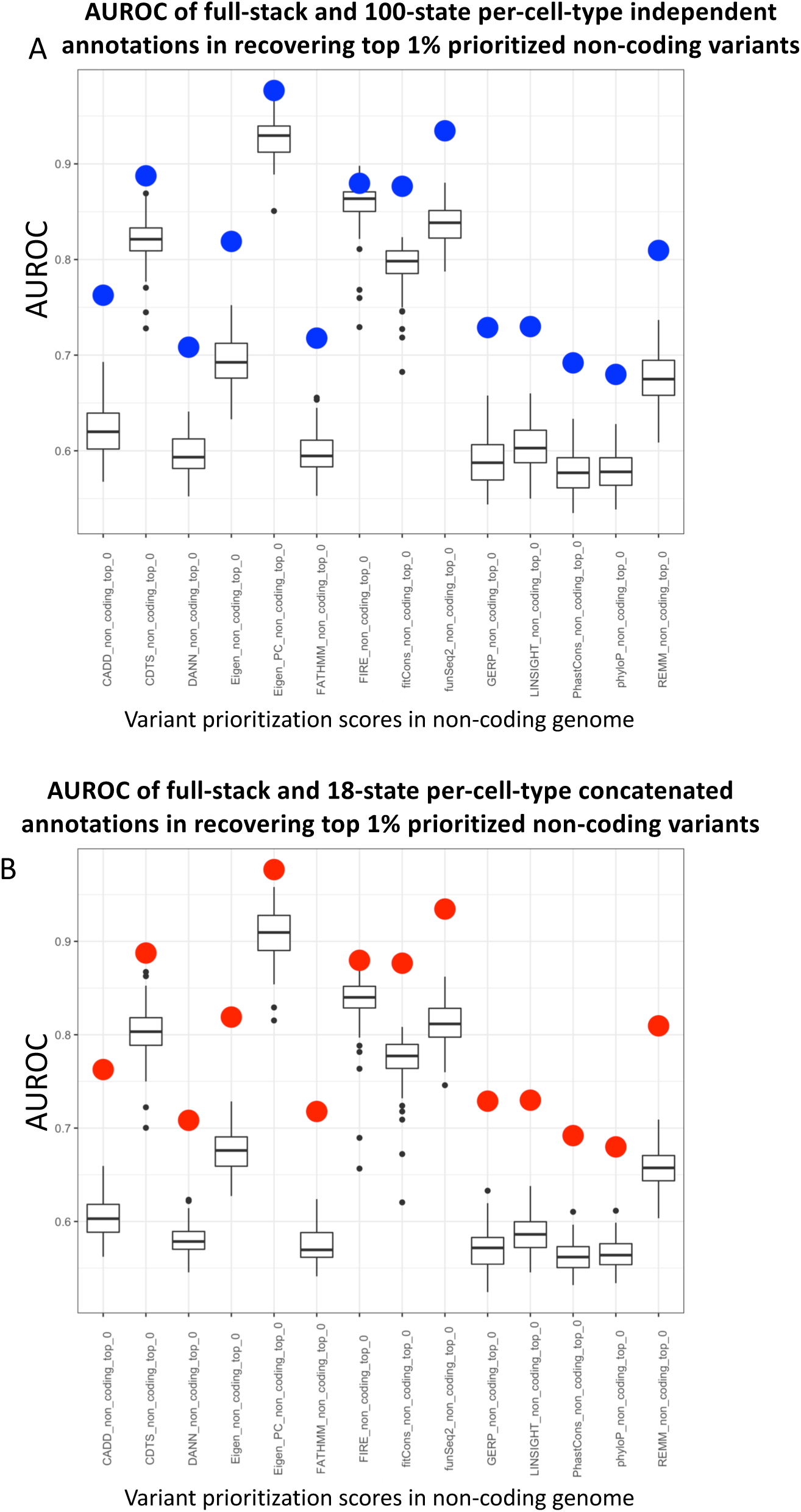
AUROC comparison of the full-stack model annotations and per-cell-type model annotations at predicting top 1% non-coding bases prioritized by various variant prioritization scores. The box-plots show AUROC of the **(A)** 127 100-state per-cell-type annotations from independent models and **(B)** 98 18-state per-cell-type annotations from concatenated models at predicting locations of the top 1% non-coding prioritized variants. In both panels, the x-axis represents different groups of top 1% non-coding bases prioritized variants previously curated in (Arneson & Ernst, 2019), based on 14-different variant-prioritization scores (**Methods**). The **(A)** blue and **(B)** red dots show the AUROC for the full-stack chromatin state annotations.

**Supplementary Figure 42:**
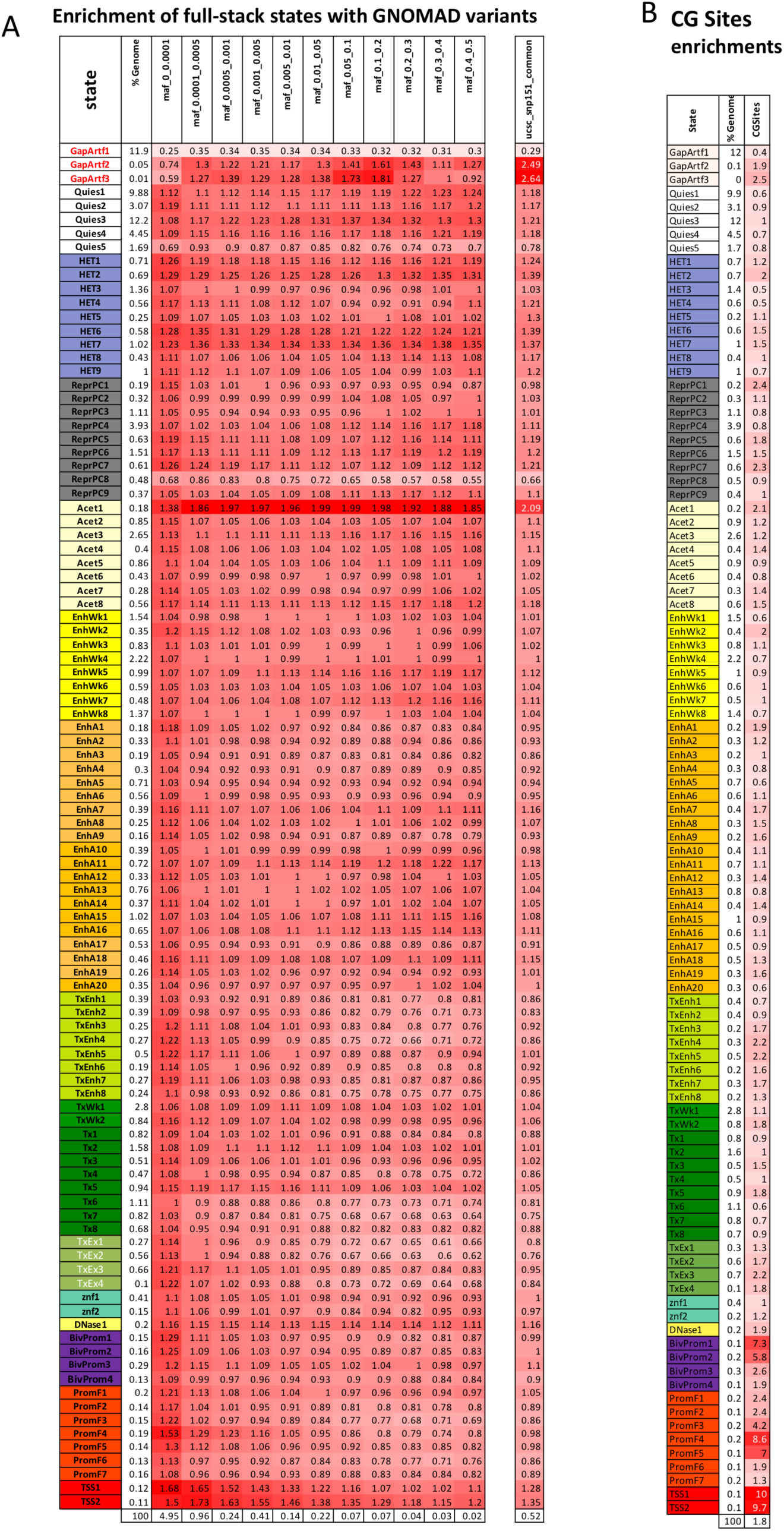
Full-stack states enrichments with variants from GNOMAD stratified by minor allele frequencies, common variants (A) and CG dinucleotides (B). In each subpanel, each row corresponds to a full-stack state. The first column gives the state labels, the second gives the percent of the genome that each state covers. The heatmap colors are on a column specific coloring scale. The last row shows the percentage of the genome that each group of variants occupy. **(A)** The last column shows the enrichment of full-stack states with common variants from UCSC Genome Browser’ snp151 track (Methods). Other columns show fold enrichments of full-stack states for GNOMAD variants with the specified ranges of MAF, which are ordered in increasing MAF (Karczewski et al., 2020). **(B)** The last column shows fold enrichment of full-stack states with CG dinucleotides. The three states showing highest enrichment if variants of lowest MAF (0 < MAF <= 0.0001) (TSS1-2, PromF4) are also the states most enriched states with CG dinucleotide sites, likely reflecting the higher mutation rates associated with CG dinucleotide sites (Karczewski et al., 2020).

**Supplementary Figure 43:**
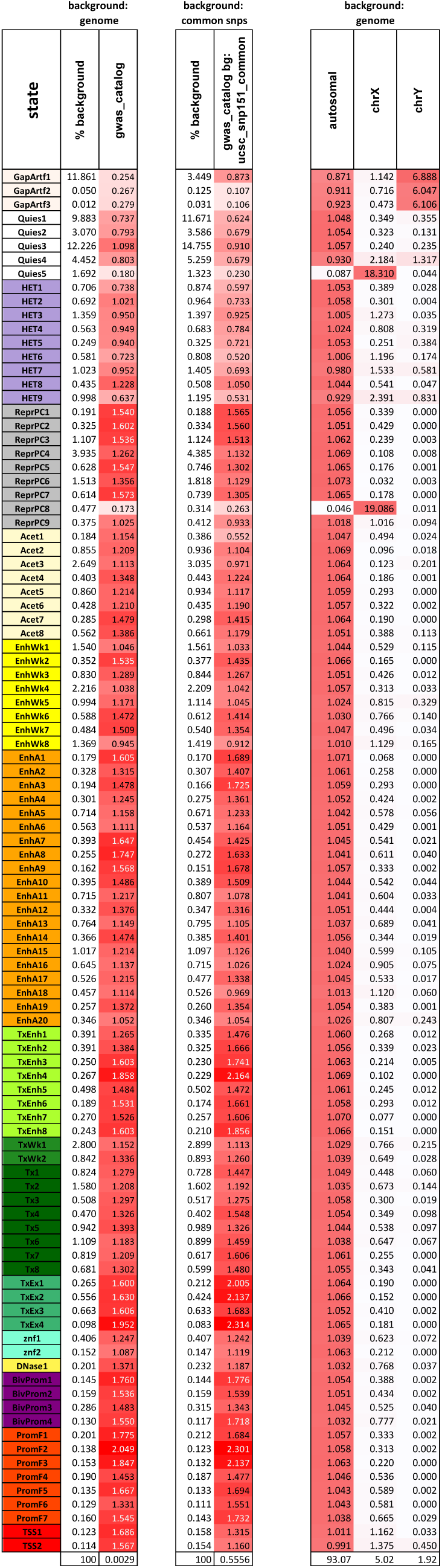
Full-stack states enrichments with GWAS catalog variants (Welter et al., 2014) and sex chromosomes. Each row corresponds to a full-stack state. The first column gives the state labels and the second column shows the percentage in the genome that each full-stack state occupies. **Th**e third column shows the fold enrichments of full-stack states with GWAS catalog variants against the whole-genome. The fourth column shows the percentage of the background context (UCSC snp151 common variants) that each full-stack state occupies. The fifth column shows fold enrichments with GWAS catalog variants against the background of common variants (**Methods**). The sixth, seventh and eight columns report the autosomal, chrX, and chrY enrichments, respectively. Columns are colored on a column specific coloring scale. The last row reports the percent of the background context (whole genome and set of common variants) that the GWAS catalog variants cover.

**Supplementary Figure 44:**
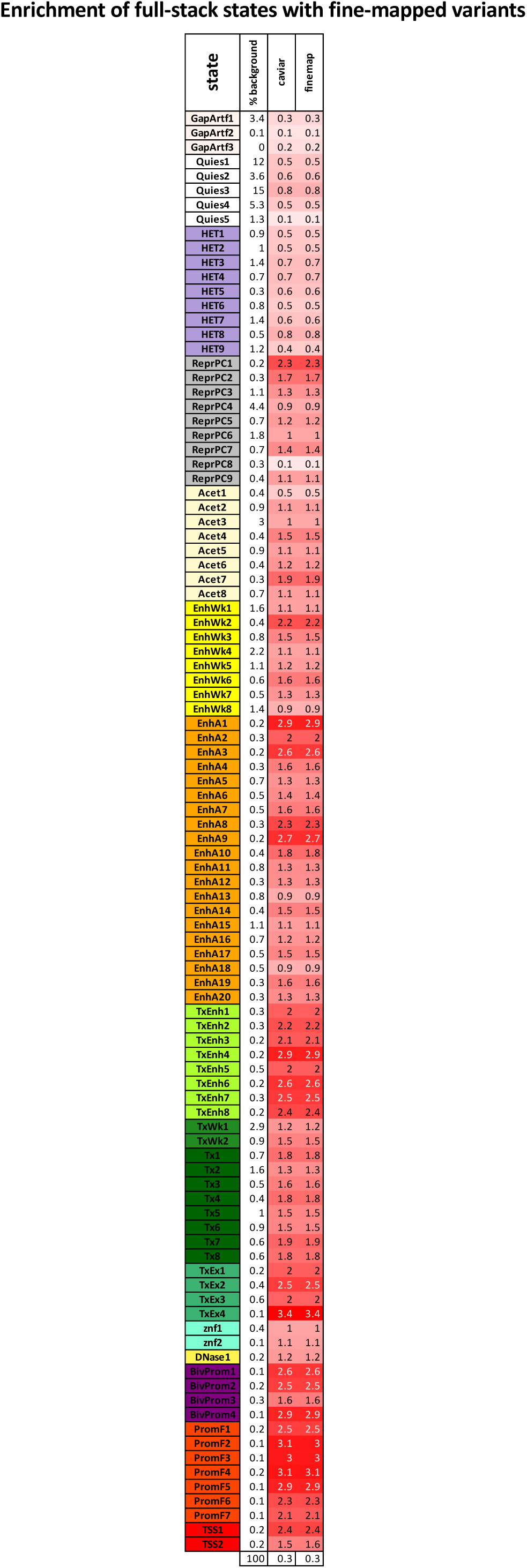
Full-stack states enrichment values for fine-mapped variants at phenotype associated loci. At phenotype associated loci, causal variants were fine-mapped by two methods, CAVIAR and Finemap (Benner et al., 2016; Tate et al., 2019). A set of lead fine-mapped variants in 1MB loci across the genome were identified (**Methods**). The figure shows the full-stack states’ enrichment values for these fine-mapped variants calculated against a background of common variants. The rows correspond to full-stack states. The first column gives the state labels, the second percent of the genome that each state covers, followed by columns with the fold enrichment for fine-mapped variants by CAVIAR and Finemap. The heatmap colors are on a column specific coloring scale. The last row shows the percentage of the background set of variants that the sets of lead fine-mapped variants occupy.

**Supplementary Figure 45:**
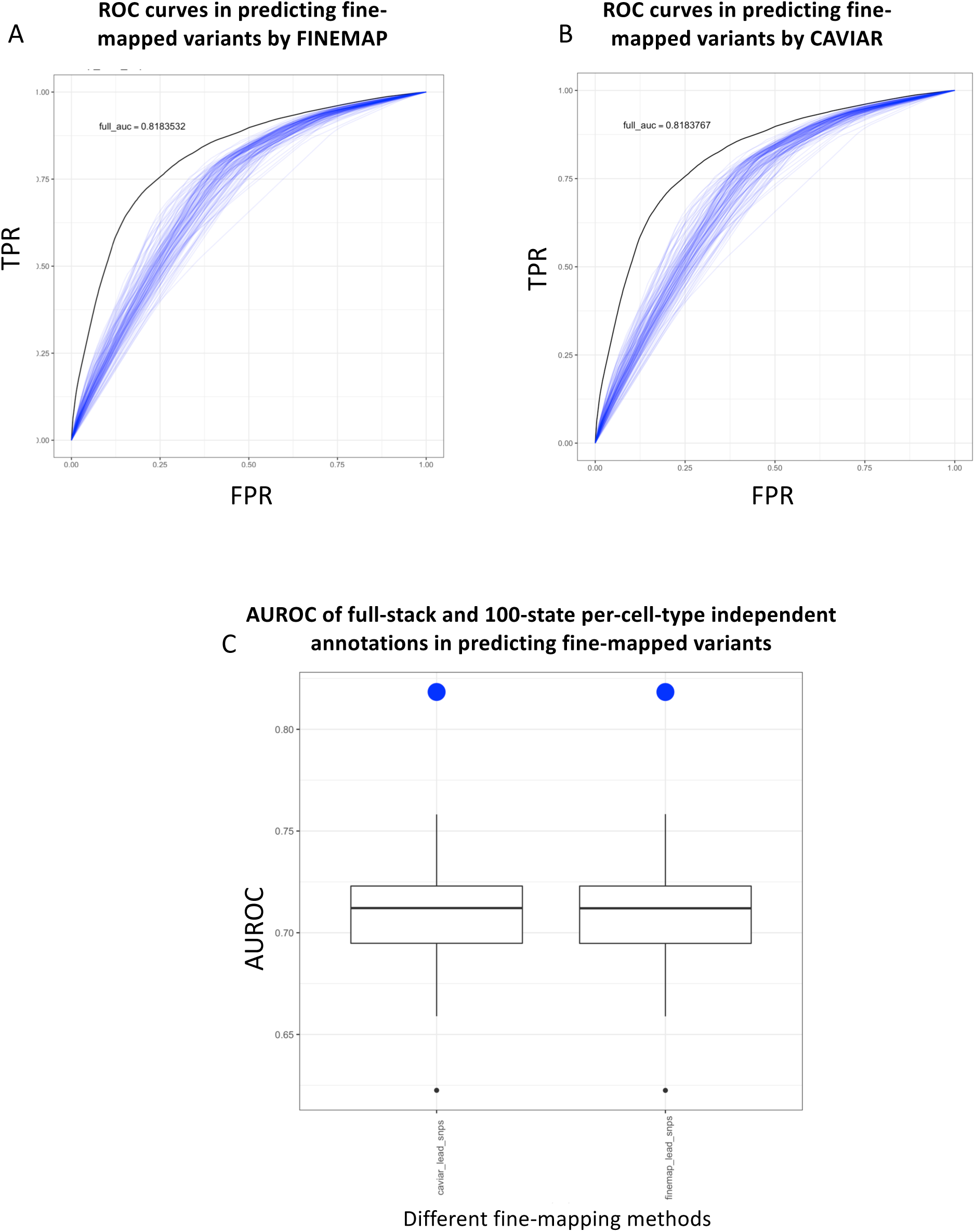
Comparison of full-stack model annotations and the 100-state per-cell-type annotations from independent models in predicting fine-mapped variants. **(A)** ROC curves for the full-stack model and the 127 100-state per-cell-type-independent models’ chromatin state annotations at predicting variants that show highest probabilities of being causal according to fine-mapping method FINEMAP (Benner et al., 2016) against the background of common variants (**Methods**). The full-stack annotation model’s ROC curve is in black and the 127 100-state per-cell-type annotations from independent models’ ROCs are shown in blue. **(B)** Similar plot as (A), but for variants evaluated by fine-mapping method CAVIAR (Chen et al., 2015). **(C)** Comparison of the AUROC in predicting fine-mapped variants from a background of common variants. The x-axis represents two different fine-mapping methods used to evaluate variants’ potential for causing diseases. The box-plots show AUROC of 127 100 per-cell-type-annotations from independent models in predicting these variants. The blue dots show the AUROC of the full-stack chromatin state annotations.

**Supplementary Figure 46:**
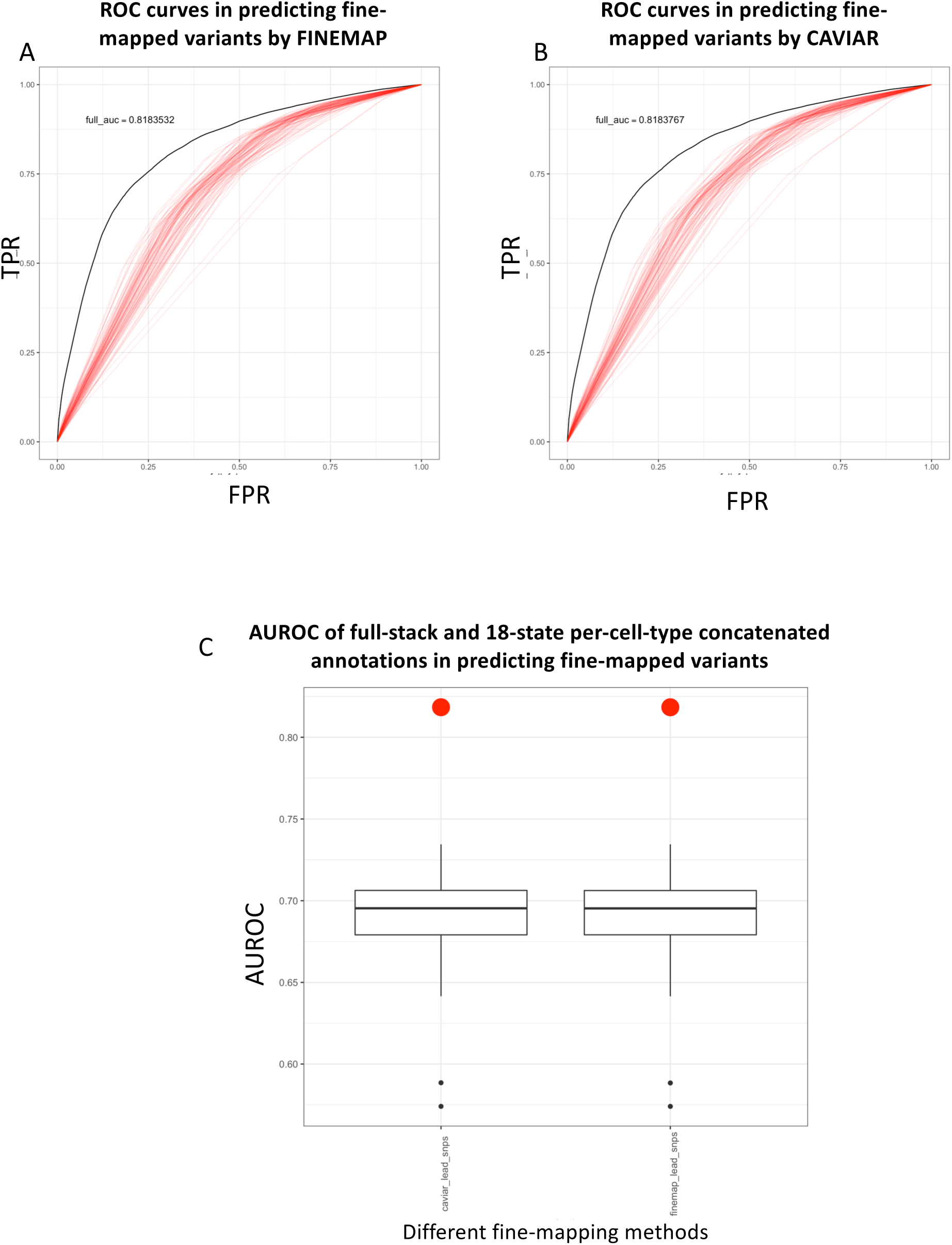
Comparison of full-stack model annotations and the 18-state per-cell-type annotations from a concatenated model in predicting fine-mapped variants. **(A)** ROC curves for the full-stack model and the 98 18-state per-cell-type annotations from concatenated models’ chromatin state annotations at predicting variants that show highest probabilities of being causal according to fine-mapping method FINEMAP (Benner et al., 2016) against the background of common variants (**Methods**). The full-stack model annotation’s ROC curve is in black and the 98 18-state per-cell-type annotations from a concatenated model ROCs are shown in red. **(B)** Similar plot as (A), but for variants evaluated by fine-mapping method CAVIAR (Chen et al., 2015). **(C)** Comparison of the AUROC in predicting fine-mapped variants from a background of common variants. The x-axis represents two different fine-mapping methods used to evaluate variants’ potential for causing diseases. The box-plots show AUROC of 98 18-state per-cell-type annotations from concatenated models in predicting these variants. The red dots show the AUROC of the full-stack chromatin state annotations.

**Supplementary Figure 47:**
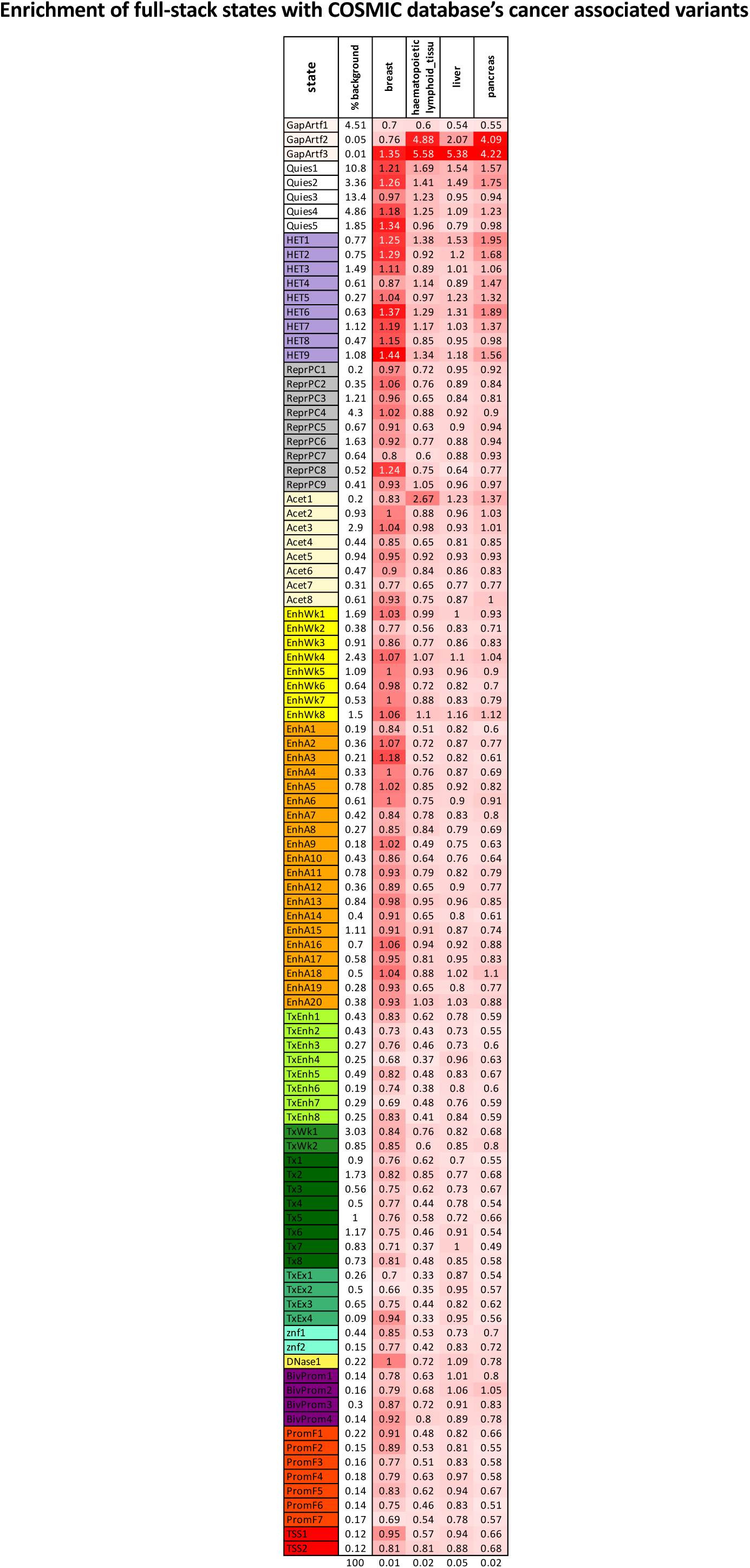
Full-stack states enrichments with cancer-associated somatic mutations in the non-coding genome. Each row corresponds to a full-stack state. The first column gives the state labels and the second column shows the percentage in the background genome context that each full-stack state occupies. For this analysis, the background context is the non-coding genome (**Methods**). The following columns correspond to one of four cancer types with the most number of mutations in the COSMIC database (Tate et al., 2019). These columns give the enrichments of full-stack states for mutations that appear at least once in the database for the cancer types. The heatmap colors are on a column specific coloring scale. The last row shows the percentage of the genome that mutations associated with each cancer type occupy.

**Supplementary Figure 48:**
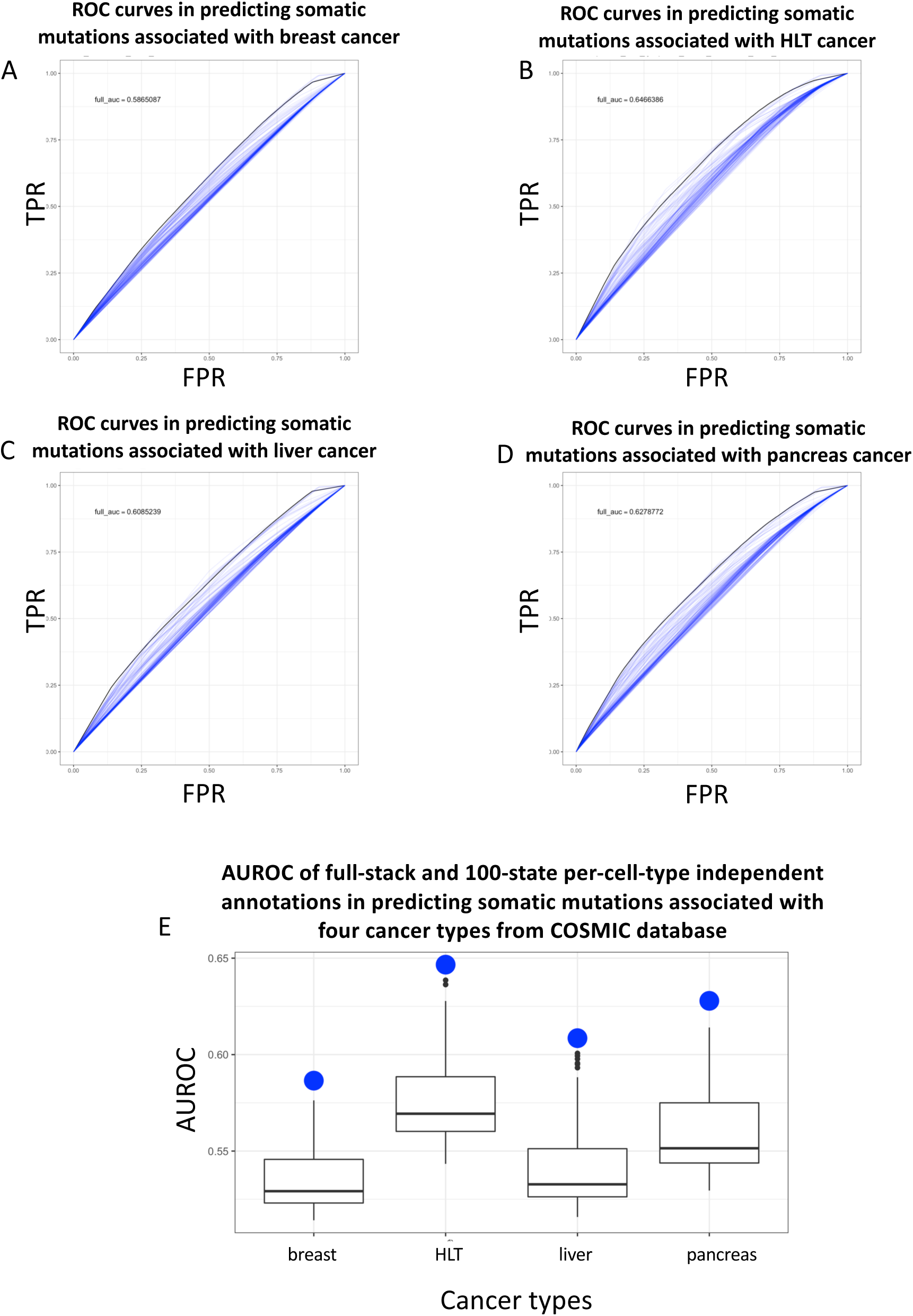
Comparison of full-stack model annotations and the 100-state per-cell-type annotations from independent models in predicting somatic mutations associated with four cancer types from COSMIC database (Tate et al., 2019). **(A)** ROC curves for the full-stack model’s annotations and the 127 100-state per-cell-type annotations from independent models at predicting somatic mutations associated with breast cancer against the background of non-coding genome (**Methods**). The full-stack model annotation’s ROC curve is in black and the 127 100-state per-cell-type-independent model annotations’ ROCs are shown in blue. **(B-D)** Similar plot as (A), but for mutations associated with **(B)** haematopoietic and lymphoid tissue (HLT) cancer, **(C)** liver cancer and **(D)** pancreas cancer, respectively. **(E)** Comparison of the AUROC in predicting cancer-associated somatic mutations from a background of non-coding genome. The x-axis represents four different cancer types that we considered in this analysis. The box-plots show AUROC of 127 100-state per-cell-type-independent models’ in predicting these mutations. The blue dots show the AUROC of the full-stack chromatin state annotations.

**Supplementary Figure 49:**
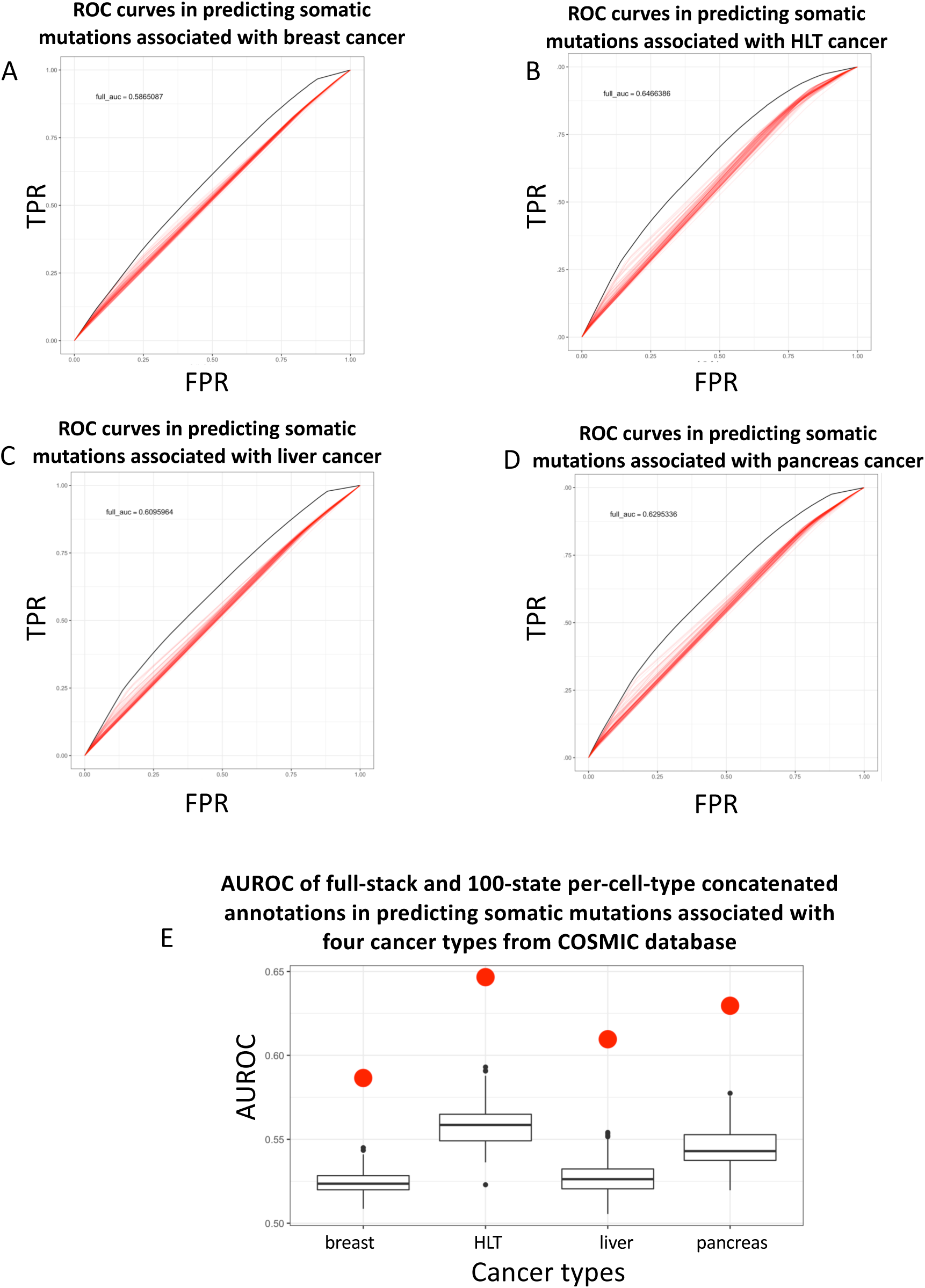
Comparison of full-stack states and the 18-state per-cell-type-concatenated chromatin state in predicting somatic mutations associated with four cancer types from COSMIC database (Tate et al., 2019). **(A)** ROC curves for the full-stack model and the 98 18-state per-cell-type-concatenated models’ chromatin state annotations at predicting somatic mutations associated with breast cancer against the background of non-coding genome (**Methods**). The full-stack model’s ROC curve is in black and the 98 18-state per-cell-type annotations from a concatenated model ROCs are shown in blue. **(B-D)** Similar plot as (A), but for mutations associated with haematopoietic and lymphoid tissue (HLT) cancer, liver cancer and pancreas cancer, respectively. **(C)** Comparison of the AUROC in predicting cancer-associated somatic mutations from a background of non-coding genome. The x-axis represents four different cancer types that we considered in this analysis. The box-plots show AUROC of 98 18-state per-cell-type annotations from a concatenated model in predicting these mutations. The blue dots show the AUROC of the full-stack chromatin state annotations.

**Supplementary Figure 50:**
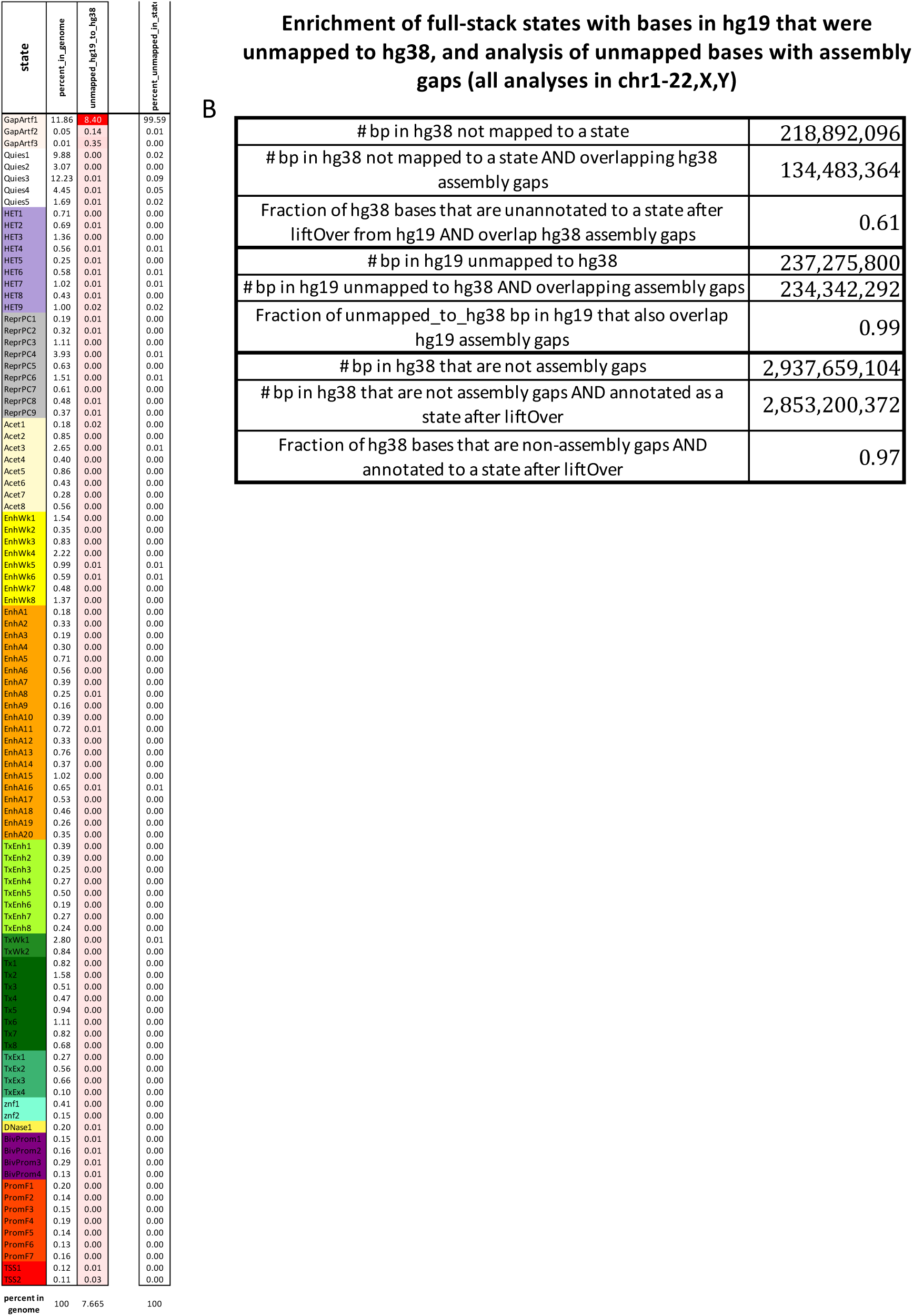
Full-stack states enrichments of bases that were not lifted over from hg19 to hg38. **(A)** The heatmap shows enrichment values for the full-stack states (rows) of genomic bases that were unmapped when lifting the state annotation from hg19 to hg38 (Methods). The first column shows the state label and the second column shows the percentage of the genome that each state covers. The third column shows enrichment values, colored such that highest enrichment values are colored red and lowest ones are colored white. The fourth column shows the percentage of the unmapped regions (from hg19 to hg38) in each state. **(B)** Table showing details of numbers of bases involved in liftOver procedure, highlighting the overlap between the unmapped and unannotated regions with assembly gaps. As part of the liftOver procedure, bases in hg38 that are mapped to from multiple bases in hg19 are left unannotated to any state in hg38 (Methods). All results are reported in chromosomes 1-22, X, Y.

**Supplementary Figure 51:**
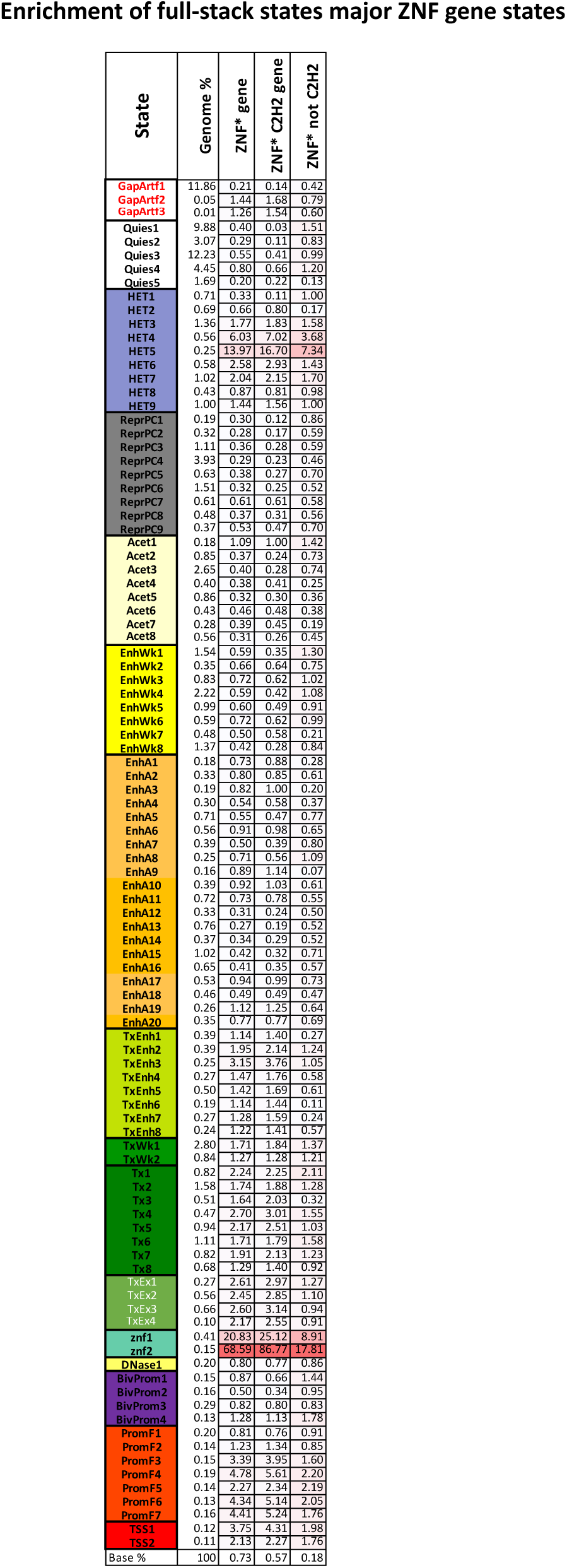
Full-stack states enrichments with different subsets of ZNF genes. The rows correspond to full-stack states. The first column presents the state labels, the second presents the percentage of the genome that each state occupies, and the remaining three columns enrichments for different subsets of zinc finger genes. The first of these is all genes with a ZNF symbol. The second is the subset of ZNF genes also annotated as C2H2 genes and the third those that are not C2H2 genes. The values correspond to the full-stack states’ fold enrichment for the ZNF gene families. Values are colored on a column-specific color scale. The last row gives the percentage of the genome that each type of ZNF gene family occupies

