## Supplementary information for "Universal annotation of the human genome through integration of over a thousand epigenomic datasets"

**Naïve-Bayes greedy search for representative experiments.**

We denote a subset of experiments whose signals can help us recover the full-stack state segmentations as the representative set $M^{*}$. We applied a greedy search of experiments to form the representative set $M^{*}$, i.e. at each iteration, we search for one experiment to add to the representative set $M^{*}$such that the newly modified set of experiments can best recover the original full-stack state segmentation. The initial representative set is empty. As re-computing chromatin state assignments for many different subsets of candidate experiments, *M*, based on ChromHMM’s normal procedure would not be computationally practical, we instead used an approximation of chromatin state assignments with a Naïve Bayes approach. First, using the full-stack state segmentation and the binarized genome-wide signals of 1032 experiments, we define the following properties:

- The state assignment at each 200bp segment, indexed $i$, along the genome is $S_{i}.$
- The prior probabilities of each state $s$ is estimated as:

$P(S_{i}= s) = \frac{\# positions where state s is assigned}{\# positions on the genome}$, for any position $i$

- The conditional probability of an experiment $m$being present ($1$) in each state $s$ is estimated as:

$P\left( m_{i}=1 \right| S_{i}=s)$ = $\frac{\# positions where mark m is present and state s is assigned}{\# positions where state s is assigned}$, for any position $i$.

Then, $P\left( m_{i}=0 \right| S_{i}=s)=1- P\left( m_{i}=1 \right| S_{i}=s)$.

Given we are using a candidate representative set of experiments $M$, At each genomic position $i$, the binarized signals of a potential representative set is denoted${\overset{⃑}{M}}_{i}$. We then calculate the posterior probabilities that this position is assigned to a particular state $s$ as:

$${P(S}_{i}=s \left| {\overset{⃑}{M}}_{i} \right)= P\left( S_{i}=s \right)\times P\left( {\overset{⃑}{M}}_{i} \right|S_{i}=s)=P\left( S_{i}=s \right)\times\prod_{m\in M} P\left( m_{i} \right|S_{i}=s)$$

At each genomic position $i$, the binarized signals of a potential representative set is denoted${\overset{⃑}{M}}_{i}$, and the chromatin state assigned here is denoted $S_{i}$. We then calculate the posterior probabilities that this position is assigned to a particular state $s$ as:

$${P(S}_{i}=s \left| {\overset{⃑}{M}}_{i} \right)= P\left( S_{i}=s \right)\times P\left( {\overset{⃑}{M}}_{i} \right|S_{i}=s)=P\left( S_{i}=s \right)\times\prod_{m\in M} P\left( m_{i} \right|S_{i}=s)$$

Position $i$ is assigned to the state with highest posterior probability, i.e. ${argmax}_{s}{P(S}_{i}=s \left| {\overset{⃑}{M}}_{i} \right)$.

The greedy algorithm we applied to obtained a representative subset of experiments $M^{*}$ is as follows:

- Step 0: The representative set $M^{*}$ at time 0 is empty: $M_{0}^{*}=\emptyset$.
- Step 1: For each of the remaining set of experiments, add each one to the current representative set separately, and denote each of those sets a candidate set of representative experiments $M$.
- Step 2: Apply Naïve Bayes framework to obtain genome-wide state assignment using the candidate representative set $M$, denoting such segmentation $S_{M}$.
- Step 3: Define the recovery score for each candidate set $M$ as:

$Re(M) = \sum_{s=1}^{100} \frac{\# positions where state s is assigned correctly in S_{M}}{\# postions where state s is assigned in full-stack segmentation}$

and choose the candidate set with the highest recovery score, i.e. ${argmax}_{M} Re(M)$.

Repeat steps 1-3 to add one experiment to the representative set $M^{*}$at each iterative.

**Asymptotic worst case time and memory usage of full-stack model**

Worst case compute time for each training iteration will grow linearly with the total length of the genome sampled per iteration. When the number of input tracks are sufficiently large, the worst-case compute time will grow linearly in the number of tracks. More specifically, let *M* be the number of input tracks (in our case, 1032), let *N* be the number of states (in our case, 100), let *L* be the length of the longest sequence (in our case, 5000, corresponding to one 1M-bp region of 200bp bins that we divided the genome into), let *S* be the number of sampled sequences per iteration (in our case, 300), and *p* the number of parallel processors (in our case, 6). For one iteration, the worst-case time on total compute usage is: $O(S(LN^{2}+M(min(L,2^{M})N+L)))$.

This is the case since before running the forward-backward algorithm, ChromHMM pre-computes the emission probabilities for each unique combination of observed marks and state in the sample sequences (in our case, the 300 regions randomly sampled at each iteration). Determining the combination of marks’ signals that each bin corresponds to requires a full pass on the sampled data, which is time $O(MSL).$ The number of observed combinations of marks per sequence is bounded by the smaller value between (1) the length of the sequence, *L*, and (2) the number of possible combinations with *M* inputs, which is $2^{M}$. The time to compute the emission probability for a single combination of marks for every state is $O(MN)$ and time $O(SMN* min(L,2^{M}))$ for every combination in each iteration (with $S$ sample sequences of maximum length $L$). With pre-computed emissions, running the forward and backward algorithm on a sequence of length *L* takes $O(LN^{2})$ time. This would need to be done on each of the *S* sampled sequences per iteration, thus taking $O(SLN^{2})$ total. The update to the emission parameters after running the forward-backward algorithm on each sequence $S$ can also be done in $O(SMN* min(L,2^{M}))$ time. The default number of training iterations, which we used here is 200.

The worst case wall-time per training iteration is $O(((S(LN^{2} + M(min(L,2^{M})N+L)))/p+ N(M+N)S)$ assuming efficient parallelization. This is the case since the entire procedure is parallelized across *p* processors except for aggregating statistics on the $O(N^{2})$ transition and $O(NM)$ emission parameters from each of the $p$ processors, which takes $O(N(M+N)S)$ time.

The worst case memory usage of the procedure is $O(p(LN^{2} +Mmin(L,2^{M}))+ N(M+N)S)$  since for each of the $p$ sequences being actively processed, the amount of memory to store the input is bounded by $O(Mmin(L,2^{M})+L)$ and the forward-backward procedure needs $O(LN^{2})$ additional memory.  Additionally, the output for each of the *S* sequences is stored before they are combined requiring $O(N(M+N))$ storage for each sequence.
